# Subcortex visualization: A toolbox for custom data visualization in the subcortex and cerebellum

**DOI:** 10.64898/2026.01.23.699785

**Authors:** Annie G. Bryant

## Abstract

Recent years have seen an expanding repertoire of code-based tools for visualizing neuroimaging data, promoting reproducibility and interpretability of brain-mapping findings. However, most open-source visualization packages for the human brain are geared toward the cerebral cortex, and the comparatively fewer options for the subcortex and cerebellum are limited in scope (i.e., atlas support) and flexibility. We address this critical gap by introducing `subcortex_visualization`, an open-source package offered in both Python and R, that provides a unified and accessible framework for programmatically visualizing non-cortical region-level data across many popular atlases for the subcortex (including thalamic nuclei and the brainstem) and cerebellum. These two-dimensional vector-based visuals are inspired by the `ggseg` R package for cortical data, which implements standardized rendering conventions to facilitate comparison across atlases in a vectorized two-dimensional format. In addition to the vectorized versions of twelve atlases—to our knowledge, the largest collection of non-cortical atlases in a single vector-based brain visualization toolbox—we also provide a step-by-step tutorial for users to generate custom vector-based visualizations from any given brain segmentation, enabling flexible extension to new atlases and structures. Collectively, `subcortex_visualization` and the accompanying documentation support reproducible and interpretable visualization of region-level data below the cortical mantle.

## 1 Introduction

The effective visualization of data across brain structures is crucial for interpreting and communicating brain-mapping results, for modalities ranging from neuroimaging to transcriptomics to cytoarchitecture. Frequently used options include volumetric slices (e.g., from NIfTI files), surface-mesh rendering, or two-dimensional (2D) vector plots. As reviewed in Chopra et al. [1], many open-access tools now programmatically render reproducible brain figures across these formats, bolstering image quality and enabling interactive exploration. However, most tools designed for the human brain target the cerebral cortex and far fewer support subcortical or cerebellar structures, leaving the field without a consistent, accessible way to depict region-level statistics from non-cortical atlases. The limited availability of code-based data visualization tools for the subcortex and cerebellum reflects a historical ‘cortico-centric’ bias in cognitive and computational neuroscience [2], arising in part from physical difficulties in obtaining reliable signal from more deeply embedded structures relative to more superficial cortical areas [3–5]. However, in tandem with increasing focus on the subcortex and cerebellum in health and disease, imaging resolution continues to improve [6, 7], such that more studies can start to depict analytical results from the subcortex and cerebellum alongside those of the cortex [8–10].

This expanding emphasis on the subcortex and cerebellum has been accompanied by a proliferation of various techniques for visualizing diverse types of data in these brain systems. The garden variety of visualization techniques now includes 2D slices sampled across neuroimaging planes [11–16], three-dimensional (3D) mesh rendering with data mapping [17–24], and 3D scatter plots of voxels, vertices, or regional centroids [9, 25–28]. Research groups generally develop their own in-house pipelines for depicting results obtained using their preferred subcortical and/or cerebellar atlases, giving rise to a fragmented visualization landscape below the cortical mantle. This fragmentation poses two key challenges for the field: (1) the independent development of code and visualization pipelines for each new atlas, which can be time-consuming and technically challenging for researchers who may not have extensive experience with 3D visualization software or programming; and (2) a lack of standardization in how subcortical and cerebellar data are visualized across studies and atlases, which can hinder comparability and reproducibility of findings. Of note, 2D vector graphics offer unique advantages for visualizing region-level data across multiple atlases in a consistent and interpretable way, as they are not subject to lighting inhomogeneities that can challenge the interpretation of color-mapped data in 3D renderings [29]. Recent years have seen ad hoc approaches to create 2D vector graphic representations for clear visualization of empirical data in the thalamus, amygdala, and cerebellum [11, 30–36], but these efforts have not been standardized across atlases or research groups—further compounding the fragmented landscape of subcortical and cerebellar data visualization.

Our vision for a comprehensive non-cortical data visualization toolbox was heavily inspired by the `ggseg` package in R [37] (and extensions in Python^1^), which provides a user-friendly interface for projecting empirical (region-level) data onto a variety of cortical atlases in two dimensions. The `ggseg` package and its extensions have collectively unified more than fifteen cortical parcellation atlases for programmatic data visualization in publication-ready vector graphics, and this software suite has been widely adopted across the field for visualizing region-aggregated cortical data in a consistent and interpretable way. While `ggseg` offers subcortical plotting with the FreeSurfer aseg atlas [38], it is not currently possible to show data from all seven subcortical regions (accumbens, amygdala, caudate, hippocampus, pallidum, putamen, thalamus) in one plot, as no single coronal or sagittal volumetric slice exists that includes all regions simultaneously. Users can project parcellated subcortical data onto 3D volumetric renderings in `ggseg` [37] as well as other packages such as the ENIGMA toolbox [39] (in Python/MATLAB), though interpretation of the color-mapped data is challenged by occlusion and lighting inhomogeneities [29]. Another key challenge is the limited atlas support beyond that of aseg, despite the increasing availability of alternative atlases for the subcortex [8, 40–42], including thalamic nuclei [43–45] and the brainstem [46, 47], as well as the cerebellum [48–50].

This tutorial and accompanying code repository address these critical gaps in two ways. First, we introduce a package implemented in both Python (`subcortex_visualization`) and R (`subcortexVisualizationR`) that collectively supports vectorized visualization of empirical (region-level) data across twelve popular parcellation atlases for the subcortex and cerebellum. To our knowledge, this is the largest compilation of subcortical (including thalamic and brainstem nuclei) and cerebellar atlases for vector-based data visualization in one unified toolbox with standardized rendering conventions. We also demonstrate how to apply the included atlases in our toolbox to flexibly summarize regional statistics from user-supplied empirical data in volumetric format, using neurotransmitter receptor and transporter positron emission tomography (PET) maps shared by Hansen and colleagues [19, 51, 52]. Beyond visualization for a single brain system, we show how this package can easily be combined with the `ggseg` package [37] to unify cortical, subcortical, and cerebellar visualizations in a consistent vectorized figure for comprehensive brain mapping. Second, we offer a step-by-step guide for adding vectorized representations for new atlases to the package that can readily interface with the visualization functionality of our Python and R packages. While the pre-rendered atlases included in our package represent a convenient starting point for visualizing data in the subcortex and cerebellum, we recognize that many users may wish to visualize region-level data in other atlases or parcellation schemes. We describe two alternative approaches that both implement openly available tools for rendering 3D meshes and converting them to reusable 2D vector graphic scaffolds, with recommendations depending on a user’s preference for prioritizing scalable automation or manual fine-tuning precision, respectively. These custom vector generation approaches are broadly applicable across both cortical and non-cortical structures; however, given the wide variety of existing approaches tailored to the cortex [1], we focus here on the subcortex and cerebellum to demonstrate the potential for visualizing data in these brain systems.

Our goal in providing both the twelve pre-rendered atlases and instructions for generating custom visualizations is to offer a flexible, accessible, and reproducible workflow for visualizing data across a wide variety of subcortical and cerebellar atlases, with the option to easily extend to new atlases as they are developed. The package allows users to map their own region-level empirical data onto any of the included atlases (or any custom atlas they may add) with a single function call in Python or R. Importantly, the resulting figures are resolution-independent vector graphics that are publication-ready, and can be further customized (e.g., color scheme, font size, etc.) as desired using standard vector graphic editing software, such as Inkscape or Adobe Illustrator. Altogether, we hope that this tutorial and accompanying code repository will provide a crucial resource for programmatically visualizing data below the cortical mantle in a clear and consistent way, supporting reproducibility and interpretability of brain-mapping findings across the whole brain.

## 2 Introducing the subcortex_visualization package

In the first half of this paper, we introduce the `subcortex_visualization` package (and its R counterpart, `subcortexVisualizationR`), demonstrating its functionality for code-based data visualization using twelve pre-defined subcortical and cerebellar atlases. We first discuss installation and then review the twelve atlases that are pre-loaded as vectorized images in the package before introducing simple code snippets that can be used to visualize empirical (region-level) data in a given parcellation atlas. Additional documentation for the toolbox is provided on the website hosted by the GitHub repository^2^. Finally, we show how users can apply any of the included atlases to extract regional statistics from volumetric data, demonstrated with the example of neurotransmitter receptor and transporter PET maps curated by Hansen and colleagues [19, 51, 52].

### 2.1 Installation and troubleshooting

Both the Python and R versions of our package are fully open-source, and users are encouraged to share any issues or questions via the GitHub repository and to contribute via pull requests.

The Python version of our package (`subcortex_visualization`) can be installed via the PyPI repository [56] or cloned from GitHub [57], with more detailed installation instructions provided in the package website^3^. The most straightforward installation via PyPI can be accomplished as below:

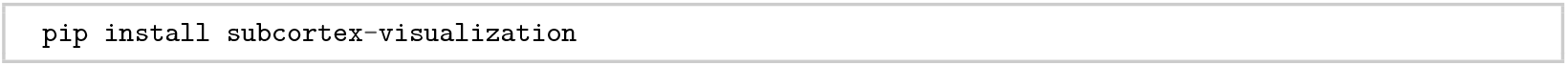

The R version of our package (`subcortexVisualizationR`) can be installed from the GitHub repository using the `remotes` package, as shown below (either in the terminal in an R session or from RStudio):

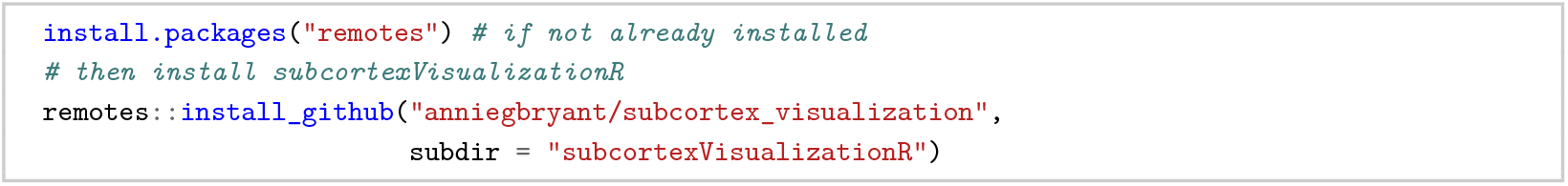

Alternatively, users can clone the repository and install the package from the local directory, which is necessary for users who wish to add new atlases to the package (see Sec. 3 for instructions on how to add a new atlas and integrate it with the existing package functionality). This also offers the advantage of more flexibility and customization of package functionality for more advanced users, as they can modify the source code directly. To clone the repository, users can run the following command in their terminal (e.g., in bash):

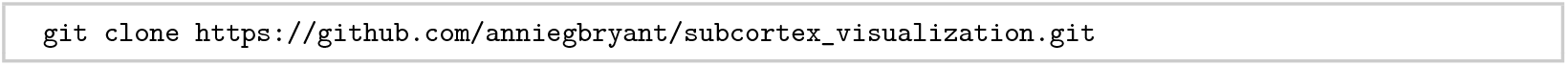

To install the Python package from the locally cloned repository, users can run the following from their terminal:

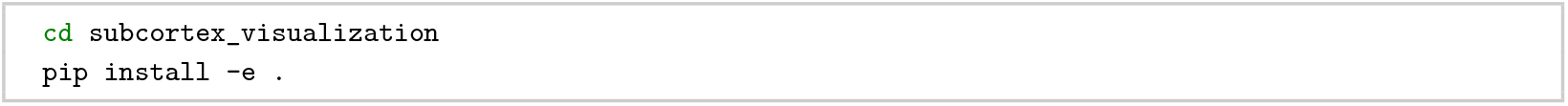

Similarly, for R installation from the locally cloned repository, users can run the following (in RStudio or an R session in the terminal):

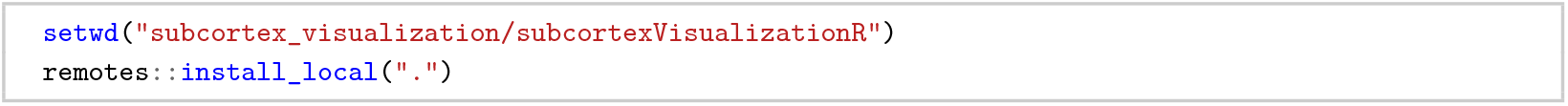

### 2.2 Use of artificial intelligence (AI) for software development

Generative AI (Claude, Anthropic) was used to assist in developing the `subcortex_visualization` toolbox in Python and R, specifically for code debugging and documentation. All suggested code snippets and documentation generated by the AI tool were reviewed and edited by the author to ensure accuracy and clarity.

### 2.3 Version control and package updates

The `subcortex_visualization` package is under active development, and we will continue to add new features and atlases over time (including those contributed by the community). In terms of Python and R versions, the package was developed with Python 3.11.15 and R 4.5.2, and we will continue working to maintain compatibility with future versions of both languages as they are released. We include library dependency information with pinned versions in the `requirements.txt` file for Python and the `renv` [58] package for R. More information about the package dependencies and how to update them can be found in the package website.

### 2.4 Viewing the atlases provided with the package

We have incorporated twelve widely used subcortical and cerebellar atlases as a starting point for data visualization—each with its own unique parcellation scheme and resolution (i.e., number of regions). These atlases are summarized in Table S3 and visualized in Fig. 1. Briefly, the twelve atlases for which we provide pre-vectorized images in this toolbox include the aseg subcortical atlas [38]; Melbourne Subcortex Atlas, from resolutions S1 through S4 [8]; AICHA subcortical atlas [40]; Brainnetome subcortical atlas [41]; CIT168 reinforcement learning subcortical atlas [53]; Human Connectome Project-based thalamic nuclei atlas from Najdenovska et al. [43]; THOMAS thalamic nuclei atlas [44]; Brainstem Navigator atlas [46]; and the surface-based probabilistic cerebellar lobule atlas [54], adapted from the volumetric SUIT lobule atlas [48, 55].

**Figure 1.**
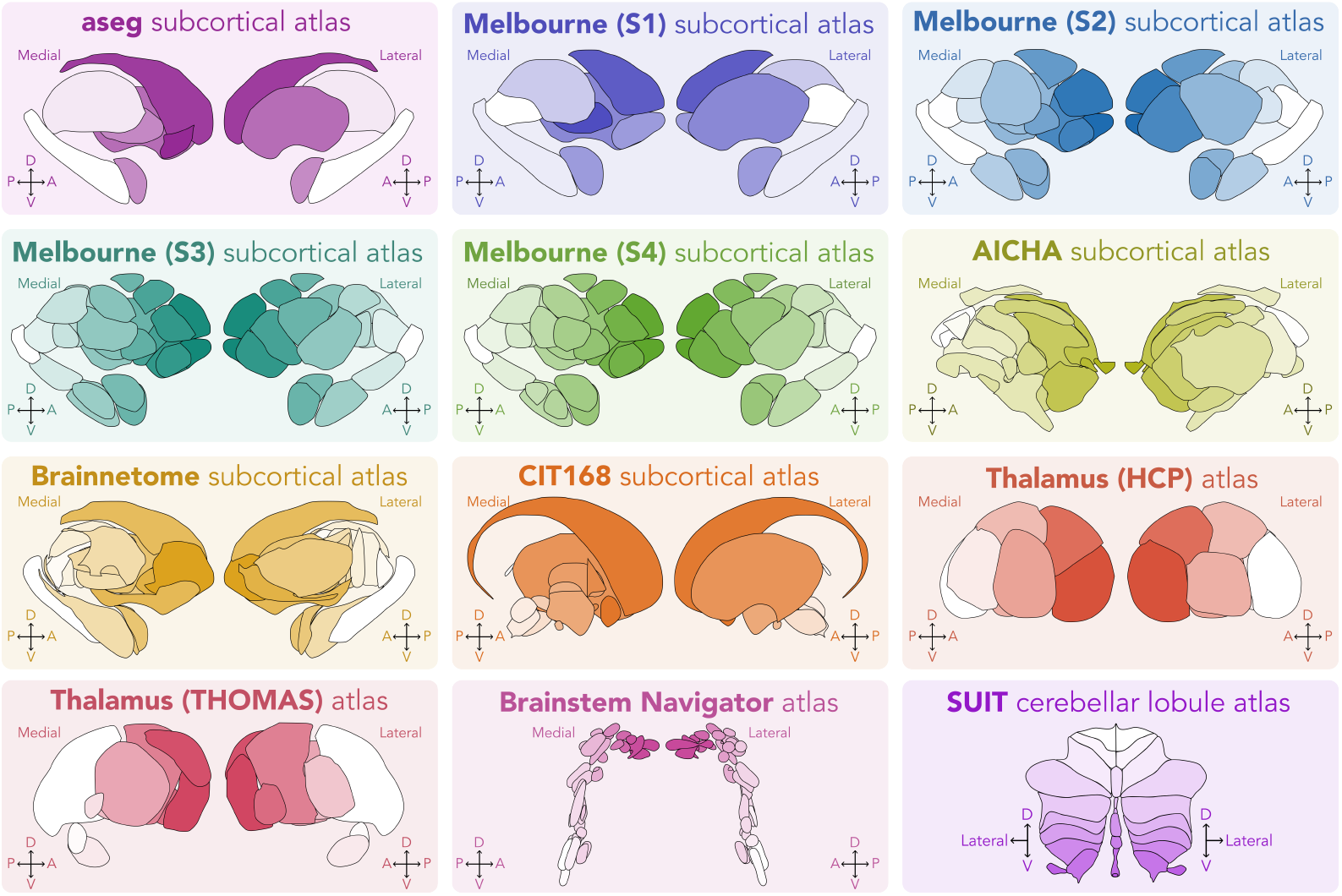
Twelve atlases are included in this package for ready-to-go visualization. These atlases include the aseg subcortical atlas [38]; Melbourne Subcortex Atlas, from resolutions S1 through S4 [8]; AICHA subcortical atlas [40]; Brainnetome subcortical atlas [41]; CIT168 reinforcement learning subcortical atlas [53]; HCP-based thalamic nuclei atlas from Najdenovska et al. [43]; THOMAS thalamic nuclei atlas [44]; Brainstem Navigator atlas [46]; and the surface-based probabilistic cerebellar lobule atlas [54], adapted from the volumetric SUIT lobule atlas [48, 55]. Atlases are shown for the left hemisphere, with the exception of the cerebellum (for which both hemispheres and the vermis are shown together). Orientation markers are included in each panel. D: dorsal; A: anterior; V: ventral; P: posterior. Each atlas is rendered with its own customized color scheme using the same plotting functionality as shown in Code snippets 3 (Python) and 4 (R).

#### 2.4.1 Introducing the plot_subcortical_data function

The central workhorse function of this package is `plot_subcortical_data`, which takes a variety of parameters designed to maximize flexibility in data visualization. If a user wishes to visualize empirical atlas-parcellated data in the form of region-aggregated statistics (e.g., the mean signal in each region of a given atlas), they can use this function to color-map the data (in the form of a `pandas DataFrame` in Python or `data.frame` in R) onto the regions of interest. Users can also specify a given atlas (any of the pre-vectorized twelve will work, along with any custom atlas that a user may incorporate), line color and thickness, transparency, and color map. For the Python package, users can specify any colormap available in `Matplotlib` (e.g., ‘viridis’ or ‘turbo’) or create a custom colormap using the `LinearSegmentedColormap` class in Matplotlib [59]; for the R package, users can specify any colormap available in the `viridis` package [60] or create a custom colormap using the `colorRampPalette` function in R (part of the base R installation). To maintain consistency across other visualizations, users can also set a fixed upper bound, lower bound, and/or midpoint for the colormap values. By default, the function will plot the index value for each region for the aseg atlas in the left hemisphere with black outlines and the *viridis* colormap. Users can also control the transparency of the plotted data through a global alpha parameter (`fill_alpha`), which is particularly useful to better resolve atlases with many overlapping nuclei. Additionally, in recognizing the benefits of visualizing statistical significance along with effect size [61], the function also allows users to specify a p-value column and a second alpha value for non-significant (i.e., *p >* 0.05) statistical results, such that regions with non-significant effects will be plotted with a user-defined level of transparency to visually de-emphasize them in the resulting figure.

In Fig. 2, we show the default output of the `plot_subcortical_data` function applied to the Melbourne Subcortex Atlas (S1 resolution), which colors the regions categorically using the *viridis* colormap. The syntax for this function is the same in both Python and R, as demonstrated in Code snippets 1 and 2, respectively (n.b., all code snippets are included in Appendix I). A similar figure can be easily generated for any of the other atlases by specifying the `atlas` keyword argument, as demonstrated in Code snippets 3 (Python) and 4 (R) with the AICHA subcortex atlas (plasma colormap) and Brainnetome subcortical atlas (rainbow colormap) in both hemispheres. The resulting brain maps are depicted in Fig. 3.

**Figure 2.**
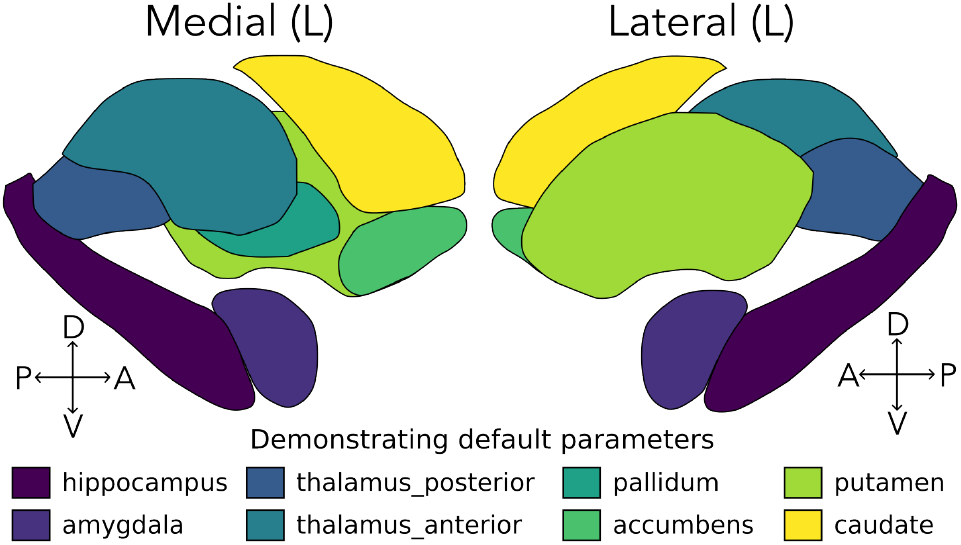
Example plot constructed using the default parameters of plot_subcortical_data. This figure was generated using the Melbourne Subcortex Atlas S1 parcellation in Code snippets 1 (Python) and 2 (R). Orientation markers were added to indicate the direction of each view. (L), left hemisphere; D, dorsal; V, ventral; A, anterior; P, posterior.

**Figure 3.**
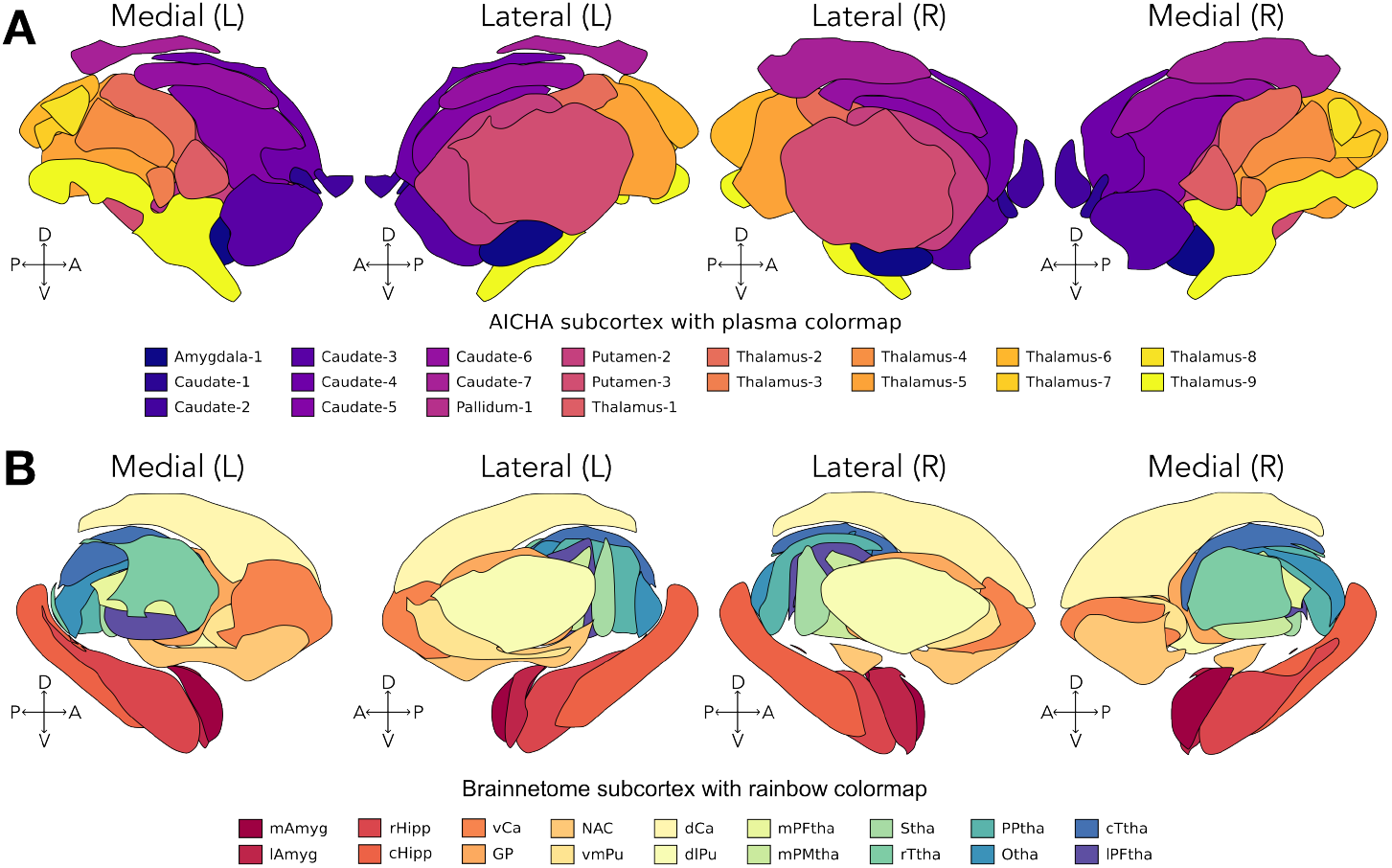
Example plot demonstrating the AICHA subcortical atlas and Brainnetome subcortical atlas in both hemispheres using different colormaps. **A**. AICHA subcortical atlas with plasma colormap. **B**. Brainnetome subcortical atlas with Spectral (rainbow) colormap. These figures were generated from Code snippets 3 (Python) and 4 (R). Orientation markers were added to indicate the direction of each view. (L), left hemisphere; (R), right hemisphere; D, dorsal; V, ventral; A, anterior; P, posterior.

#### 2.4.2 Choosing the right view(s) for your atlas

For each of the atlases included in this package, we include medial, lateral, superior, and inferior views for both hemispheres, either separately or combined, depending on the atlas. The only exception is the SUIT cerebellar lobule atlas, as this is based on a 2D flatmap representation of the cerebellum [54] that is not designed to be visualized in three dimensions. Users can specify which view(s) to plot with the `views` keyword argument in the `plot_subcortical_data` function, which accepts a list of views to plot (e.g., [‘`medial`’, ‘`lateral`’]). We demonstrate this functionality with two atlases: (1) THOMAS thalamic nuclei atlas, which includes separate medial and lateral views for each hemisphere, and (2) the Brainstem Navigator atlas, the latter of which includes combined left/right superior and inferior views as well as separate medial and lateral views for each hemisphere. The resulting brain maps for each atlas are depicted in Fig. 4A and Fig. 4B, respectively.

**Figure 4.**
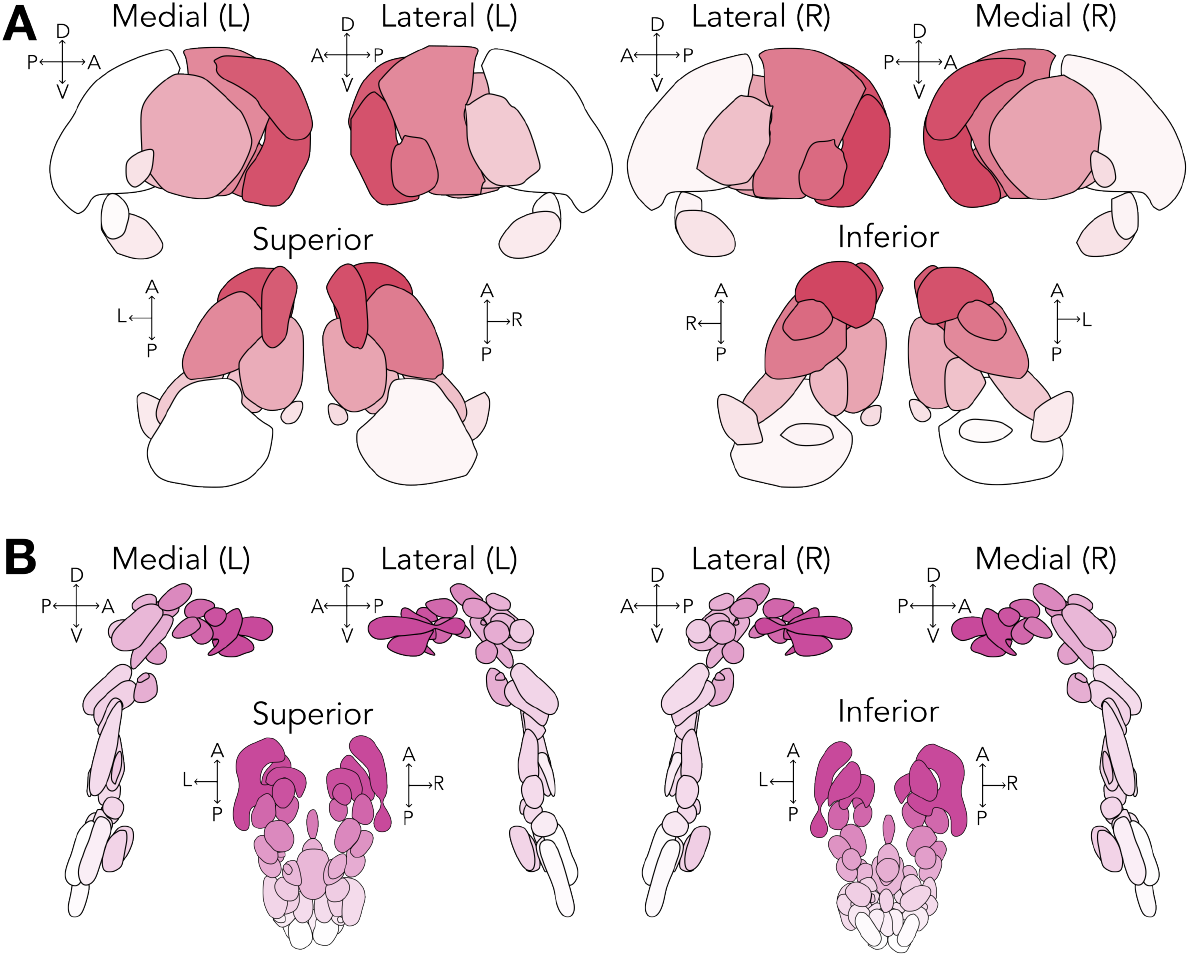
Four view angles are supported for plotting per hemisphere: Lateral, medial, superior, and inferior. **A**. The four view angles are depicted for the THOMAS Thalamus atlas. Orientation markers are used to indicate the direction of each view. (L), left hemisphere; (R), right hemisphere; D, dorsal; V, ventral; A, anterior; P, posterior. **B**. The four view angles are depicted for the Brainstem Navigator atlas. These atlases were visualized using Code snippets 5 (Python) and 6 (R).

### 2.5 Working with your own data

There are myriad types of empirical data from which region-level statistics can be meaningfully summarized and visualized, including neuroimaging data (e.g., functional MRI [62–64], diffusion MRI [65, 66], or PET [19]), bioinformatic data (e.g., gene expression [67, 68], protein abundance [69, 70], or neuropeptide density [71]), and cytoarchitectural data (e.g., cell density [72–74]). Given the heterogeneous nature of these potential underlying data sources, we encourage users of our package to compute region-level summary statistics of their choosing with the appropriate pipeline tailored to their data, so long as they use one of the twelve atlases we currently support (or one they have added to the library; cf. Sec. 3).

Regardless of the underlying data type or method for deriving regional summary statistics, the key requirement for visualizing data with the `plot_subcortical_data` function is that the resulting regional statistics are organized into tabular format (i.e., a `DataFrame` in Python or `data.frame` in R) with a column of region names that match those defined in the atlas exactly (including capitalization and spacing). The region names for a given atlas can be shown directly with the following code, using the Melbourne Subcortex Atlas with S1 resolution as an example (same syntax in R and Python):

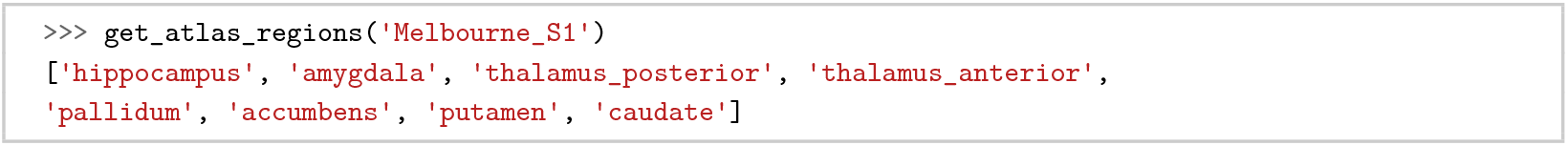

The region names for each atlas can also be found on the package website^4^ and are provided in tabular format for each atlas in the Supplementary Tables (Appendix III, Table S1).

#### 2.5.1 Plotting continuous data across atlas regions

To demonstrate this functionality, we simulate random continuous data to plot across the regions of the aseg subcortex atlas in Code snippets 7 (Python) and 8 (R). The first five rows of the resulting `DataFrame` are shown below as Table 2 to illustrate the required format for visualizing data with the `plot_subcortical_data` function, which includes a column of region names that match those defined in the atlas (e.g., ‘hippocampus’, ‘amygdala’, etc.), the hemisphere (e.g., ‘L’ or ‘R’) and a column of values to plot (in this case, ‘value’). This simulated data can be directly mapped onto the aseg subcortex atlas using the `plot_subcortical_data` function, as shown below and depicted in Fig. 5—with the corresponding code included in Code snippets 9 (Python) and 10 (R).

**Table 1.**
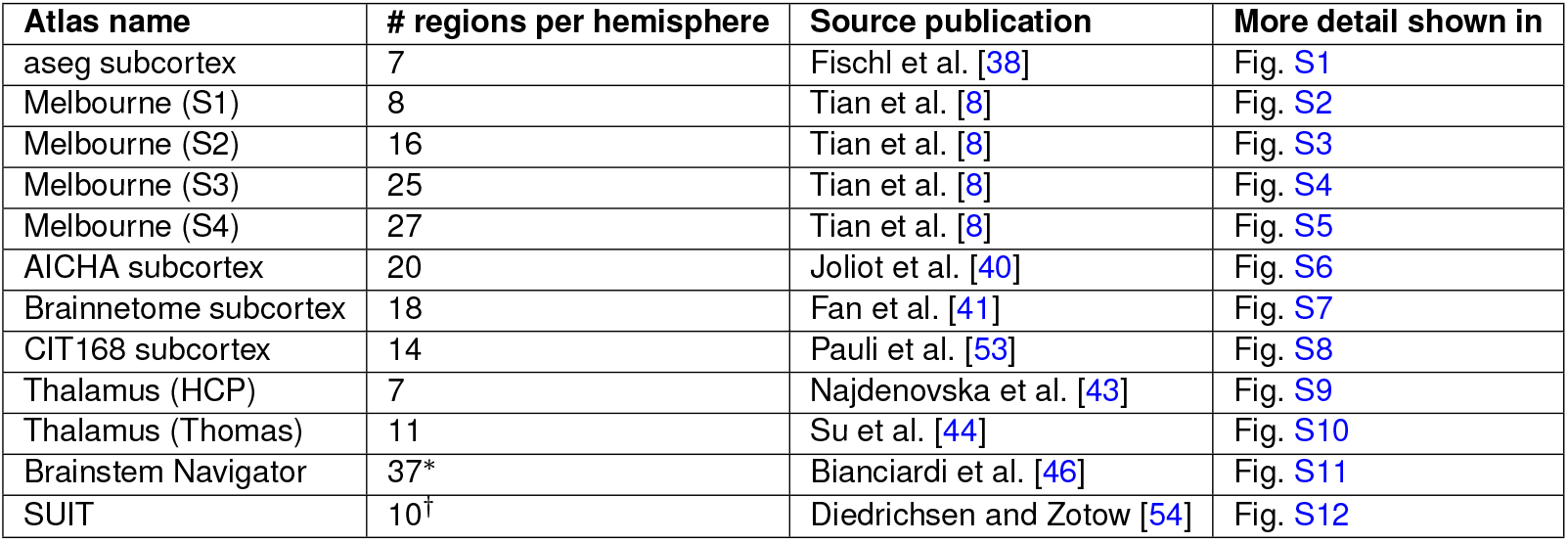
The twelve subcortical and cerebellar parcellation atlases provided in `subcortex_visualization` are summarized. For each atlas, we indicate the number of regions per hemisphere, the source publication and the figure detailing conversion from 3D mesh to 2D vector. ^∗^The Brainstem Navigator atlas contains 37 unique nuclei (regions) per hemisphere, though 8 of these are midline and thus shared between the two hemispheres, yielding 66 regions in total (29 left hemisphere, 29 right hemisphere, and 8 midline). ^*†*^The SUIT cerebellar lobule atlas contains 10 unique lobules per hemisphere, though the vermis is represented as a separate region for 7 of these lobules, yielding 27 regions in total (10 left hemisphere, 10 right hemisphere, and 7 vermis).

**Table 2.** An example Pandas DataFrame with simulated continuous data in the left hemisphere of the Melbourne Subcortex Atlas (S1 resolution).

| region | Hemisphere | value |
| --- | --- | --- |
| hippocampus | L | -0.5718095 |
| amygdala | L | 0.02962396 |
| thalamus_posterior | L | 0.56259194 |
| thalamus_anterior | L | -0.6476519 |
| pallidum | L | -0.8454361 |

**Figure 5.**
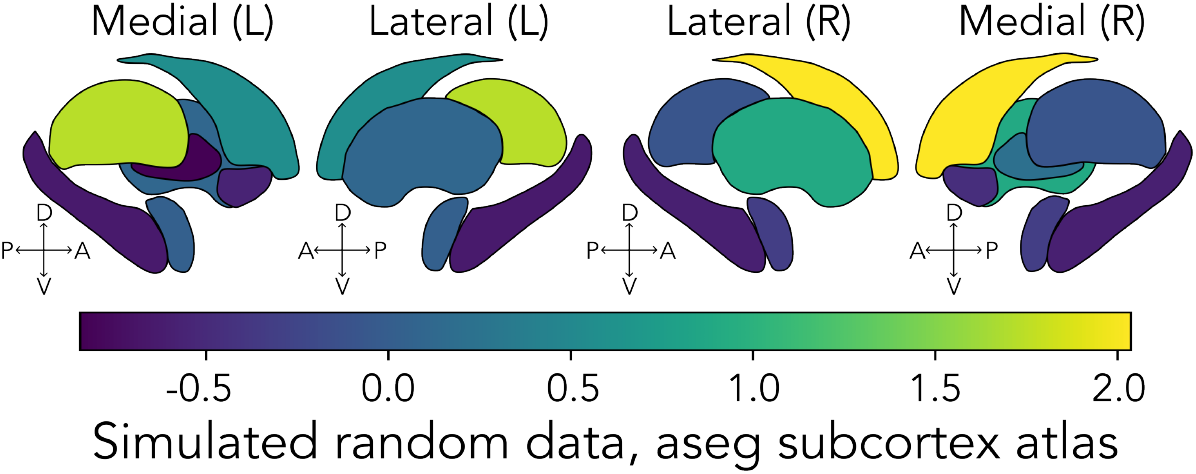
Simulated continuous data plotted in the `aseg_subcortex` atlas in both hemispheres. Orientation markers were added to indicate the direction of each view. (L), left hemisphere; (R), right hemisphere; D, dorsal; V, ventral; A, anterior; P, posterior. This figure was generated using Code snippets 9 (Python) and 10 (R), which use the default ‘viridis’ colormap.

One can also pass in a custom colormap to the `cmap` argument, as demonstrated in Code snippets 11 (Python) and 12 (R)—using a blue-white-red colormap to highlight both positive and negative values in the regional data (Fig. 6). `plot_subcortical_data` also takes an argument, `midpoint`, which specifies the center point for the color palette, as demonstrated in Code snippets 11 (Python) and 12 (R). Setting this to 0 here enforces the center value to be white. Without explicitly setting the upper and lower bounds (`vmax` and `vmin`, respectively), the color range will be defined symmetrically around `midpoint` to capture the full range of the data.

**Figure 6.**
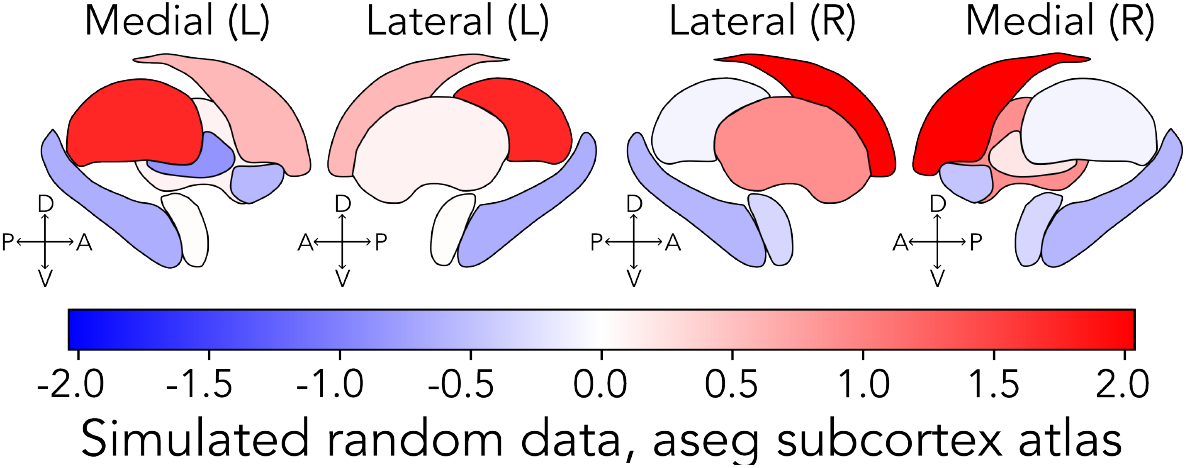
Simulated continuous data plotted in the `aseg_subcortex` atlas in both hemispheres, with a custom diverging color palette. Orientation markers were added to indicate the direction of each view. (L), left hemisphere; (R), right hemisphere; D, dorsal; V, ventral; A, anterior; P, posterior. This figure was generated using Code snippets 11 (Python) and 12 (R).

#### 2.5.2 Controlling transparency

As mentioned above, users can control the transparency of the plotted data through the global (i.e., all-region) transparency parameter `fill_alpha` in the `plot_subcortical_data` function, which is particularly useful to better resolve atlases with many overlapping nuclei. This is demonstrated in Code snippets 13 (Python) and 14 (R) with the Melbourne Subcortex Atlas (S2 resolution), in which the `fill_alpha` value ranges from 0.2 (high transparency) to 1 (no transparency). The resulting brain maps for each `fill_alpha` value are depicted in Fig. 7.

**Figure 7.**
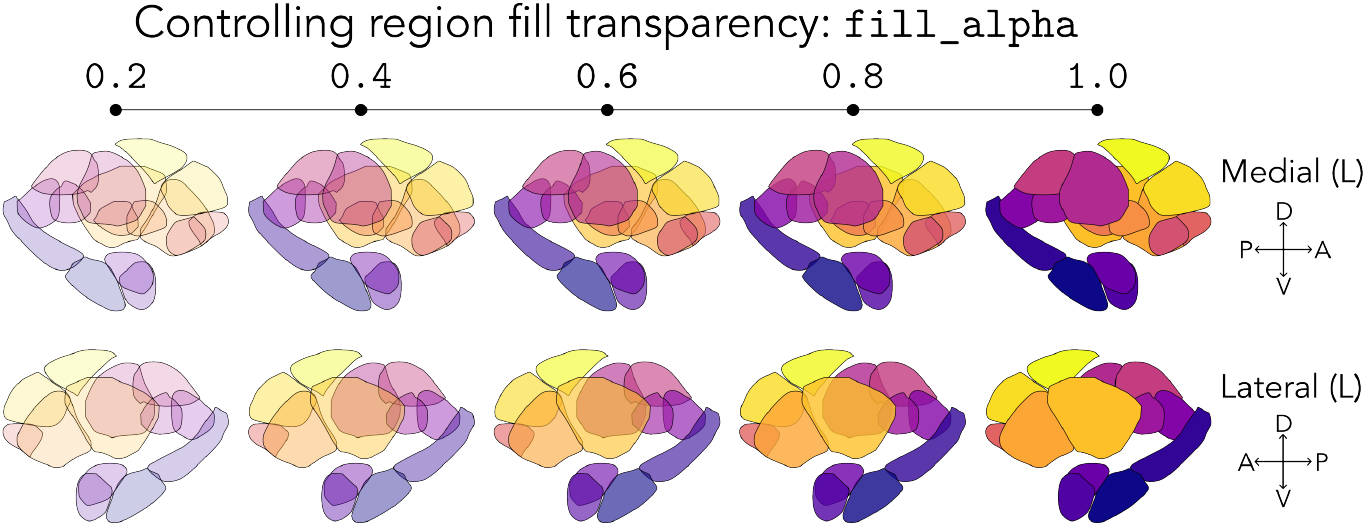
The transparency of region fill colors can be controlled with the fill_alpha parameter. This figure depicts the Melbourne Subcortex Atlas (S2 resolution) with the ‘plasma’ colormap, with each region labeled by its segmentation index, across five different values of the `fill_alpha` parameter, which controls the transparency of region fill colors. Specifically, the atlas is shown with `fill_alpha =` {0.2, 0.4, 0.6, 0.8, 1.0}. Orientation markers are used to indicate the direction of each view. (L), left hemisphere; D, dorsal; V, ventral; A, anterior; P, posterior. The figure was created using Code snippets 13 (Python) and 14 (R).

Alternatively, users may wish to visualize statistical significance along with effect size, for which transparency is particularly well suited as a visualization tool [61]. The `plot_subcortical_data` function accepts a `fill_by_significance` Boolean argument, which, when set to `True`, allows users to specify a p-value column in the `subcortex_data` DataFrame to dictate the transparency for each region (controlled by a separate `nonsig_fill_alpha` parameter, with a default value of 0.5). If *p* < 0.05 for a region, the alpha value for that region will be set to the value specified in `fill_alpha` (1.0 by default), and the line thickness for that region will be set to the value specified in `line_thickness` (1.5 by default in Python, 0.5 in R). By contrast, for a region with *p* ≥ 0.05, its alpha value will be `nonsig_fill_alpha` (0.5 by default) and its line thickness will be set to 0.25 *×* `line_thickness`; this effectively de-emphasizes non-significant regions in the resulting figure. An example of this functionality is depicted using the SUIT cerebellar lobule atlas in Fig. 8, in which we simulated random p-values for each region and used these to specify the transparency of each region in the resulting figure (as shown in Code snippets 15 for Python and 16 for R).

**Figure 8.**
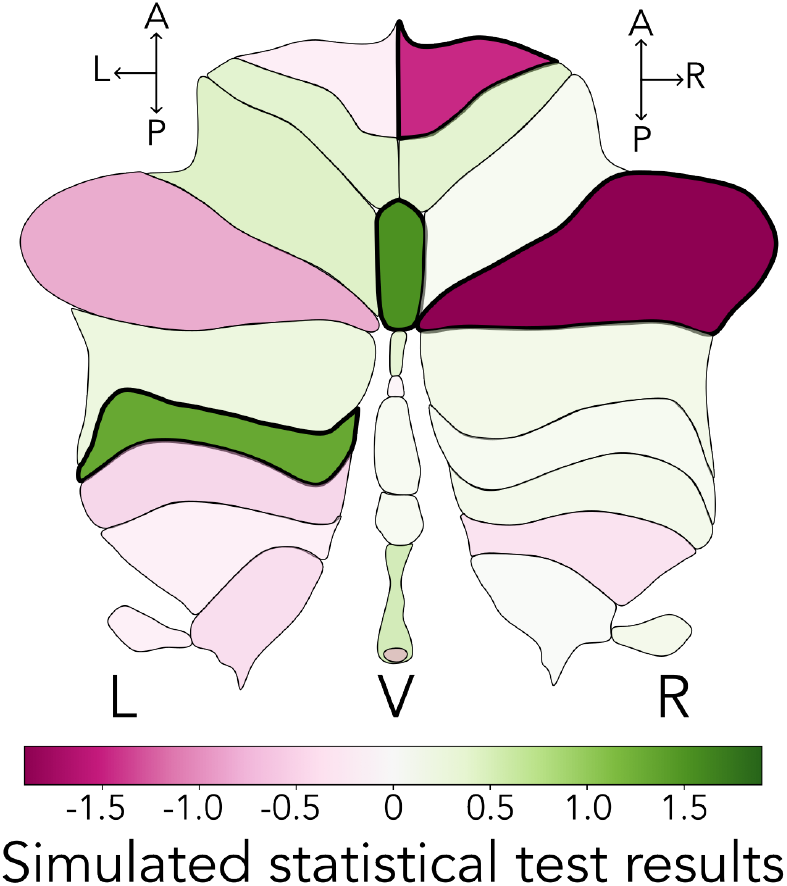
Statistical significance can be used to determine opacity and line thickness across regions. Here, we have simulated data and *p*-values in the SUIT Cerebellar Lobule atlas and set `fill_by_significance=True`. Regions with non-significant results (i.e., *p* ≥ 0.05) are shown with lower opacity (governed by the `nonsig_fill_alpha` parameter, here set to 0.5) and thinner lines (0.25*×* the line width of significant regions, the latter of which is governed by the `line_thickness` parameter). This plot was created using Code snippets 15 (Python) and 16 (R). (L), left hemisphere; R, right hemisphere; V, vermis.

### 2.6 Extracting regional statistics from volumetric data

In the case of a 3D volumetric image (from any imaging modality), there are many principled ways to reduce the dimensionality down from thousands of voxels to region-level aggregates, from basic central tendencies (e.g., mean or variance) to more complex dimensionality reduction methods or graph-theoretical network properties. If a user wishes to simply compute the average signal across all voxels per region in a given atlas, there are many suitable options that are straightforward to implement programmatically. For example, this includes the `Parcellator` object from the *neuromaps* package [52], the `fslmeants` function from FSL [75], or the `NiftiLabelsMasker` function from *nilearn* [76]. However, aside from the powerful *BrainSpace* toolbox by Vos de Wael et al. [77], the flexibility to compute statistics other the mean value across voxels has not yet been implemented in leading programmatic neuroimaging parcellation pipelines. In an effort to emulate the streamlined programmatic interface these packages offer to compute the voxel-averaged signal per region in a given atlas, we include the `parcel_segstats` function in `subcortex_visualization`, which takes as input a volumetric dataset (in NIfTI format) and an atlas name (any of the twelve included atlases, or a custom atlas that a user may incorporate). This function is inspired by the `reduce_by_labels` function in the *BrainSpace* toolbox [77], which allows users to summarize across all voxels within a given region of an atlas using any function they choose; our function is specifically tailored to handle the atlases included in our toolbox, to check for compatibility between the atlas and the user-supplied volumetric data, and to format the output in a way that directly interfaces with our visualization pipeline. For our implementation, users can specify any function that takes a one-dimensional array of values and returns a single summary statistic (e.g., `numpy.mean`, `numpy.std`, `numpy.median`, etc. [78]) for the `parc_stat` argument to compute the desired summary statistic for each region per hemisphere (or the vermis, in the case of the cerebellum). Background voxels are ignored by default with a value of `background_value` (0 by default), but this feature can be turned off (i.e., background voxels included) by setting `ignore_background=False`.

#### 2.6.1 Compatibility of input volumetric data with the included atlases

In order to compute regional statistics from empirical volumetric data in NIfTI format, the atlas(es) itself must be applied in volumetric format that aligns with that of the empirical data. The twelve subcortical and cerebellar atlases that we include in this package were originally shared in different alignment templates from the Montreal Neurological Institute (MNI) 152 space [79]. The specific template to which each atlas was originally aligned and shared is summarized in Table S2, which includes the MNI152NLin6Asym, MNI152NLin2009cAsym, MNI152NLin2009bAsym, and MNI152NLin2009cSym templates. We selected two commonly used templates, MNI152NLin6Asym and MNI152NLin2009cAsym, to offer two sets of our twelve atlases in a unified format. For all required space transformations (e.g., from MNI152NLin6Asym to MNI152NLin2009cAsym), we used the *TemplateFlow* [80] Python API. Some conversions already had pre-computed transformations included as part of the TemplateFlow archive; for others, we performed registration using the `registration` function from the ANTS toolbox in Python [81], using the `type_of_transform=`‘`SyN`’ function to specify symmetric normalization. In either case, the corresponding transform file was applied to each atlas as needed using the `apply_transforms` function from ANTS, specifying nearest-neighbor interpolation with the `interpolator=`‘`nearestNeighbor`’ to preserve discrete region indices. We provide the MNI152NLin6Asym and MNI152NLin2009cAsym formats of all twelve atlases in the `subcortex_visualization` repository for ease of use; more details on the specific transformations performed for each atlas are provided in the supplementary methods (Appendix IV). These are two commonly used spaces for template alignment of empirical neuroimaging data, and if user volumes are in one of these two spaces, they can directly apply any of our twelve atlases to compute regional statistics with `parcel_segstats`, specifying which of the two with the `atlas_space` argument (‘`MNI152NLin6Asym`’ by default).

If the empirical volumetric data are in a different space (either another template or subject space), users wishing to use the `parcel_segstats` function to compute regional statistics must take care to normalize their imaging volume(s) to one of those two spaces. Even after normalization to the same MNI space, users should be aware that there may still be alignment mismatches between the empirical volumetric data and the atlas(es) included in this package, which can lead to inaccurate regional statistics if not properly addressed. For example, the empirical data may have different voxel sizes and/or different affine transformations, which is important to consider given the small subcortical structures included in many of the atlases included here [82]. The `parcel_segstats` function checks the affine alignment and dimension sizes between the empirical input volume and the specified atlas in the specified MNI space, and if one or both do not match, the function will raise an error by default to flag this with the user. The user does have the option to specify an interpolation method (‘nearest’ by default, for nearest neighbor) to resample the atlas into input empirical volume space using either the `resample_img` function from `nilearn` [76] for the Python version or the `resampleImageToTarget` function from the `ANTsR` package [81] for the R version, with a warning message still printed in either case. Generally speaking, we encourage users to take great care with atlas alignment and space interpolation; for those wanting to learn more about these topics, we refer the interested reader to Evans et al. [83].

### 2.7 An empirical case application with neurotransmitter receptor maps

Since `parcel_segstats` accepts any summary statistic function, users have flexibility to choose the most appropriate parcellation scheme and summary statistic for their research question, with the resulting regional summary statistic data pre-formatted for compatibility with the `plot_subcortical_data` function. To demonstrate this functionality, we provide a worked example using neurotransmitter receptor and transporter PET density maps curated by Hansen and colleagues [19, 51, 52]. These maps are openly available from GitHub (https://github.com/netneurolab/hansen_receptorvar) as well as the `neuromaps` Python package [52]. There are currently twenty distinct group-averaged PET maps in this dataset, spanning seven distinct neurotransmitter systems and pre-synaptic terminal density.

As depicted in Fig. 9A, we selected group-averaged PET maps for four distinct neurotransmitter receptors for our applied case study: Mu opioid receptor (MOR) [84–86], 5HT1a serotonergic receptor [87], SV2a synaptic vesicle protein [88], and CB1 cannabinoid receptor [89]. For each map, we selected a different summary statistic to aggregate across voxels per region: the mean for MOR, the standard deviation (SD) for 5HT1a, the maximum for SV2a, and the median for CB1. All group-averaged PET volumes were provided in MNI152NLin6Asym space at 1mm isotropic resolution, so we selected this normalized space for all atlases queried in `parcel_segstats`. The resulting regional statistics were then plotted for the left hemisphere with `plot_subcortical_data` per atlas, as shown in Fig. 9B. We include a worked example in Code snippets 17 (Python) and 18 (R), showing how to compute the median across voxels of the CB1 receptor density across all regions in the aseg subcortical atlas as one example; the syntax is the same for any specified empirical volumetric map, atlas, and desired summary statistic. Of note, many receptor maps are quantified using either the standardized uptake value (SUV) or binding potential (BP_ND_), which generally involve normalizing the raw PET signal to a reference region that is typically the cerebellum (either whole cerebellum or cerebellar gray matter) [19]—with the exception of the MOR tracer, which was normalized to the occipital cortex. This is an important consideration when selecting which atlas to use for visualization of a given PET volume, since resulting values in the cerebellar lobules are more likely to reflect noise rather than true receptor density.

**Figure 9.**
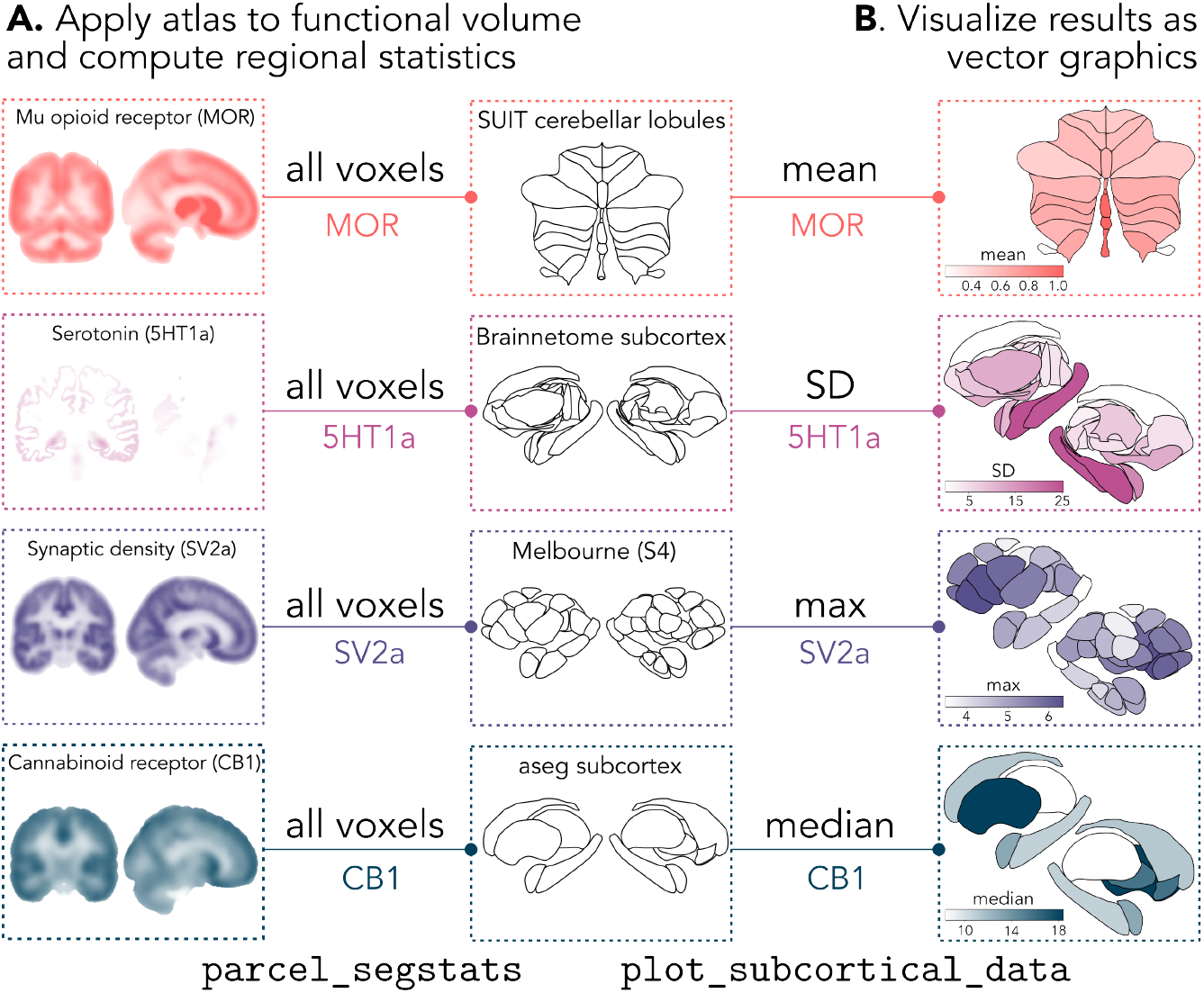
Users can combine any summary statistic function with any of the included atlases to compute regional statistics from their own data. **A**. Any 3D NIfTI image can be used as input data, so long as it is registered to one of the two template spaces supported by the toolbox (MNI152NLin6Asym or MNI152NLin2009cAsym). Here, we selected four example neurotransmitter receptor density PET maps (all shared in MNI152NLin6Asym space) from Hansen et al. [19, 51] and Markello et al. [52] as example input data: Mu opioid receptor (MOR) [84–86], serotonin (5HT1a) [87], pre-synaptic terminal density (SV2a) [88], and cannabinoid receptor (CB1) [89]. For each map, we chose a different atlas and summary statistic function to demonstrate the flexibility of the toolbox. Specifically, we computed the mean value of the MOR map across voxels within each region of the SUIT cerebellar lobule atlas, the standard deviation (SD) of the 5HT1a map within each region of the Brainnetome subcortex atlas, the maximum (max) of the SV2a map within each region of the Melbourne Subcortex Atlas (S4 resolution), and the median value of the CB1 map within each region of the aseg subcortex atlas. **B**. The resulting summary statistics are plotted for each atlas and functional map using the `plot_subcortical_data` function. An example of this workflow for the bottom row (median CB1 receptor density across regions in the aseg atlas) is shown in Code snippets 17 (Python) and 18 (R).

#### 2.7.1 Applying multiple atlases simultaneously

For any given research question and empirical brain-mapping dataset, there is often no singular optimal parcellation scheme or summary statistic, and it can be informative to compare results across multiple parcellation schemes and/or summary statistics. To facilitate this, the `parcel_segstats` function allows users to specify multiple atlases simultaneously (in the format of a list of atlas names), in which case the resulting output `DataFrame` will contain the user-specified regional statistic(s) across all specified atlases. As long as the empirical neuroimaging volume to quantify is in one of the two spaces compatible with our atlases (MNI152NLin6Asym or MNI152NLin2009cAsym), users can flexibly choose any combination of statistics and atlases to extract and visualize region-averaged results.

We demonstrate this functionality in Fig. 10A, in which we compute and visualize the regional average GABA_A_-*α*1 receptor subunit density [90] for all twelve atlases included in this package. This represents one of nineteen known receptor subunits for GABA_A_, a ligand-gated chloride ion channel subtype of GABA, the most abundant inhibitory neurotransmitter in the brain [91, 92]. The GABA_A_-*α*1 receptor subunit was measured using the [^11^C]Ro15-4513 PET tracer and quantified as the total volume of tracer distribution (V_T_), derived using kinetic modeling with an arterial input function [90]—and is not normalized to a reference region, enabling visualization in the cerebellar lobules. We include a worked example for this functionality in Code snippets 20 (Python) and 21 (R). This comparison highlights the variability in regional receptor density patterns across different parcellation schemes, which can be informative for selecting the most appropriate atlas for visualizing a given empirical dataset. For example, GABA_A_-*α*1 V_T_ receptor density values visually differ between thalamic nuclei in the HCP-based atlas from Najdenovska et al. [43] and the THOMAS atlas from Su et al. [44], which is likely because the two atlases subdivide the thalamus into different numbers of nuclei (7 versus 11) with different spatial boundaries. More generally, the regional average values for any given functional map will likely differ across regions between any two atlases due to different boundaries and spatial averaging.

**Figure 10.**
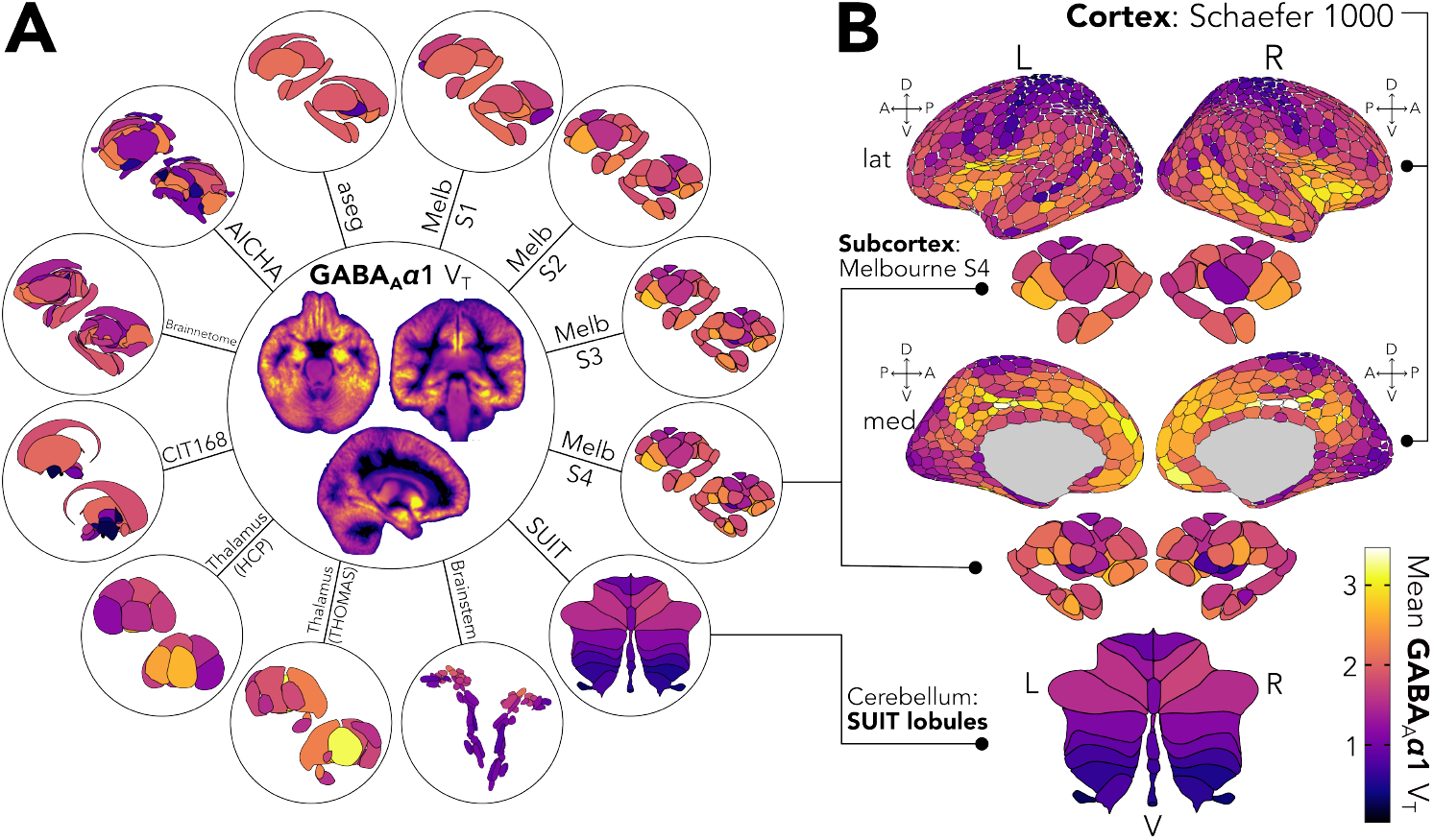
Users can visualize summary statistics in all supported subcortical and cerebellar atlases simultaneously alongside cortical parcellation results. **A**. We use the GABA_A_-*α*1 PET tracer as an example, showing the same summary statistic (the mean regional V_T_ value) across all supported subcortical and cerebellar atlases. These subcortical and cerebellar plots were created using Code snippets 20 (Python) and 21 (R). **B**. To demonstrate how these subcortical and cerebellar vector visualizations can be combined with cortical parcellation results, we applied the 1000-region Schaefer cortical atlas to the same GABA_A_-*α*1 PET tracer data from **A** and visualized the cortical maps using the `ggseg` package in R. We selected the Melbourne Subcortex Atlas (S4 resolution) and SUIT cerebellar lobule atlas as representative examples of the subcortical and cerebellar atlases supported by our toolbox, respectively, visualized alongside the cortical Schaefer atlas results. The color ranges for the subcortical, cerebellar, and cortical maps are matched to facilitate visual comparison across the whole brain, with the legend shown in the bottom right corner. The cortical parcellation results were generated using Code snippet 19 (Python) and visualized using Code snippet 22 (R).

#### 2.7.2 Combining visualizations across cortical and non-cortical structures

Since our visualizations were designed in the style of the `ggseg` cortical plots in R [37], we finally demonstrate how to combine cortical and non-cortical visualizations in one figure for comprehensive brain mapping. Importantly, the R functionality with `subcortexVisualizationR` allows the user to easily combine cortical plots from `ggseg` with subcortical and cerebellar plots from `subcortexVisualizationR` in one figure, all within the `ggplot2` [93] framework. Alternatively, Python package users can save outputs from `plot_subcortical_data` in their preferred file format (e.g., .png or .svg) and combine with the saved `ggseg` output from R in the image editor of their choice, from PowerPoint to Inkscape to Adobe Illustrator. As a fully R-based example, we visualize the regional average GABA_A_-*α*1 receptor subunit density [90] from the previous example in the cortex alongside the subcortex and cerebellum together, as shown in Fig. 10B. We selected the 1000-parcel resolution from Schaefer et al. [94] to parcellate the cerebral cortex, which we applied to the PET data mapped to *fsaverage* surface space using the `neuromaps` package [52]. For the subcortex, we selected the Melbourne Subcortex Atlas [8] at the S4 resolution to match the visualizations in Hansen et al. [19]; we additionally applied the SUIT cerebellar lobule atlas [54].

In this workflow (included in Code snippets 19 through 22), we construct a separate `ggplot2` plot for the cortex (using `ggseg` [37]) and for the subcortex and cerebellum (using `subcortexVisualizationR`), respectively, which we then combine into one figure using the `patchwork` package [95]. The resulting composite GABA_A_-*α*1 subunit density figure is shown in Fig. 10B. Note that the color range is consistent across all three plots, allowing for direct comparison of receptor subunit density across the cortical and non-cortical structures. This presents a clear and interpretable way to visualize brain-wide data in one unified figure with consistent rendering conventions, offering a practical alternative to manually combining cortical surface plots with volumetric slices or 3D mesh renderings of the subcortex and cerebellum.

## 3 Creating custom 2D vectorized visualizations for new atlases

The `subcortex_visualization` package provides a set of twelve pre-vectorized subcortical and cerebellar atlases that serve as scaffolds for visualizing region-level summary statistics from user-supplied empirical data. For users with region-level summary statistics in one (or more) of these atlases, either derived through their own analytical workflow or using the `parcel_segstats` function included in this package (described above in Sec. 2.6), the `plot_subcortical_data` function presents a straightforward way to visualize these data in one or more of the included atlases. However, users may wish to map region-level summary statistics onto a different atlas that is not currently included in the package, or to visualize a custom parcellation scheme that they have developed themselves. We note that if—and only if—the user wishes to add a new atlas that is not currently included in the package, they will need to go through the following workflow once per atlas to create the required 2D vector visualization scaffold, which can be reused thereafter for any future visualizations with that atlas using the `plot_subcortical_data` function. In such cases, we actively encourage users to contribute their vectorized scaffolds for new atlases to the project repository for the benefit of the broader research community.

For those interested in creating custom visualizations for their own atlases, we detail two alternative workflows for creating the required vector visualization scaffolds, based on either (1) automated tracing or (2) manual tracing of the rendered surface meshes. We will refer to the pipeline with automated tracing as ‘semi-automated’ since it still requires one manual downstream step: defining the region layering order in a tabular data file (see below for details). The semi-automated pipeline is considerably faster and is programmatically reproducible using the yabplot package [96] and Potrace algorithm [97], making it a good option for atlases with fewer and/or larger nuclei that are more easily resolved (and thus less likely to require much manual editing). This approach is also more scalable for users who wish to create 2D vectorized representations for many atlases (or, perhaps, many individual subject-space parcellations), as it can be easily scripted to batch-process multiple atlases in one go. On the other hand, while the manual tracing workflow is more time-consuming and is not programmatically reproducible, it allows for greater interactive control over mesh rendering and region tracing, which may be a better option for some atlases (e.g., those with many small nuclei in close proximity, which may be difficult to resolve and would benefit from individual manual tracing). With either choice, users should clone the `subcortex_visualization` repository as described above (cf. Sec 2.1) before proceeding.

In general, the approach to create a vector scaffold for a given atlas involves two key steps, as depicted in Fig. 11 with the Melbourne Subcortex Atlas [8] (S1 resolution) as an example: (1) converting the volumetric segmentation of the atlas (Fig. 11A) into a 3D surface mesh (Fig. 11B), and (2) tracing the rendered mesh (Fig. 11B) to create a 2D vector graphic (Fig. 11C) that can be used for region-level data mapping with `plot_subcortical_data`. For both the manual and the semi-automated workflows, we use free open-source tools to promote accessibility and reproducibility, which we detail below. Regardless of which workflow is used to create the initial vectorization, we strongly encourage users to inspect the quality of the resulting vectorized visualization and to perform manual edits as needed to ensure quality control.

**Figure 11.**
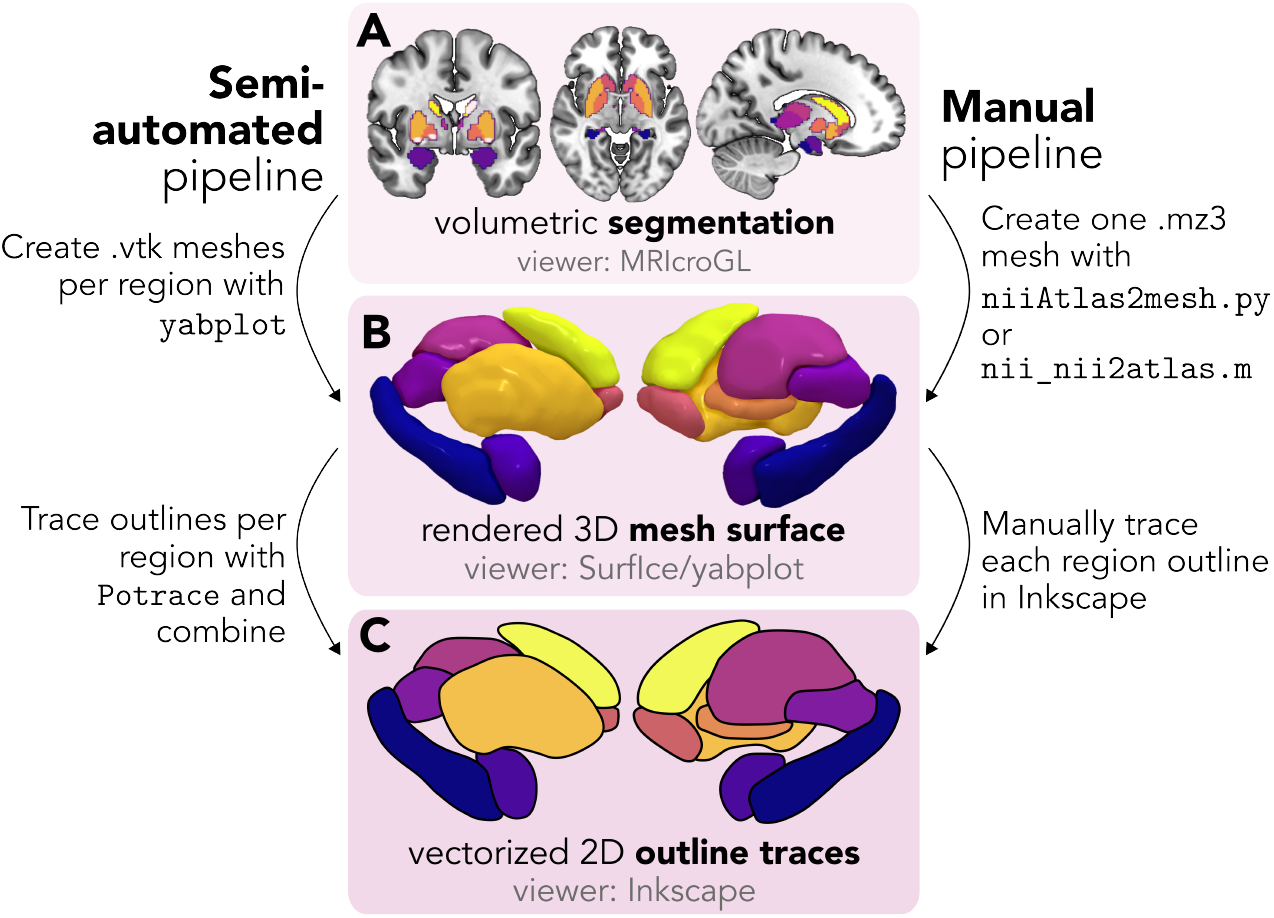
The overall workflow for the custom conversion from volumetric segmentation through to 2D vectorized scaffold images. **A**. For an atlas stored as a volumetric segmentation (e.g., in NIfTI format), this can be viewed as an overlay atop 3D structural images using software like MRIcroGL [98]. **B**. The segmentation volume is then converted to a 3D mesh, highlighting the surface boundaries of the segmented regions. In the semi-automated pipeline, the `build_subcortical_atlas` function from `yabplot` creates individual .vtk mesh files for each region separately, which can be viewed separately or together as an overlay using the `plot_subcortical` function from `yabplot` (which internally calls the `pyvista` library for 3D visualization). In the manual pipeline, the user can choose to convert the entire segmentation volume into a single mesh file, which is then loaded into Surf Ice [99] for interactive visualization and customization of the surface. Rasterized images of the 3D surface (in either pipeline) are captured as bitmap screenshots from the desired views (e.g., medial, lateral, superior, and inferior) for both hemispheres separately and together. **C**. For each segmented region, the edges of the region are traced to create a 2D vectorized image (either automatically with the `Potrace` Python package or manually in Inkscape or similar vector graphics software), in the form of an .svg file.

In order to create 2D vector scaffolds for the twelve atlases included in this package, we used a combination of both the semi-automated and manual tracing workflows, depending on the complexity of the atlas and the quality of the resulting vectorization. Specifically, we applied the semi-automated workflow for all atlases except for the SUIT cerebellar lobule and Brainstem Navigator atlases. For the SUIT cerebellar lobule atlas, the starting representation was already a 2D projected flatmap rather than a 3D volumetric segmentation, so we traced the flatmap directly to create the vector scaffold. For the Brainstem Navigator atlas, the large number of small nuclei in close proximity made it difficult to achieve a high-quality vectorization with the semi-automated workflow, so we opted for the manual tracing workflow with the Surf Ice toolbox [99] and related scripts that enabled us to trace each region individually with greater control over the rendering and tracing process.

### 3.1 Semi-automated workflow

We will first walk through the semi-automated workflow to create a new vectorized scaffold for a given atlas, using the Melbourne Subcortex Atlas (S1 resolution) as an example. As depicted schematically in Fig. 12, this workflow involves the following four steps, which we detail in the following sections: (1) creating a 3D surface mesh for each region in the atlas with the `build_subcortical_atlas` function from `yabplot` [96] (Fig. 12A); (2) rendering each surface mesh with the ‘flat’ lighting style and no cortical mesh overlay using the `plot_subcortical` function from `yabplot` and saving the rendered view as a PNG image (Fig. 12B); (3) automatically tracing the outline of each region in its rendered view with the Potrace algorithm [97] as implemented in the `potracer` Python package to create a 2D contiguous vector graphic (Fig. 12C); and (4) creating a tabular data file to specify the layering order of each region for proper *z*-stacking in the code-based rendering (Fig. 12D). The scripts needed to perform each of these steps for a given atlas are shared with the package repository at https://github.com/anniegbryant/subcortex_visualization/tree/main/adding_new_atlases/semi_automated_pipeline_code and are described in detail in the online documentation^5^, and we include an example shell script call to run this workflow for a new atlas in Code snippet 23.

**Figure 12.**
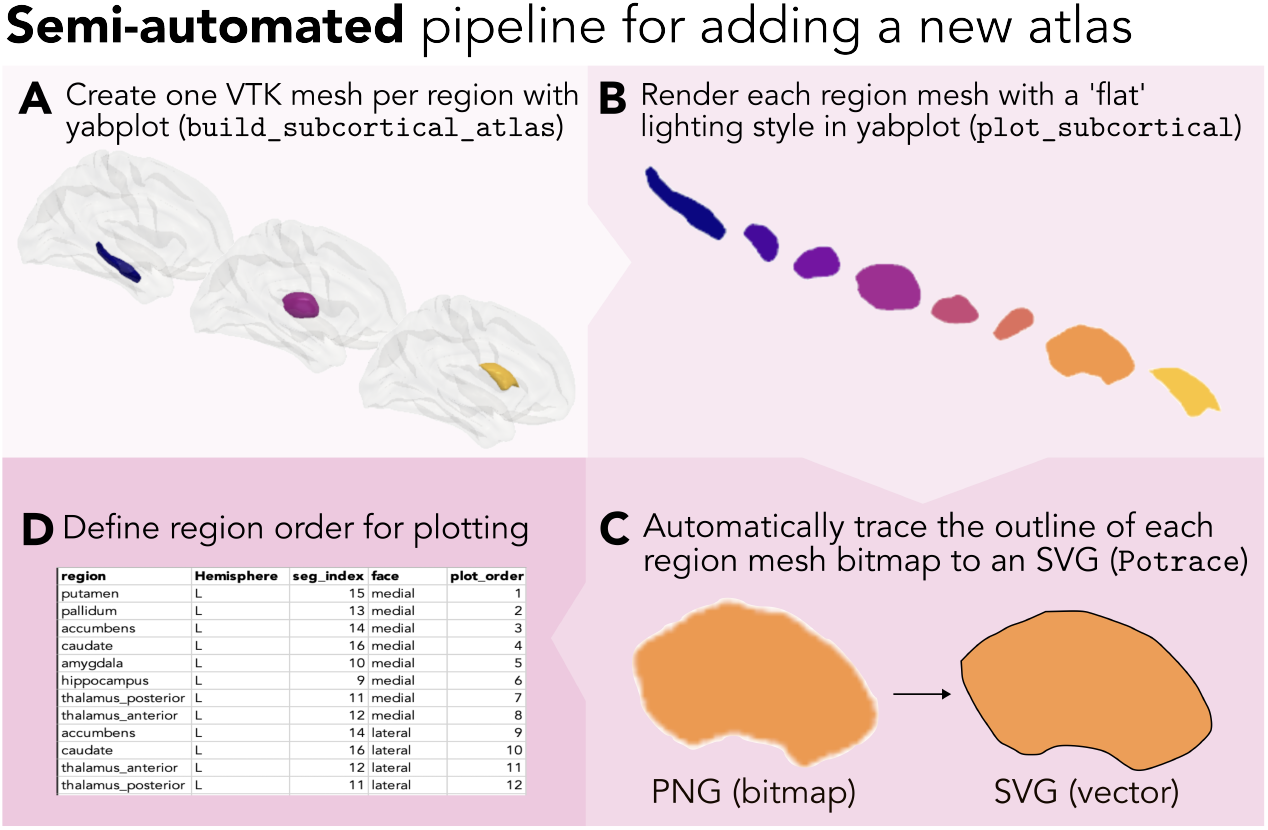
The semi-automated pipeline for creating a vectorized scaffold for a new atlas. **A**. The first step is to create one 3D mesh (in .vtk format) for each region in the atlas, using the `build_subcortical_atlas` function from the `yabplot` package [96]. **B**. Each mesh is then rendered separately with a ‘flat’ lighting style (i.e., no shading) using the `plot_subcortical` function from `yabplot`. **C**. The rendered bitmap (.png) image for each region is automatically converted to a vector object (and saved as an .svg file) using the Potrace tracing algorithm [97] as implemented in the `potracer` Python package. **D**. The region order is manually defined by the user to govern the *z*-axis layering of the regions in the final figure, enabling vectorized visualization with the `plot_subcortical_data` function in the `subcortex_visualization` package.

#### 3.1.1 Building a 3D surface mesh for each atlas region

The first step in the semi-automated workflow is to create a 3D surface mesh for each region in the atlas, which can be done with the `build_subcortical_atlas` function from the `yabplot` package [96]. This function reads in the NIfTI volumetric segmentation of the atlas and applies a marching cubes algorithm to extract the surface mesh for each region, applying Laplacian mesh smoothing with user-defined parameters to reduce artifacts from voxel boundaries. The mesh smoothing parameters are the number of iterations (`smooth_i`, with a default of 15) and the smoothing factor (`smooth_f`, with a default of 0.6), which can be tuned to achieve the desired level of smoothing for a given atlas. For the ten atlases to which we applied this workflow (i.e., all except the SUIT cerebellar lobule and Brainstem Navigator atlases), details about the specific smoothing parameters are given in Appendix IV, Sec. 12. The resulting surface meshes are saved as .vtk files for each region, which can be rendered in the next step.

#### 3.1.2 Rendering each surface mesh and exporting as a PNG image

In order to trace the outline of the surface mesh for each region, we render the mesh for each region separately to isolate the outline of each region in the desired views (i.e., medial, lateral, superior, and inferior per hemisphere). We implemented a modified version of the `plot_subcortical` function from the `yabplot` package [96] to render each region separately with the ‘flat’ lighting style and no cortical mesh overlay, which we found to be optimal for tracing the outline of each region with the Potrace algorithm [97] (described in the next section). The rendered view for each region is saved as a PNG image with a transparent background.

#### 3.1.3 Automatically tracing the rendered mesh to create a 2D vector graphic

For each region, the alpha channel of the PNG image containing the rendered mesh is thresholded at *α* = 128 (i.e., 50% transparency) to create a binary bitmap mask, which is passed to the Potrace algorithm [97] as implemented in the `potracer` Python package (https://github.com/tatarize/potrace). The Potrace algorithm fits a series of lines and Bèzier curves to the boundary of the binary mask governed by three key parameters that can be tuned to achieve the desired level of detail in the resulting vector graphic: `turdsize` (the minimum size of a region to be traced, setting this to 4 pixels rather than the function default 2), `alphamax` (the ‘corner threshold parameter’, dictating whether a vertex should be a corner or a smooth curve, keeping the default value of 1.0), and `opttolerance` (the optimization tolerance for the curve fitting, keeping the default value of 0.2). The other parameter dictating path drawing direction, known as the turn policy, was left at the default setting of ‘minority’. We found that these parameters generally worked well across all ten atlases for which we applied this workflow, but users may wish to tune these parameters for their specific atlas to achieve the desired level of detail in the resulting vector graphic. Potrace adds a full-canvas background rectangle to the resulting vector graphic, which we discard before the SVG files are saved for each region.

#### 3.1.4 Defining the region layering order for proper *z*-stacking in the code-based rendering

Since the rendered vector graphics for each region are separate files, they need to be layered on top of each other in the correct order along the *z*-axis to ensure that regions that are closer to the surface are drawn on top of regions that are deeper in the brain for lateral views (and vice versa for medial views). Similarly, for superior views, regions that are more dorsal should be drawn on top of regions that are more ventral, and vice versa for inferior views. This comprises the fourth and final step in the semi-automated workflow, which involves creating a tabular data file to (manually) specify the layering order of each region for proper *z*-stacking in the code-based rendering. For a given atlas, this requires a separate tabular data file for each hemisphere (and the vermis, in the case of the cerebellum) to specify the layering order for each view (i.e., medial, lateral, superior, and inferior), as well as a combined file for both hemispheres together to specify the layering order for plotting the two hemispheres together. Note that the plot order has to be specified separately for each view, since the layering order of regions can differ across views (e.g., a region that is more dorsal may be deeper in the brain than a more ventral region, but would need to be drawn on top for the superior view versus underneath for the inferior view). The first few rows of an appropriate region ordering table are shown in Table 3, which includes the name of the region, the view, the order in which that region should be drawn (‘plot_order’, with 1 being lowest in the *z*-axis; in other words, placed at the bottom of the stack), the hemisphere, and the index mapping to that region/hemisphere in the original segmentation volume.

**Table 3.** Format of the user-generated region layer ordering in the SVG file.

| region | view | plot_order | Hemisphere | seg_index |
| --- | --- | --- | --- | --- |
| putamen | medial | 1 | R | 7 |
| pallidum | medial | 2 | R | 5 |
| accumbens | medial | 3 | R | 6 |
| caudate | medial | 4 | R | 8 |
| thalamus_anterior | medial | 5 | R | 4 |

For the example atlas ‘My_Atlas’, the file structure for these layer ordering tabular files should look like:

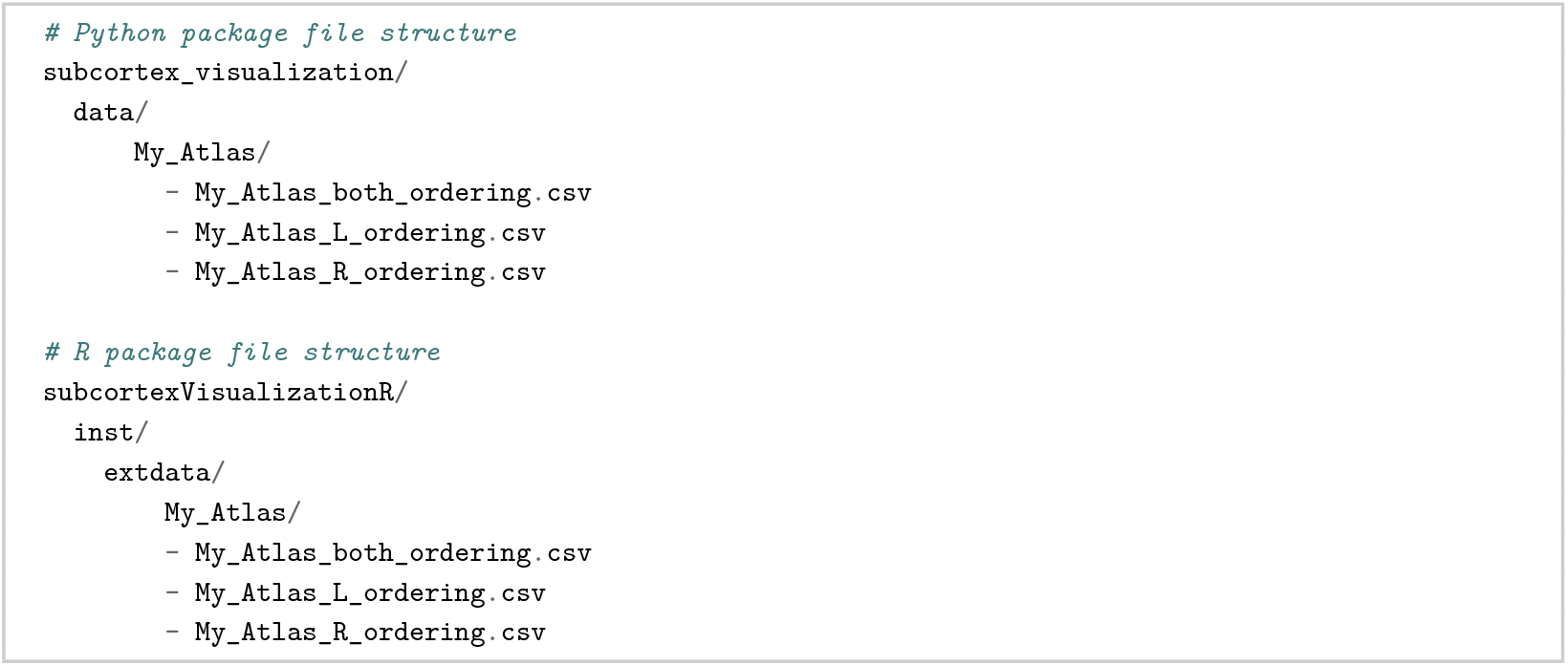

The `plot_subcortical_data` function will automatically read in these files to determine the layering order for each region when plotting with the custom vector scaffold, so it is important to ensure that these files are correctly formatted and placed in the appropriate directory within the package file structure. Constructing these tables generally requires some trial and error to refine the correct ordering for *z*-stacking each region. After creating this region ordering table for a given atlas, the final step is to reinstall the package with the new atlas, which is described below in 3.3.3.

### 3.2 Manual workflow

As an alternative approach to the semi-automated workflow that offers more hands-on control over the final output, users can also create fully interactive 3D visualizations with the Surf Ice toolbox [99] and manually trace the outline of each region in Inkscape (or another vector graphics editor of choice) to create the required vector scaffold for plotting with `plot_subcortical_data`. Since this workflow was only applied end-to-end for the Brainstem Navigator atlas (as the SUIT cerebellar lobule atlas was already provided in 2D), we will walk through this workflow with that atlas as an example, and we share the Python scripts to perform each of these steps for a given atlas with the package repository at https://github.com/anniegbryant/subcortex_visualization/tree/main/adding_new_atlases/manual_pipeline_code (and walk through the steps in the online package documentation^6^). The overall process is depicted schematically in Fig. 13 to summarize the four key steps, which we detail in the following sections: (1) creating a 3D composite surface mesh that includes all regions in the segmentation atlas and viewing it in the Surf Ice software, from which screenshots are captured at the desired view angles (Fig. 13A); (2) tracing the outline of each region with the freehand line drawing tool in Inkscape to create contiguous closed objects for each region (Fig. 13B); (3) populating the object metadata with the appropriate region name (e.g., ‘hippocampus’), view (e.g., ‘medial’), and hemisphere (e.g., ‘L’) for each region in the SVG file (Fig. 13C); and (4) creating a tabular data file to specify the layering order of each region for proper *z*-stacking in the code-based rendering (Fig. 13D).

**Figure 13.**
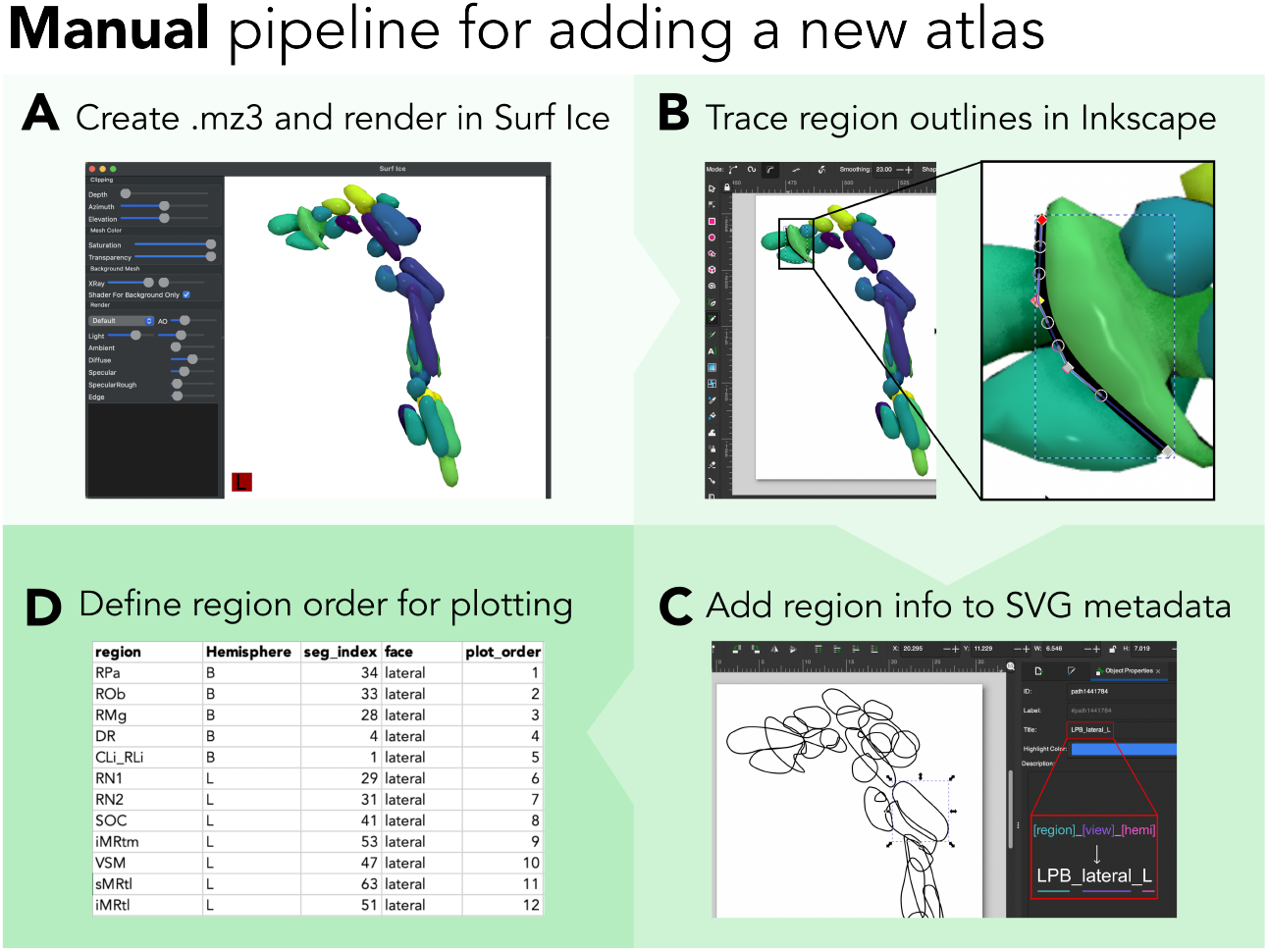
The manual pipeline for creating a vectorized scaffold for a new atlas. **A**. This screen capture depicts the Surfice software rendering the generated .mz3 mesh surface file for the Brainstem navigator atlas (left hemisphere only, including midline structures) using the viridis color map. Users can export a still image of their desired angles (we recommend lateral, as depicted, and medial) using the ‘File > Save bitmap’ action and exporting as a .png image. **B**. Here, the ‘Draw freehand lines’ tool is selected (along the toolbar on the left) and the outline of the left lateral parabrachial nucleus (LPB) is being traced. **C**. Each traced structure is given a name in the Inkscape ‘Object Properties’ panel (shown on the right side of the screenshot). The naming convention used here is `[region]_[view]_[hemisphere]`, where [region] is the name of the brain region in the corresponding atlas (e.g., ‘LPB’); [view] indicates the view to which the traced object corresponds (e.g., ‘lateral’); and [hemisphere] indicates the hemisphere (‘L’ for left, ‘R’ for right, ‘B’ for both, or ‘V’ for vermis).

#### 3.2.1 Rendering a 3D surface mesh

Unlike the semi-automated workflow, which creates a separate surface mesh file for each region, the manual workflow involves creating a single composite surface mesh that includes all regions in the segmentation atlas to be rendered together. Here, we implement a Python-based solution developed by Professor Christopher Rorden at the University of South Carolina, which uses an enhanced marching cubes implementation [100]: niiAtlas2mesh.py (which can be found among other useful Python-based neuroimaging scripts at https://github.com/rordenlab/pythonScripts). This script utilizes the `niimath` library [101]—developed by the same group—which incorporates command-line tool functionality from FSL [75] into Python. Specifically, niiAtlas2mesh.py calls `niimath` (which emulates `fslmaths`) with a series of steps that binarize and smooth each region in the segmentation volume before extracting a mesh surface with an enhanced marching cubes algorithm [100, 102], and finally (slightly) inflating and smoothing the mesh to mitigate small irregularities such as holes or sharp edges. For more discussion about the details of this workflow, we refer the interested reader to Rorden et al. [103]. Together with combinemz3.py, these two scripts first generate individual mesh surfaces for each region in the segmentation volume and then combine them into one cohesive mesh file in .mz3 format. While the .mz3 mesh format can be opened in various neuroimaging applications, it is primarily designed for Surf Ice [99], which we use here; more information about this file format can be found at https://github.com/neurolabusc/surf-ice/tree/master/mz3. As shown in Fig. 13A, the resulting mesh file (`Brainstem_Navigator.mz3`, in this example) is opened in Surf Ice, which renders the entire mesh object with color coding for each region. We recommend rotating the mesh to capture screenshots of the medial, lateral, superior, and inferior views for tracing, as discussed in the next section.

The Python scripts are adapted in our workflow to allow the user to specify a custom color map that assigns an RGBA value to each vertex, corresponding to one color per unique region index. One can construct a new colormap with the desired number of colors (i.e., the number of regions to be visualized in the mesh) in Python as demonstrated in Code snippet 24, which selects 37 evenly spaced shades across the ‘viridis’ colormap from `Matplotlib` [59] for the left hemisphere of the Brainstem Navigator atlas as an example. Note that any colormap can be used (including a custom-generated one), so long as the number of discrete colors to sample matches the number of subregions in the given atlas and number of hemispheres—for example, for the left subcortex in the Melbourne Subcortex Atlas (S2 resolution), 16 colors should be sampled. The script `generate_color_map.py` yields a .txt file with RGBA-formatted colors that can then be supplied to the `volume_to_mesh.py` script with the `–colors` command-line argument, along with the segmentation volume and desired output filename, as shown in Code snippet 25. Generally speaking, we recommend separating out the two hemispheres for maximum visibility along the medial view (i.e., so medial areas are not obstructed by the opposite hemisphere). If there are substantial structural differences between the left and right hemispheres, this process should be repeated separately for each hemisphere; otherwise, the traced regions can simply be duplicated and then mirrored along the horizontal axis, though one should still use the appropriate region name suffixes as discussed below.

One key advantage of this particular mesh construction and visualization approach is that it allows for interactive manipulation of the mesh rendering at high resolution, including zooming, rotating, and adjusting the lighting and shading parameters to achieve the desired level of detail for tracing. However, several alternative graphical user interface (GUI)-based and code-based options are also available to convert NIfTI files into 3D surface meshes, including the `build_subcortical_atlas` function from the `yabplot` package [96] used in the semi-automated workflow described above. For example, Madan [104] provides a detailed tutorial for constructing 3D surface meshes from a volumetric atlas using a combination of the ITK-SNAP [105] and ParaView [106] programs, which we recommend for users who prefer a GUI-based approach. There are also excellent resources for web-based mesh surface rendering through the NiiVue package (https://niivue.com/demos/features/meshes.html); for example, the SUIT probabilistic cerebellar lobule atlas is rendered as a mesh with interactive viewing options at https://niivue.com/demos/features/mesh.atlas.suit.html. Alternatively, other programmatic approaches exist for constructing these triangulated meshes from voxel-based volumes (including a Python version of NiiVue with Jupyter notebooks, at https://github.com/niivue/ipyniivue). Any of these approaches can be used to generate the 3D mesh surface as an intermediate step toward creating the final 2D vector graphic.

#### 3.2.2 Tracing the outline of each region

The next step is to trace the outline of each region in the segmentation mesh to create a vector-based graphic. We use the freely available `Inkscape` software for this process (https://inkscape.org/), which allows users to import bitmap images (e.g., screenshots of the mesh renderings from Surf Ice) and trace around each region to create vectorized polygons—though other vector graphic editing software can also be used for this purpose, such as Adobe Illustrator or Blender. One should create a traced object for each region (e.g., putamen) and view (e.g., lateral and medial) separately, as discussed below. The final traced images can be stacked along the *z*-axis and combined with fill colors matching those in the original mesh rendering as shown in Fig. 13A.

After opening the bitmap image (named, e.g., ‘My_Atlas_lateral_L.png’) in Inkscape, one should select the ‘Draw freehand lines’ option from the toolbar on the left-hand side of the window (denoted with a pencil and drawing icon), as shown in a screenshot in Fig. 13B. In this example, we start with the left LPB (lateral parabrachial nucleus) by simply tracing loosely around the border of this region. The line can be enclosed into a polygon encapsulating this region by connecting the line back to the original trace starting point. This process should be repeated to construct one traced polygon for each region visible in the corresponding view angle (i.e., not all regions will be visible in the medial, lateral, superior, and/or inferior views). Once all regions have been traced for the corresponding hemisphere(s) and view angle(s), the final step in Inkscape is to assign a title to each region in the Object properties panel. This step enables the `subcortex_visualization` toolbox to programmatically access each region and map them to user-supplied data for flexible downstream data visualization, using either the `svgpath2mpl` package in Python (https://github.com/nvictus/svgpath2mpl) or the `svgparser` package in R (https://github.com/coolbutuseless/svgparser). As depicted schematically in Fig. 13C, one should select each traced region one by one, and in the ‘Object properties’ panel on the right-hand side (denoted with a wrench icon), assign an appropriate title to the region. Both our Python package (`subcortex_visualization`) and R package (`subcortexVisualizationR`) rely on a consistent naming convention to identify each region, view, and hemisphere for data mapping (if added through this manual pipeline), as color-coded in Fig. 13C: `[region]_[view]_[hemisphere]`, with the example of `LPB_lateral_L` shown here. The `view` can be ‘medial’, ‘lateral’, ‘superior’, or ‘inferior’ and the `hemisphere` can be ‘L’ (left), ‘R’ (right), ‘B’ (‘both’, for bilateral midline regions, like the dorsal raphe nucleus in the Brainstem Navigator atlas [46]), or ‘V’ (‘vermis’, for the surface-based cerebellar atlas [54]).

#### 3.2.3 Defining the region layering order for proper *z*-stacking in the code-based rendering

As with the semi-automated workflow, the final step in the manual workflow is to create a tabular data file to specify the layering order of each region for proper *z*-stacking in the code-based rendering. The same principles apply here as in the semi-automated workflow, such that a separate tabular data file is required for each hemisphere (and another file for both hemispheres together) to specify the layering order for each view (i.e., medial, lateral, superior, and inferior). The first few rows of an appropriate region ordering are shown in Table 3 in the previous section (3.1.4).

### 3.3 Incorporating custom visualizations into the subcortex_visualization package

Once a user has created a new vectorized scaffold for a given atlas with either the semi-automated or manual workflow, they can then use this custom visualization to plot their empirical data with the `plot_subcortical_data` function in the same way as the included atlases. First, the user needs to organize files into the appropriate structure within the package file system in the cloned `subcortex_visualization` repository (cf. Sec 2.1), as described below, and then re-install the package to integrate the new atlas with the existing functionality. The `subcortex_visualization` toolbox is designed to work with two distinct types of file structure, depending on the workflow used to create the vector scaffold for a given atlas. For the semi-automated workflow, there is a separate .svg file for each region in the atlas, and these .svg files are programmatically layered on top of each other in the code-based rendering to create the final visualization. By contrast, for the manual workflow, there is one .svg file per hemisphere (and one for both hemispheres together) for each view that contains all regions in the atlas as separate objects within that file, which are programmatically accessed and colored according to user-supplied data for plotting with `plot_subcortical_data`.

#### 3.3.1 Organizing files from the semi-automated workflow into the package file structure

After running the semi-automated workflow for a given atlas, users should have a directory of individual .svg files for each region in the atlas as well as tabular data files specifying the layering order for each view for the left hemisphere, right hemisphere, and both hemispheres together. These files should be organized in the following structure within the cloned `subcortex_visualization` repository, using the following general format as shown below for the example of ‘My_Atlas’ with the first region ‘RegionA’:

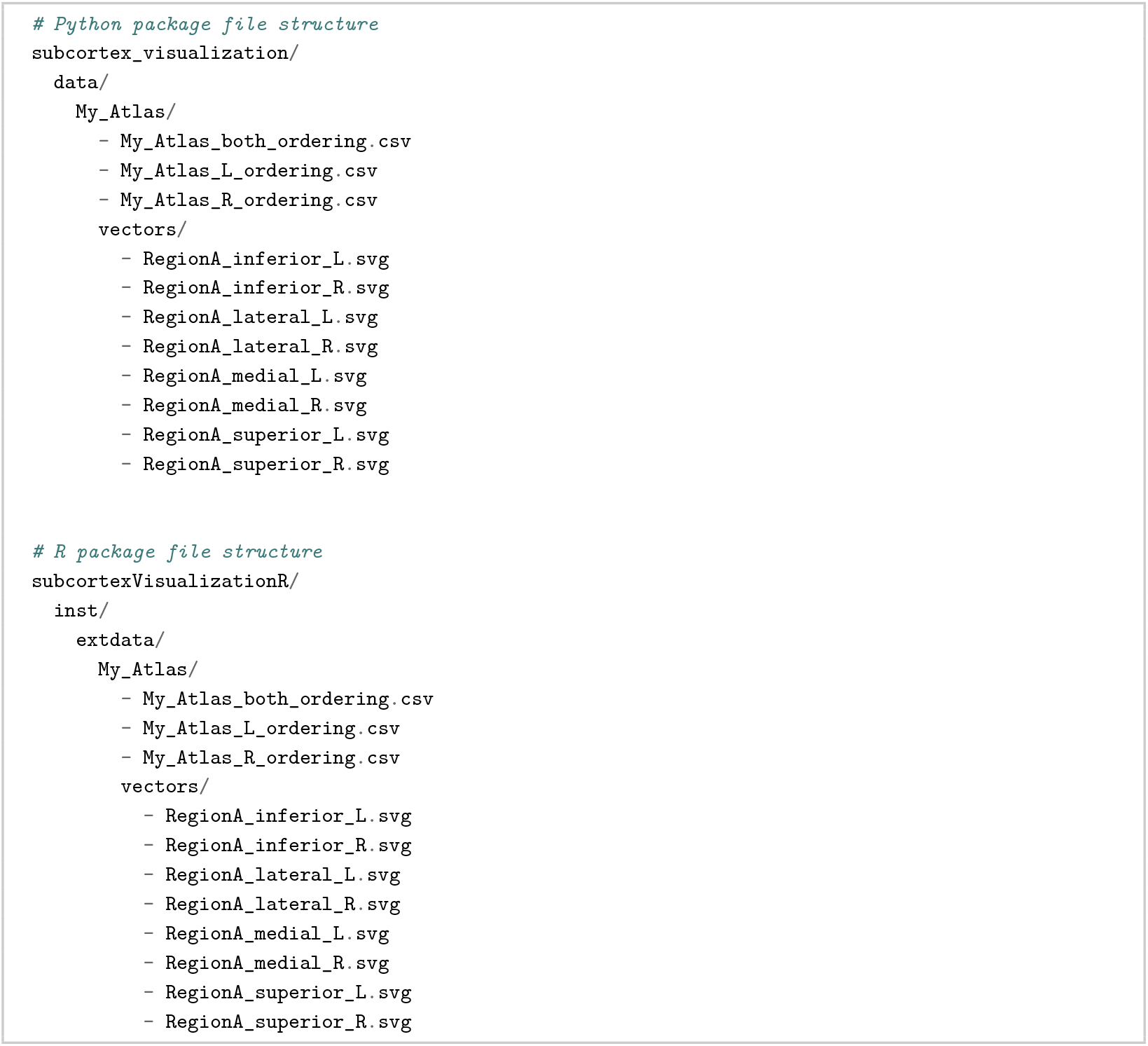

#### 3.3.2 Organizing files from the manual workflow into the package file structure

After running the manual workflow for a given atlas, users should have one .svg file per hemisphere (and one for both hemispheres together) that contains all regions in the atlas as separate objects within that file, as well as tabular data files specifying the layering order for each view for the left hemisphere, right hemisphere, and both hemispheres together. These files should be organized in the following structure within the cloned `subcortex_visualization` repository, using the following general format for ‘My_Atlas’ as an example:

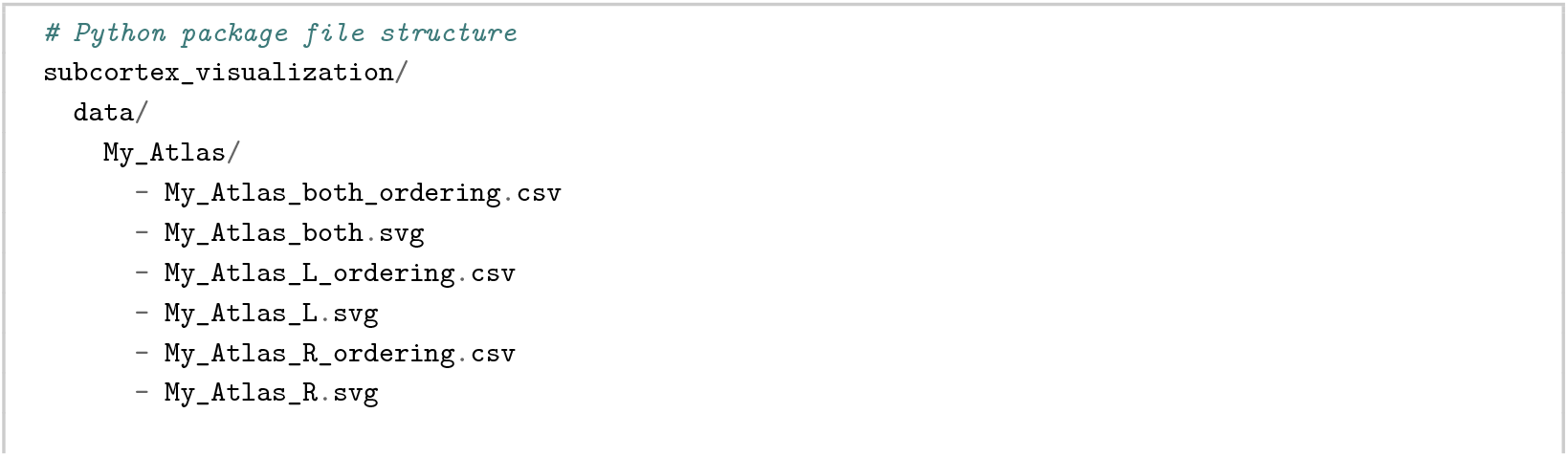

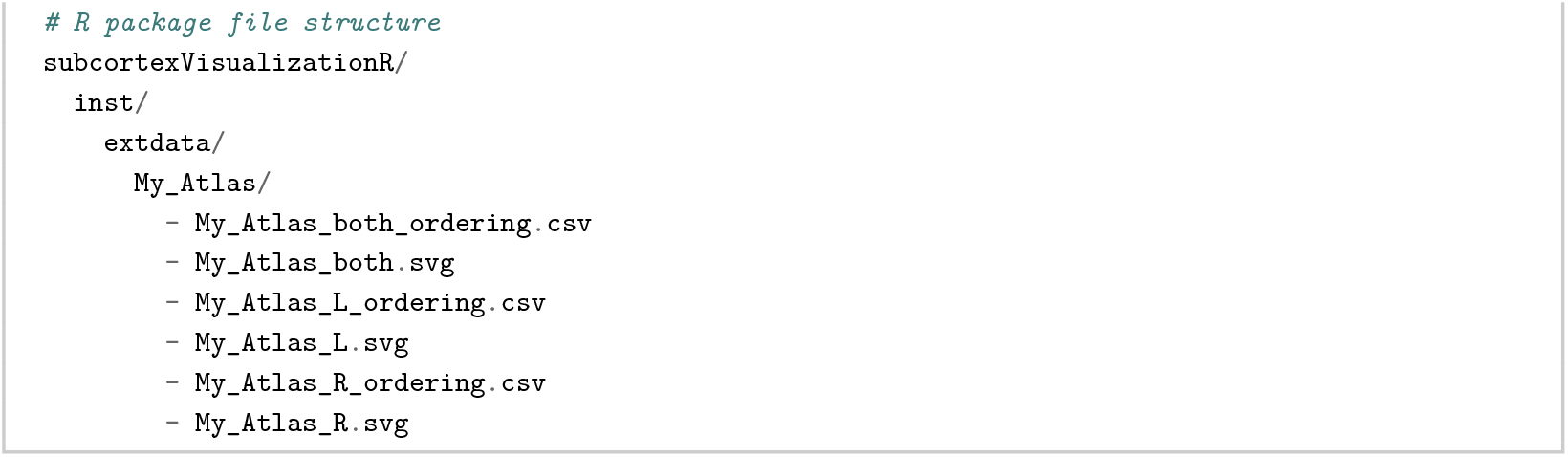

#### 3.3.3 Re-installing the package to integrate the new atlas with existing functionality

After organizing the files for the new atlas into the appropriate directory within the package file structure, the final step is to re-install the package to integrate the new atlas with the existing functionality. The Python package can be re-installed with the following commands from the cloned repository directory (in the terminal):

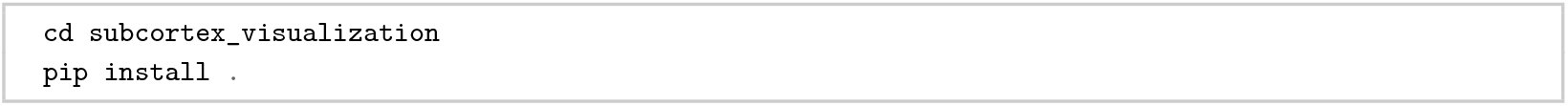

Similarly, for the R package, one should run the following commands, either from the terminal (in an R session) or in RStudio:

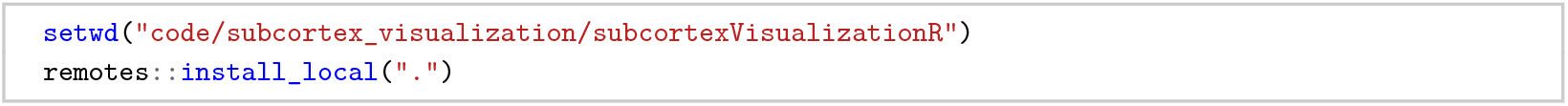

Now, all data visualization functionality introduction in Sec. 2.5 can be implemented with the atlas ‘My_Atlas’. Users are also encouraged to share any added custom vectorized scaffolds for new atlases to the project GitHub repository (https://github.com/anniegbryant/subcortex_visualization) through a pull request, which will allow other researchers to benefit from the expanded library of subcortical and cerebellar atlases.

## 4 Discussion

In this work, we introduce the `subcortex_visualization` toolbox (in both Python and R), with the aim of broadening the availability of non-cortical data visualization tools for the human brain in a consistent and accessible manner. This presents a unified subcortical and cerebellar visualization tool that complements existing software for 2D cortical rendering (e.g., ggseg [37], along with pycortex [107] for cortical flatmap projections) and projection of empirical data onto a 3D surface for cortical and non-cortical structures alike (e.g., ggseg3d [37], ENIGMA toolbox [39], yabplot [96], fsbrain [108], and brainrender [109]). We curated a set of twelve existing atlases that are frequently used in human brain-mapping studies, including eight for the broader subcortex, two for thalamic nuclei, one for the brainstem, and one for the cerebellum. The toolbox allows users to map empirical region-level data derived from myriad domains (from gene expression to receptor density to functional activation) onto any of these atlases with flexibility in color-mapping, region border lines (which are particularly useful to delineate region–region boundaries), view angles, transparency, and statistical significance annotation. To our knowledge, `subcortex_visualization` is the first toolbox to offer standardized and publication-ready vector-based visualizations across multiple commonly used subcortical and cerebellar atlases, filling a critical gap to unify and streamline the visualization of results from below the cortical mantle. Importantly, the vector graphic outputs from this toolbox provide high-quality, resolution-independent rendering that is ideal for publication-quality figures and further editing in vector graphics software (e.g., Adobe Illustrator or Inkscape).

Additionally, we have introduced two complementary workflows for creating new vectorized scaffolds for additional atlases that expand upon open-source resources (including Surf Ice [99] and yabplot [96]), paving the way for flexible incorporation of new atlases across both cortical and non-cortical structures in the future. This collection of atlases (twelve in the present release) is not meant to be exhaustive, but rather to provide a substantial and extensible starting point, and we actively encourage users to contribute their own vectorized scaffolds for new atlases to the project repository for the benefit of the broader research community. For example, the Lead-DBS software and accompanying documentation cover a much broader range of subcortical atlases that are commonly used in the deep brain stimulation field [110, 111], many of which could be added to our toolbox in the future, such as the Sydney Subcortical Gray Matter (SydSGM) parcellation [42], the FreeSurfer probabilistic thalamic nuclei atlas [45], and the Ascending Arousal Network brainstem nuclei segmentation from Olchanyi et al. [47].

We encourage users to explore the wide array of existing tools and approaches for computing region-level summary statistics from raw data that are available in the field [38, 75, 112], particularly given the heterogeneity of potential underlying data sources (including multimodal neuroimaging, multi-omics, and histology) [6, 113]. Our toolbox is designed to be agnostic to the source of the region-level data, and it can thus be used to visualize any region-wise summary statistic that can be mapped to the regions in the included atlases (or any custom atlas added by the user). The `parcel_segstats` function in our package does offer a convenient way to flexibly compute various region-wise summary statistics from a given NIfTI file in any of the twelve atlases in our library (so long as the user-supplied volume is normalized to one of our supported reference spaces). This functionality expands previously limited statistical flexibility among programmatic imaging parcellation workflows, offering a straightforward way to transform raw volumetric data into region-level aggregates that can be visualized in publication-quality and customizable vector graphic figures with the `plot_subcortical_data` function.

One limitation of the current toolbox is that it is exclusively designed for visualizing data at the level of discrete brain regions, and thus does not currently support voxel-wise or vertex-wise visualization. We note that this is a deliberate design choice to focus on region-based visualization, which distills high-dimensional data down to an interpretable set of regions defined by structure, function, connectivity, and/or cytoarchitecture [37, 114]. Given the prevalence of atlas parcellation-based approaches for summarizing and visualizing neuroimaging and bioinformatic data across the brain [21, 115–119], the principal contribution of `subcortex_visualization` is to provide a much-needed resource for creating publication-ready vector-based visuals of summary statistics across a wide range of subcortical and cerebellar atlases simultaneously. On the other hand, aggregating across voxels within a region may not fully capture the richness of spatial variation within that region—a limitation common to parcellation-based approaches for cortical and non-cortical regions alike [120]. Future work could therefore explore the possibility of extending this toolbox to support voxel-wise or vertex-wise data visualization in an atlas-independent manner. This would require a different approach to rendering and visualizing data, particularly for projecting voxel-wise data from 3D volumes onto 2D surfaces for visualization, which is complicated for subcortical regions given their complex geometry as 3D structures and the fact that they are not organized in a sheet-like manner like the cerebral cortex. Alternative approaches could draw upon 3D scatter plot visualizations for different subcortical and cerebellar atlases, as in Wu et al. [28], which depicts voxelwise data without the need for surface-based rendering.

Another limitation of the 2D surface-based rendering approach is that some regions may be obscured by others in certain views, particularly for atlases with many small and densely packed regions (such as the Brainstem Navigator atlas [46], with 37 unique nuclei per hemisphere, including eight bilateral midline structures). The transparency parameter `fill_alpha` can be used to mitigate this issue to some extent, but it may not be possible to fully resolve this issue for all atlases and view angles. One potential future direction to address this issue would be to programmatically increase the spacing between rendered regions from the center outwards. Such a solution could draw inspiration from the shape analysis pipeline developed by the ENIGMA Consortium to project data onto each subcortical region mesh individually [18, 121]. However, the regions are not inherently arranged to preserve their spatial relationships with one another in this pipeline, and it is a notable technical challenge to programmatically increase the spacing between regions while preserving their relative spatial relationships and anatomical fidelity. Alternatively, although it would not directly resolve over-plotting of regions, future work could explore programmatically adding region label text annotations to the visualizations. This could help to disambiguate regions that are close together and difficult to distinguish based on border lines alone. Automated (or even semi-automated) label placement is a non-trivial problem in data visualization, for brain-mapping and cartography alike [122, 123], however, and we will leave this as a future direction for development of this feature in our toolbox.

Altogether, by providing both out-of-the-box vectorized representations of several atlases and instructions with which users can further expand this vector scaffold library, the `subcortex_visualization` toolbox represents a crucial resource for data visualization options below the cortical mantle. We hope that this serves as an important step towards increasing the availability of open-source programmatic data visualization tools for non-cortical structures in the human brain, which can be readily integrated with other R- and Python-based imaging analysis workflows. Future work could also extend this toolbox to support standardized visualizations of non-human brain atlases for comparative neuroanatomy [124], such as the rhesus macaque [125], marmoset [126], and mouse [127]. Recognizing the importance of open science and community engagement, we have made this package available on GitHub at https://github.com/anniegbryant/subcortex_visualization alongside documentation at the project website (https://anniegbryant.github.io/subcortex_visualization/).

## 5 Acknowledgments

A.G.B. was supported by an Australian Government Research Training Program (RTP) Scholarship and the Paulette Isabel Jones Career Award, both granted by The University of Sydney. We are indebted to Sidhant Chopra, Christopher Rorden, Justine Hansen, Ye Tian, and Toomas Erik Anijärv for their suggestions and continued development of open tools for neuroimaging visualization that enabled the development of this project. We also wish to thank Giulia Baracchini for critical discussion about visualization design, R implementation, and the current subcortex visualization landscape. We are also grateful to package users for their feedback and suggestions for improvement, which have been invaluable in refining the functionality of this toolbox. Finally, we thank Ben D. Fulcher and James M. Shine for their mentorship, feedback, and financial support throughout this project, as well as their broader commitment to open science and reproducibility in neuroscience, which has been a major inspiration for this work.

## 6 Ethics

This study did not involve the collection of any new human data. All demonstrations use previously published, openly available datasets, and no additional ethical approval was required.

## 7 Competing interests

The author has no competing interests to declare.

## 8 Author contributions

A.G.B. developed the software and wrote the manuscript.

## 9 Data and code availability

All code for `subcortex_visualization` is openly available at https://github.com/anniegbryant/subcortex_visualization. The PET data used in this work was curated by Hansen and colleagues [19, 51, 52] and is openly shared at https://github.com/netneurolab/hansen_receptorvar.

## Appendix I Code supplement

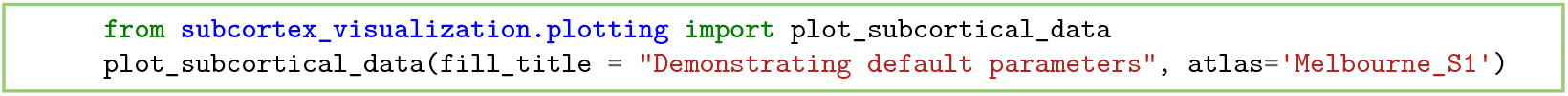

**Code snippet 1**. Demonstrating default parameters of `plot_subcortical_data` [Python].

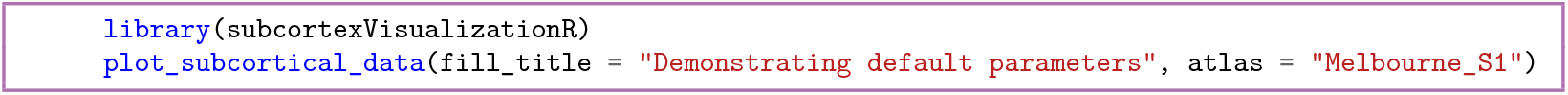

**Code snippet 2**. Demonstrating default parameters of `plot_subcortical_data` [R].

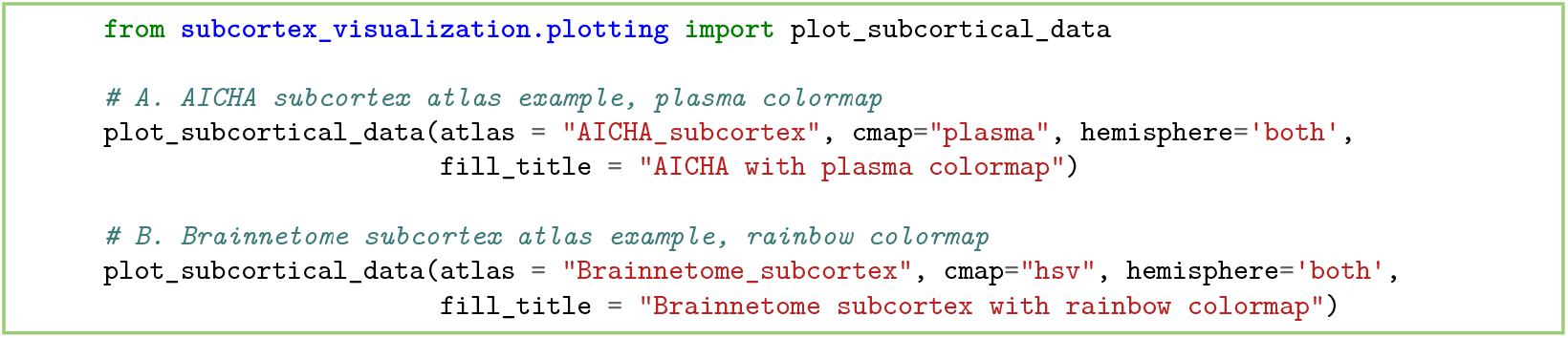

**Code snippet 3**. Visualizing the AICHA and Brainnetome subcortex atlases with one color per subregion [Python].

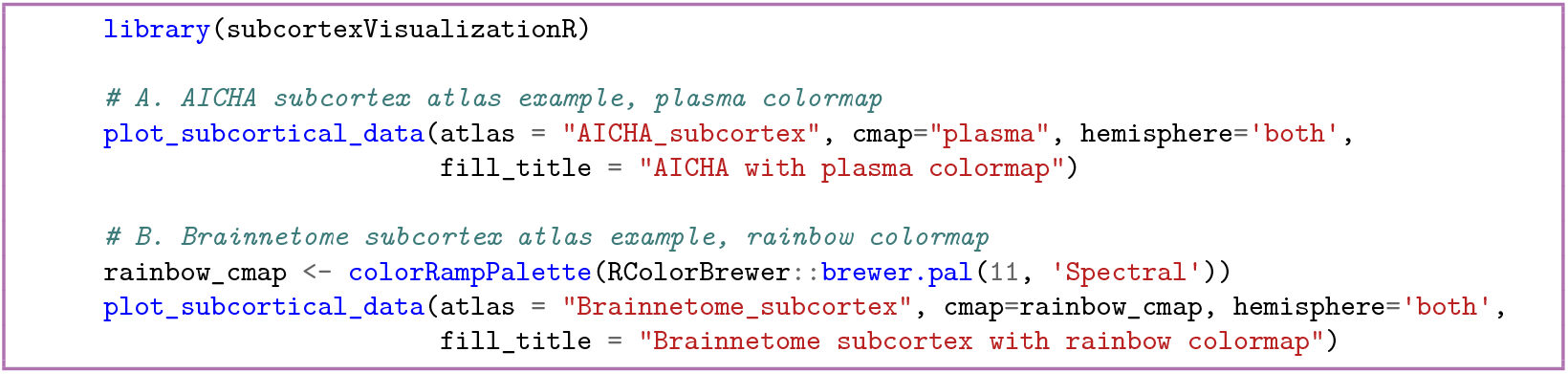

**Code snippet 4**. Visualizing the AICHA and Brainnetome subcortex atlases with one color per subregion [R].

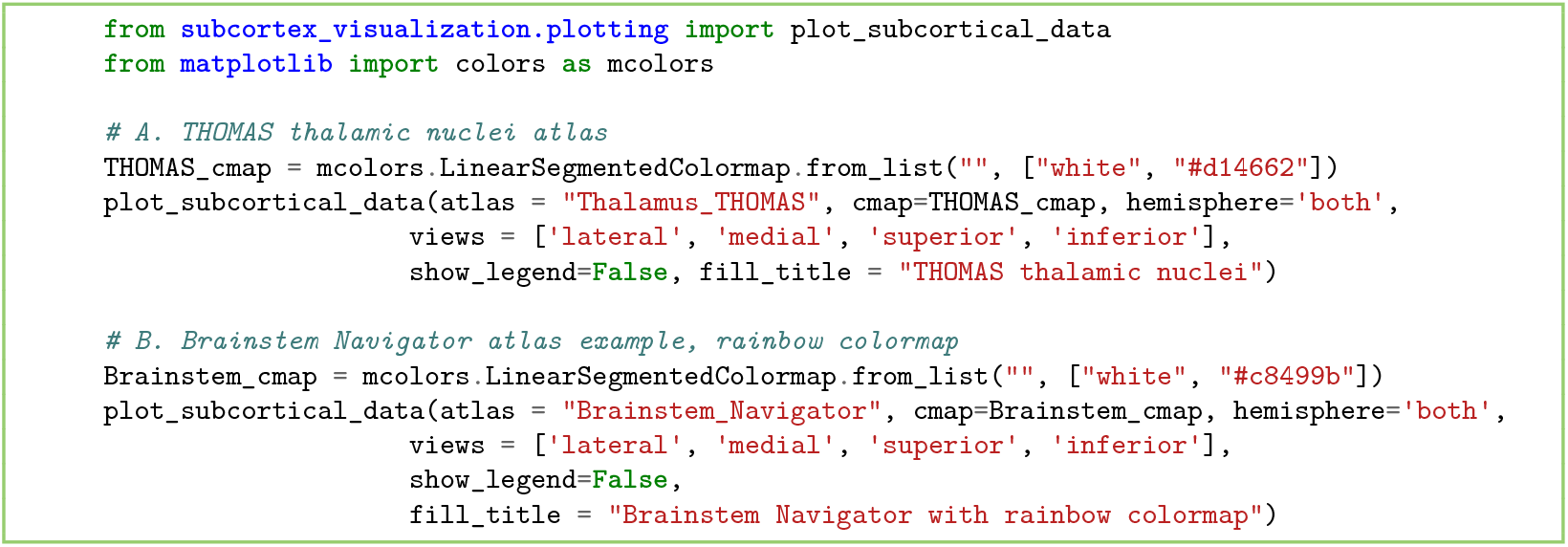

**Code snippet 5**. Demonstrating the four available view angles with the THOMAS thalamic nuclei atlas and Brainstem Navigator atlas [Python].

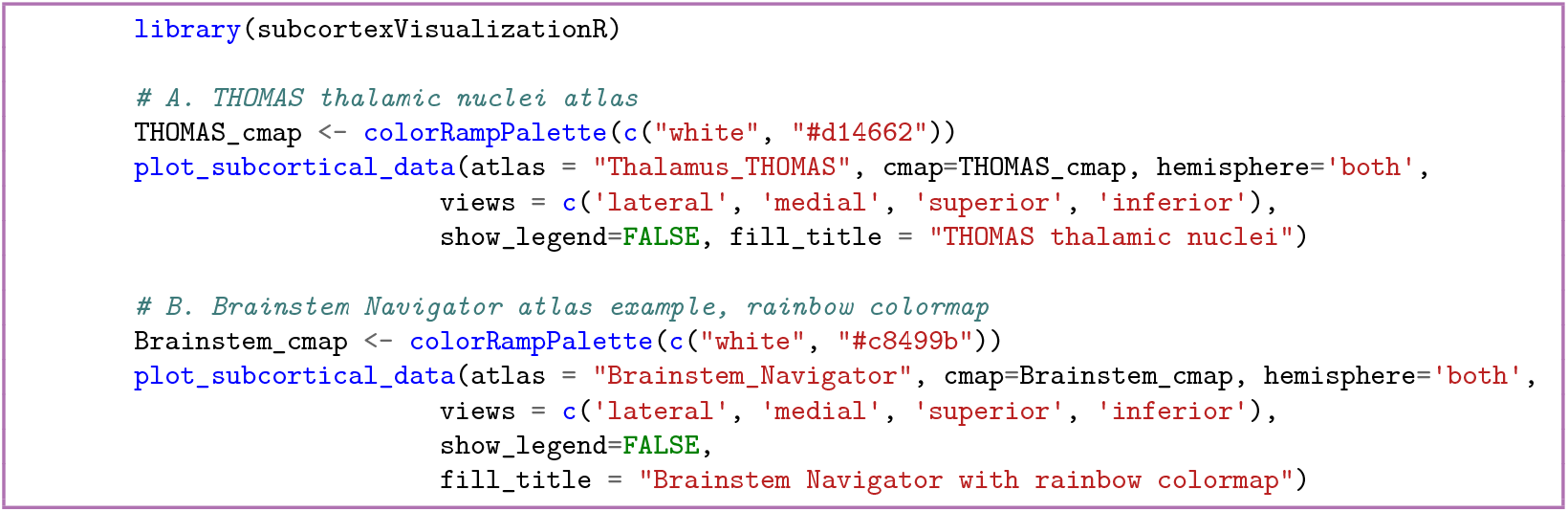

**Code snippet 6**. Demonstrating the four available view angles with the THOMAS thalamic nuclei atlas and Brainstem Navigator atlas [R].

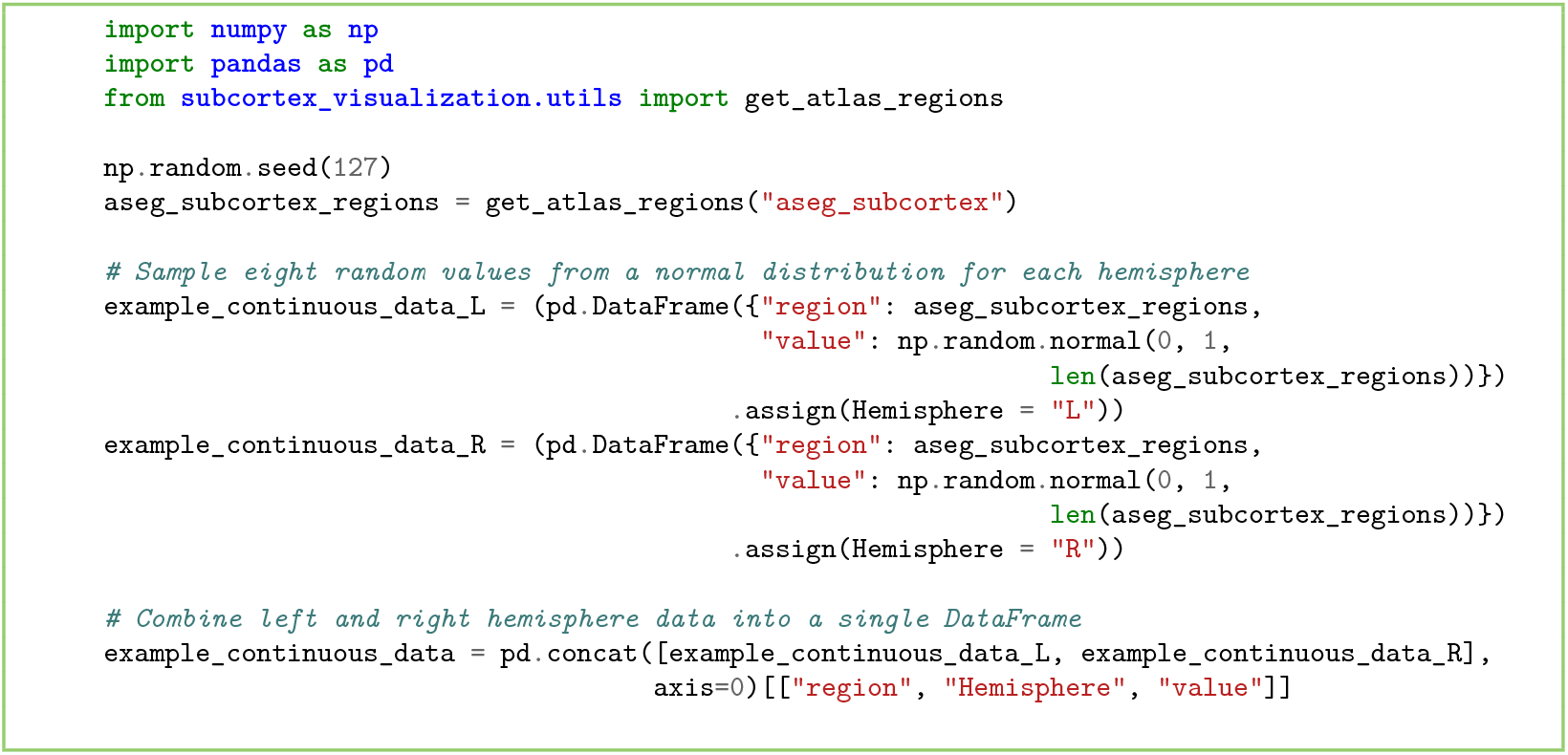

**Code snippet 7**. Simulating empirical data for the left hemisphere of the aseg subcortex atlas [Python].

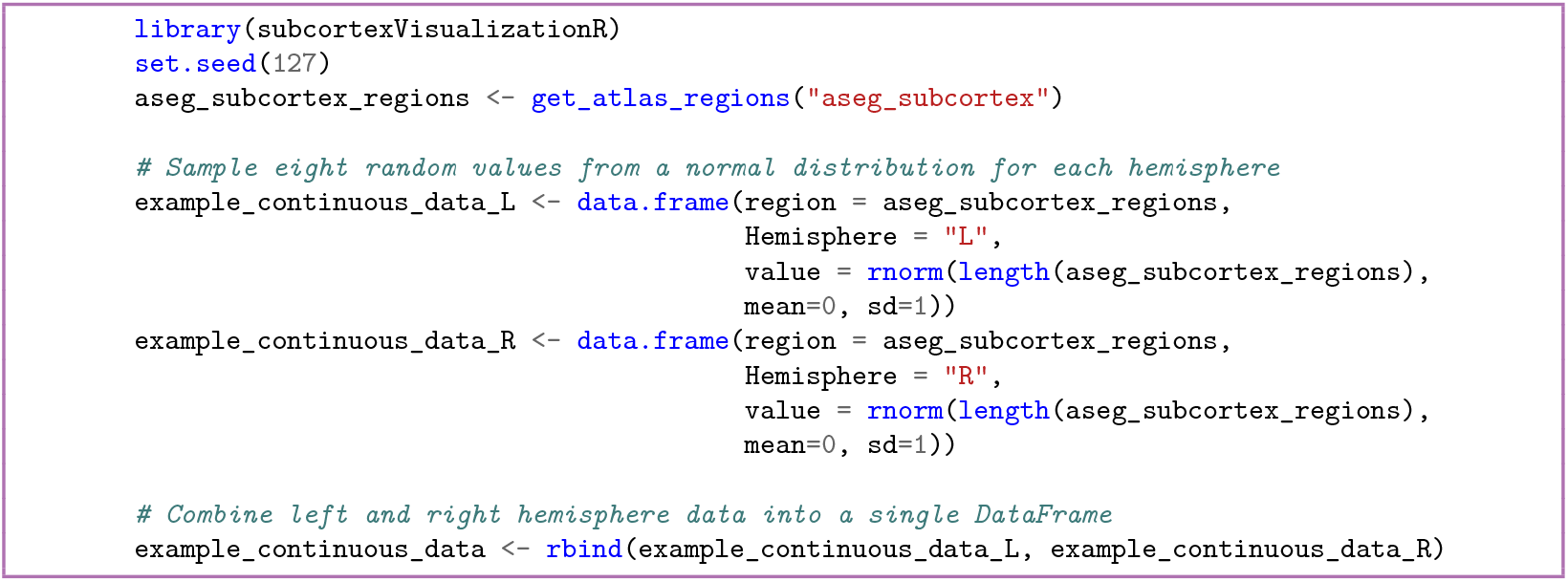

**Code snippet 8**. Simulating empirical data for the left hemisphere of the aseg subcortex atlas [R].

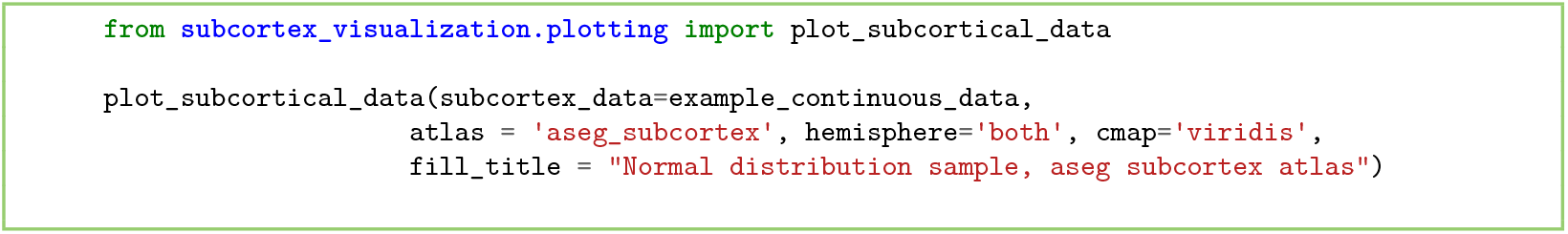

**Code snippet 9**. Plotting simulated data for the left hemisphere of the aseg subcortex atlas [Python].

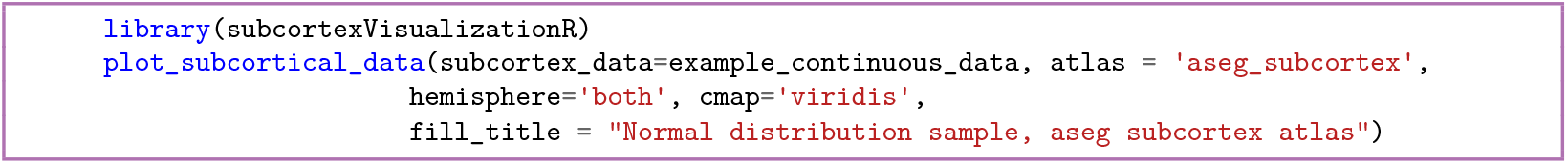

**Code snippet 10**. Plotting simulated data for the left hemisphere of the aseg subcortex atlas [R].

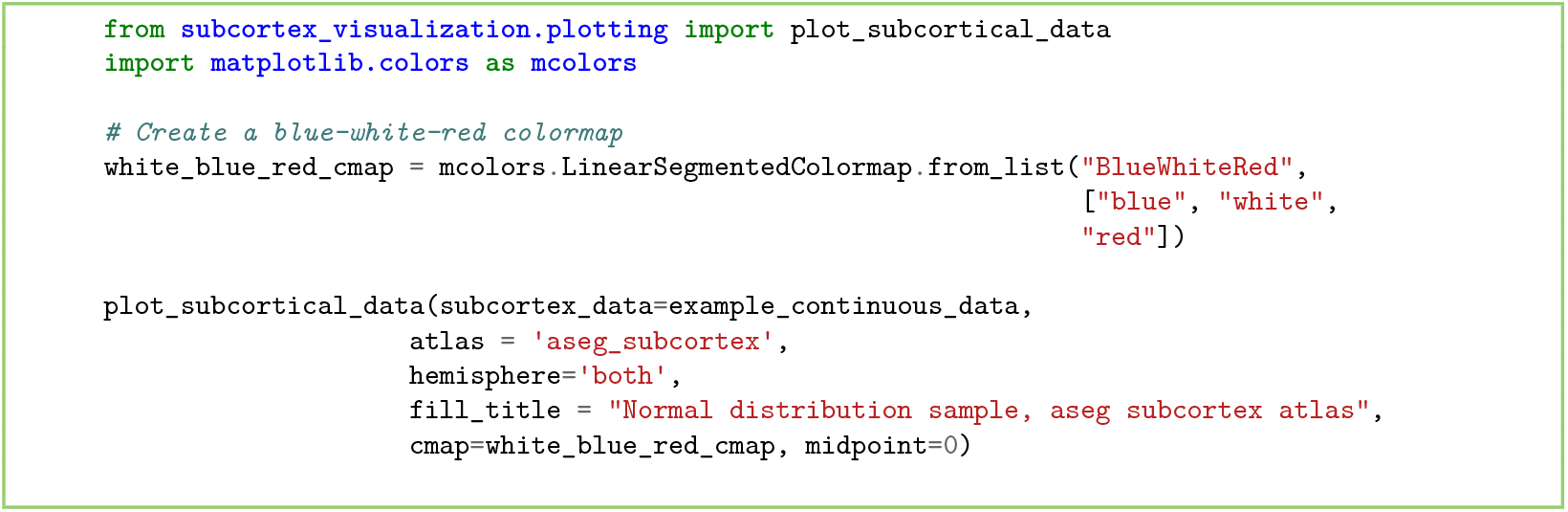

**Code snippet 11**. Plotting simulated data from Code snippet 7 with a custom colormap for the aseg subcortex atlas, left hemisphere [Python].

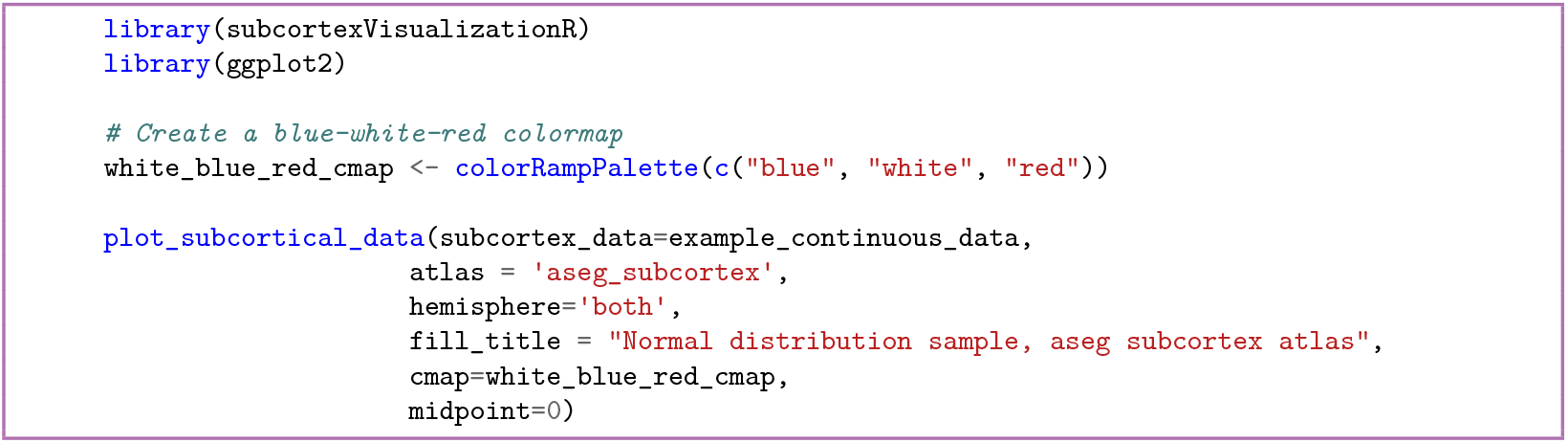

**Code snippet 12**. Plotting simulated data from Code snippet 8 with a custom colormap for the aseg subcortex atlas, left hemisphere [R].

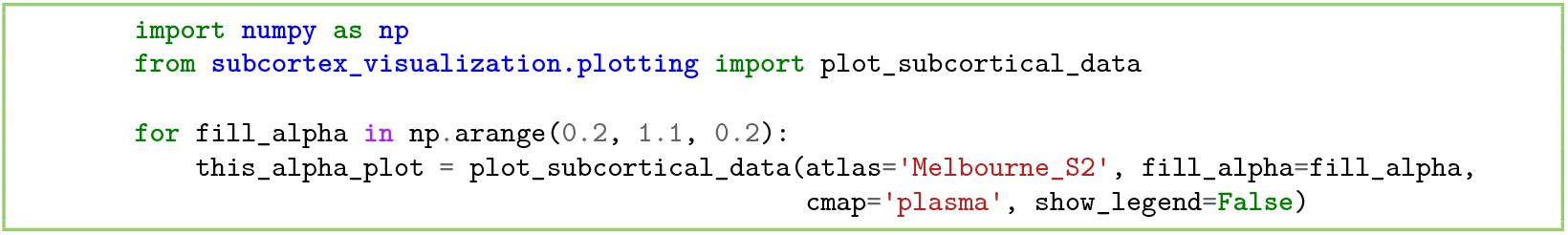

**Code snippet 13**. Demonstrating the `fill_alpha` transparency parameter [Python].

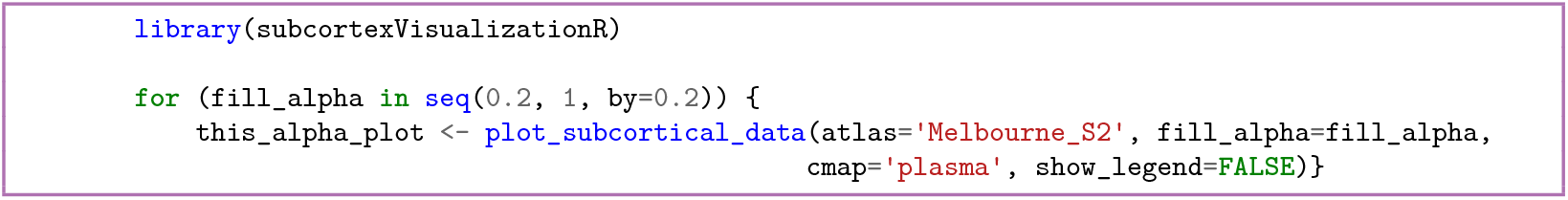

**Code snippet 14**. Demonstrating the `fill_alpha` transparency parameter [R].

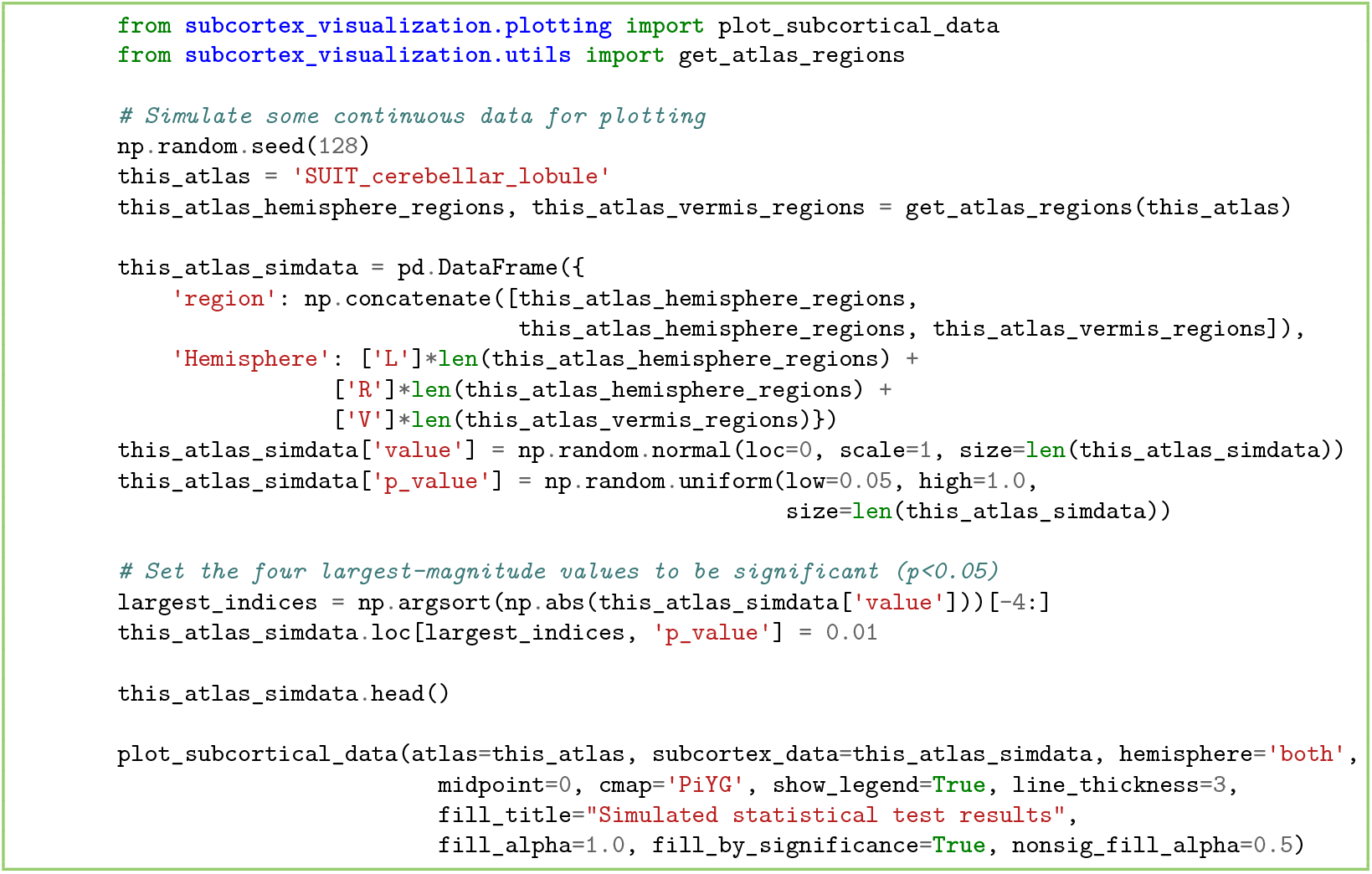

**Code snippet 15**. Simulating statistical significance data to visualize in the SUIT cerebellar lobule atlas [Python].

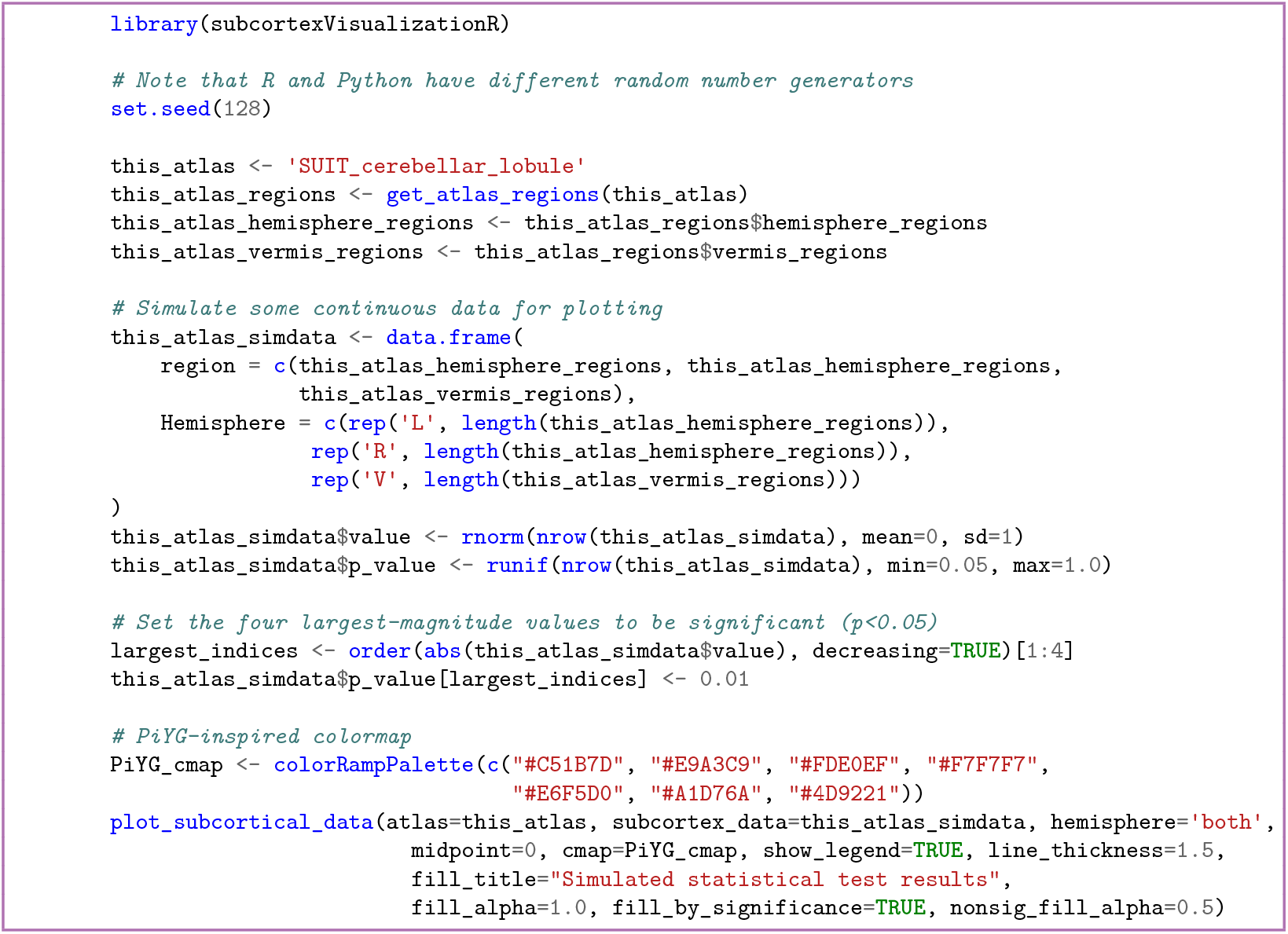

**Code snippet 16**. Simulating statistical significance data to visualize in the SUIT cerebellar lobule atlas [R].

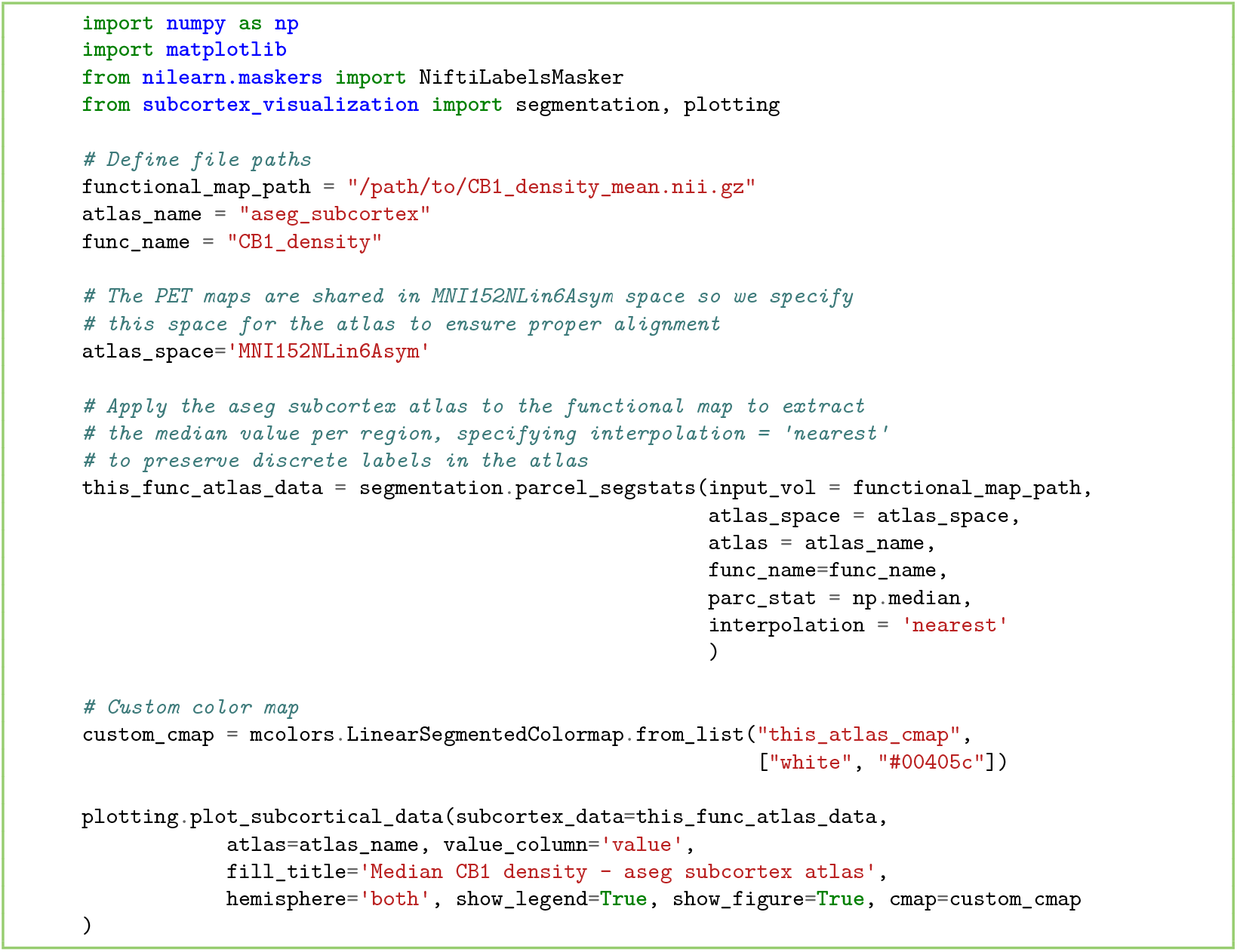

**Code snippet 17**. Computing and plotting region-averaged CB1 densities in the aseg subcortical atlas [Python].

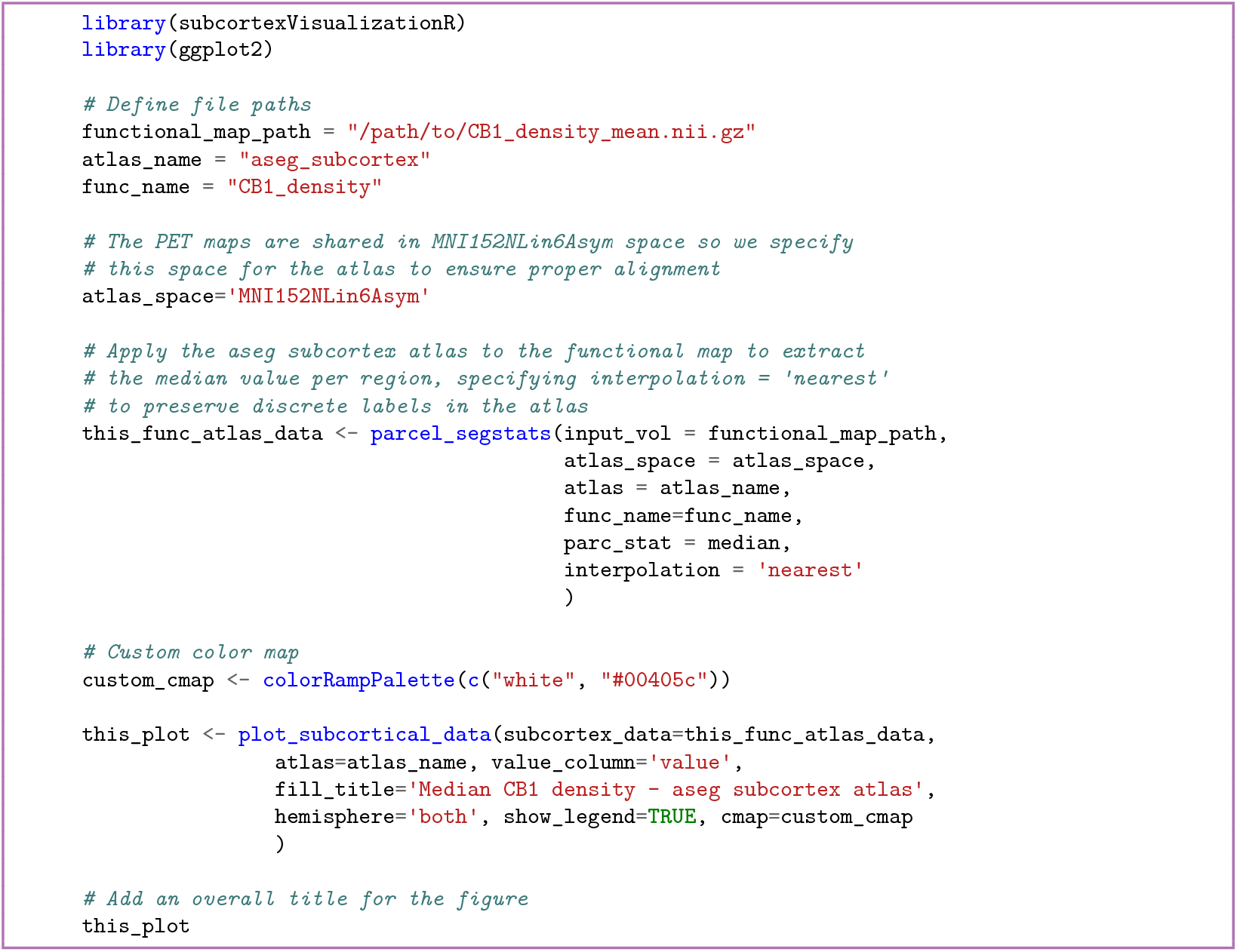

**Code snippet 18**. Computing and plotting region-averaged CB1 densities in the aseg subcortical atlas [R].

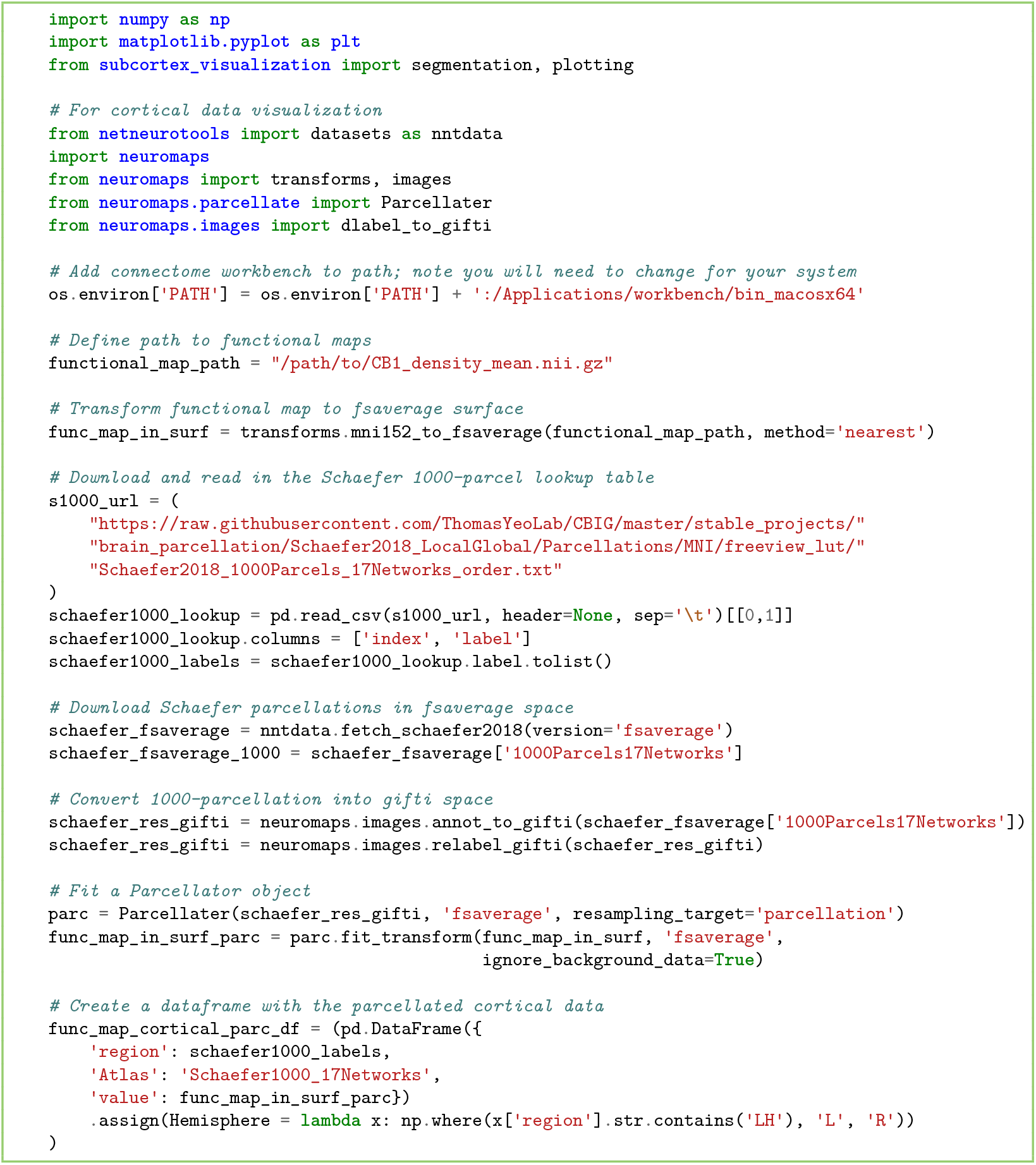

**Code snippet 19**. Computing the region-averaged GABA_A_-*α*`1` densities for the Schaefer-1000 cortical parcellation [Python].

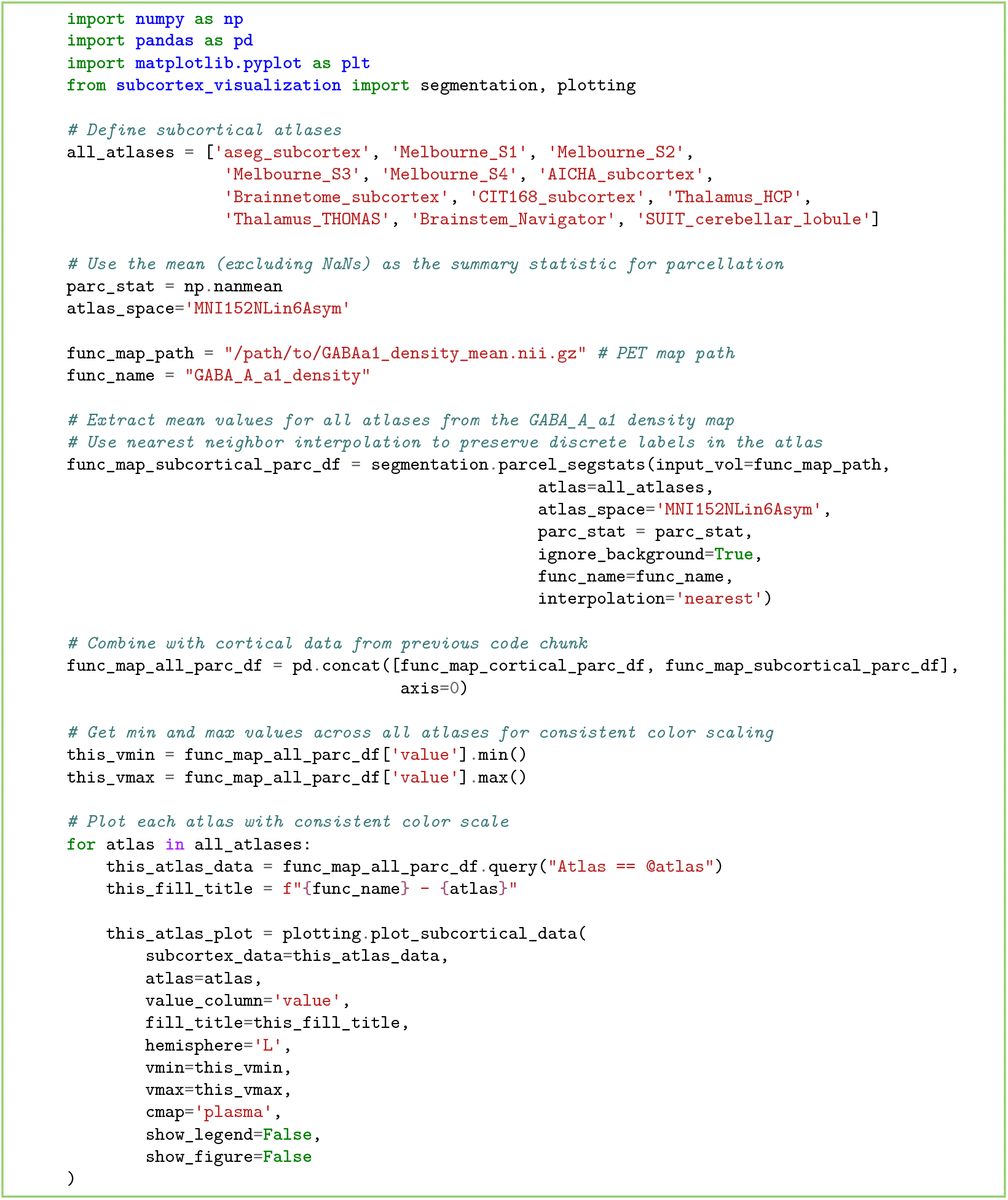

**Code snippet 20**. Computing and plotting the region-averaged GABA_A_-*α*`1` densities in all twelve provided subcortical/cerebellar atlases [Python].

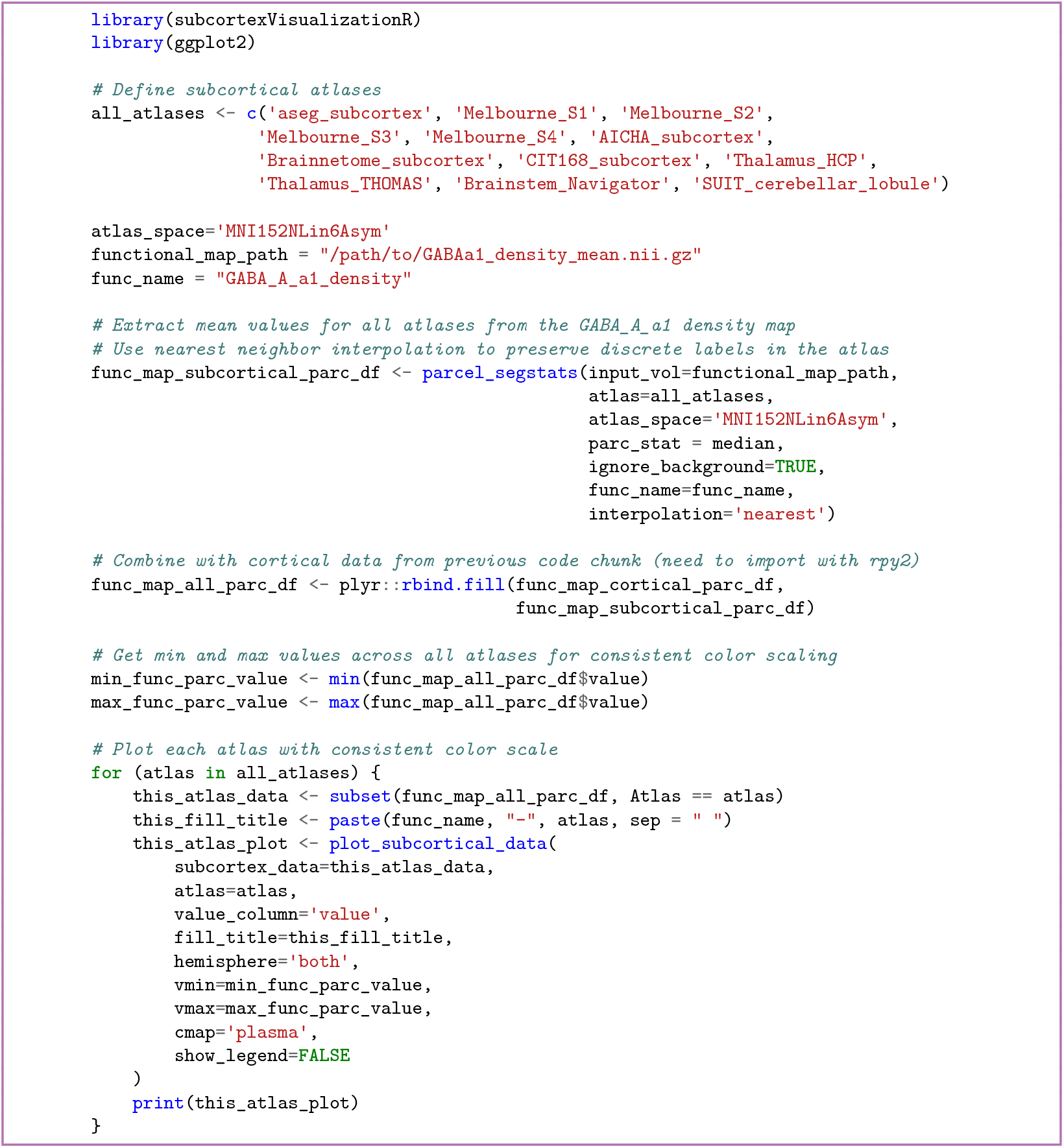

**Code snippet 21**. Computing and plotting the region-averaged GABA_A_-*α*`1` densities in all twelve provided subcortical/cerebellar atlases [R].

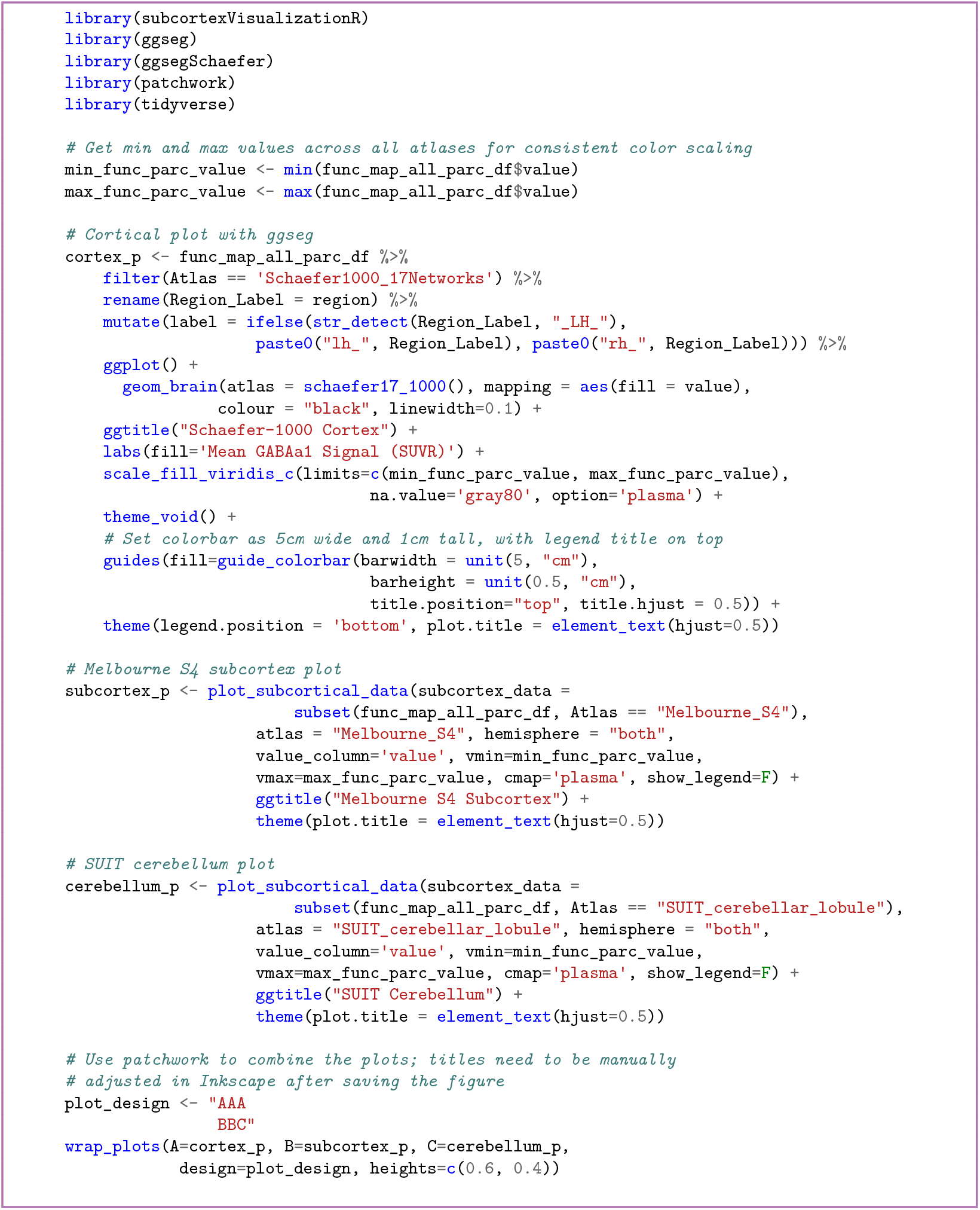

**Code snippet 22**. Plotting GABA_A_-*α*`1` receptor density in the cortex (from Code snippet 19), subcortex, and cerebellum all together [R].

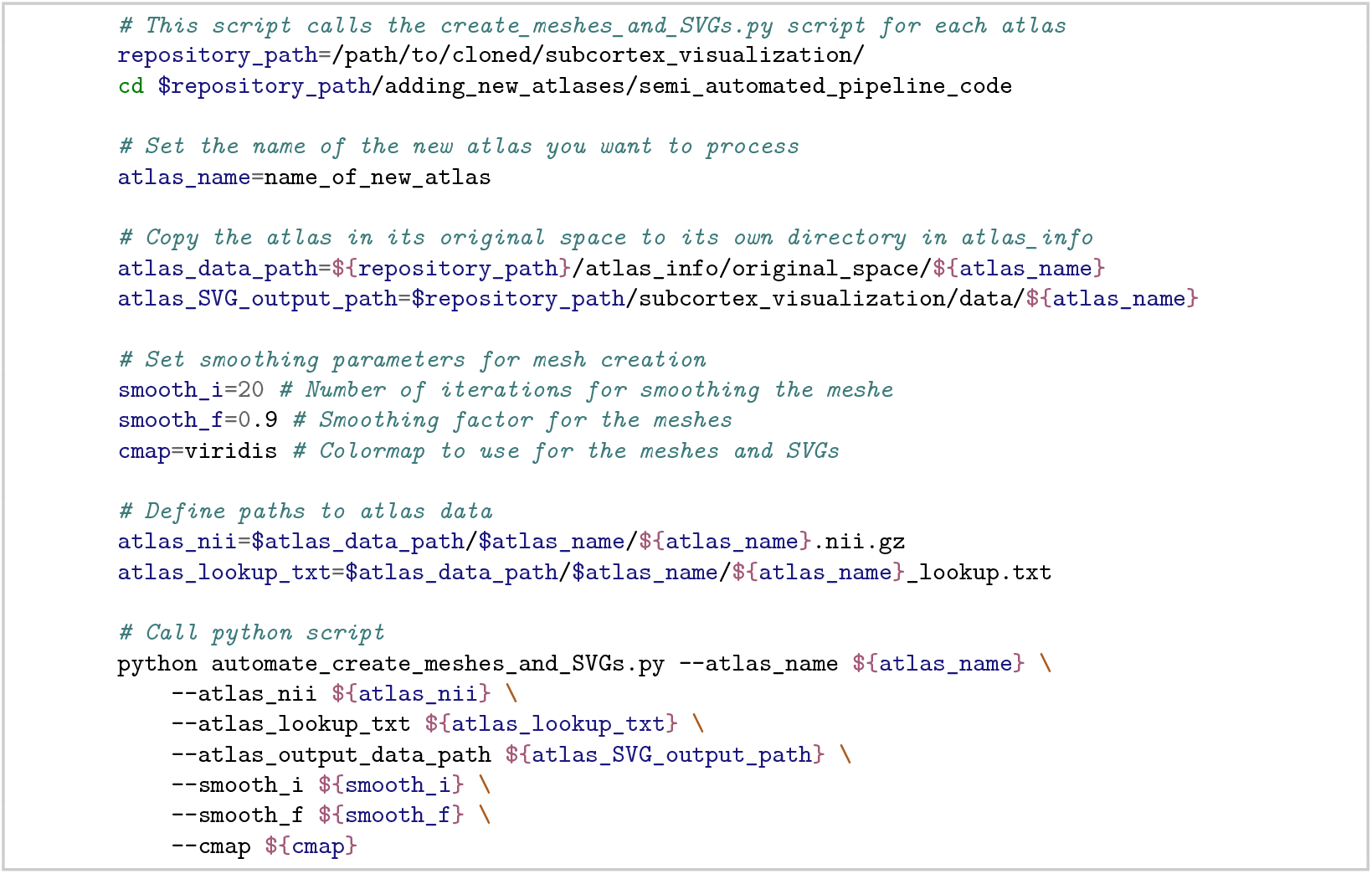

**Code snippet 23**. The semi-automated mesh creation pipeline for a new atlas [bash].

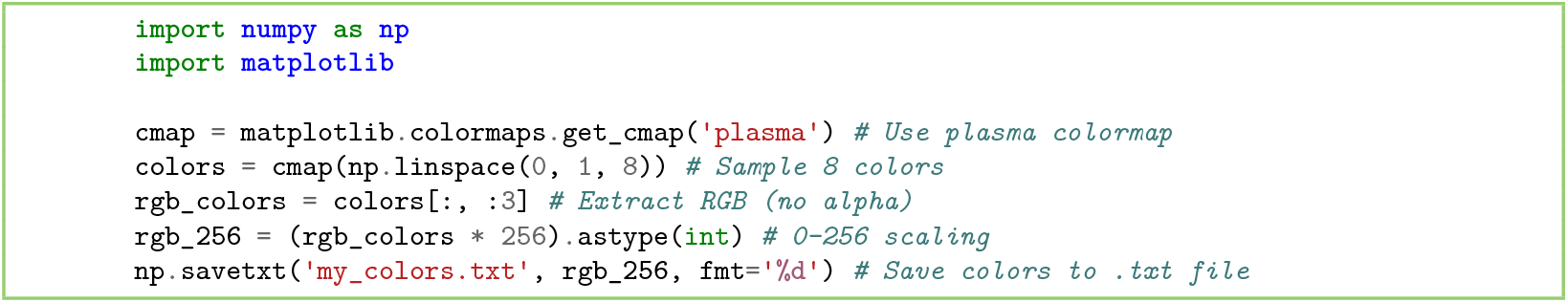

**Code snippet 24**. Generating a custom RGB colormap for the 3D mesh object [Python].

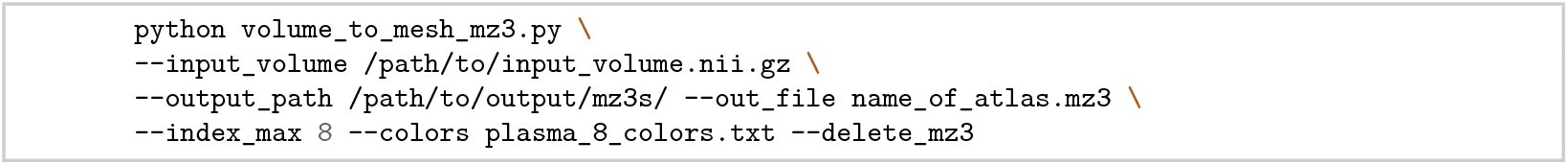

**Code snippet 25**. Rendering the segmentation volume as an .mz3 mesh object for use in Surf Ice [bash].

## Appendix II Supplementary figures

**Figure S1.**
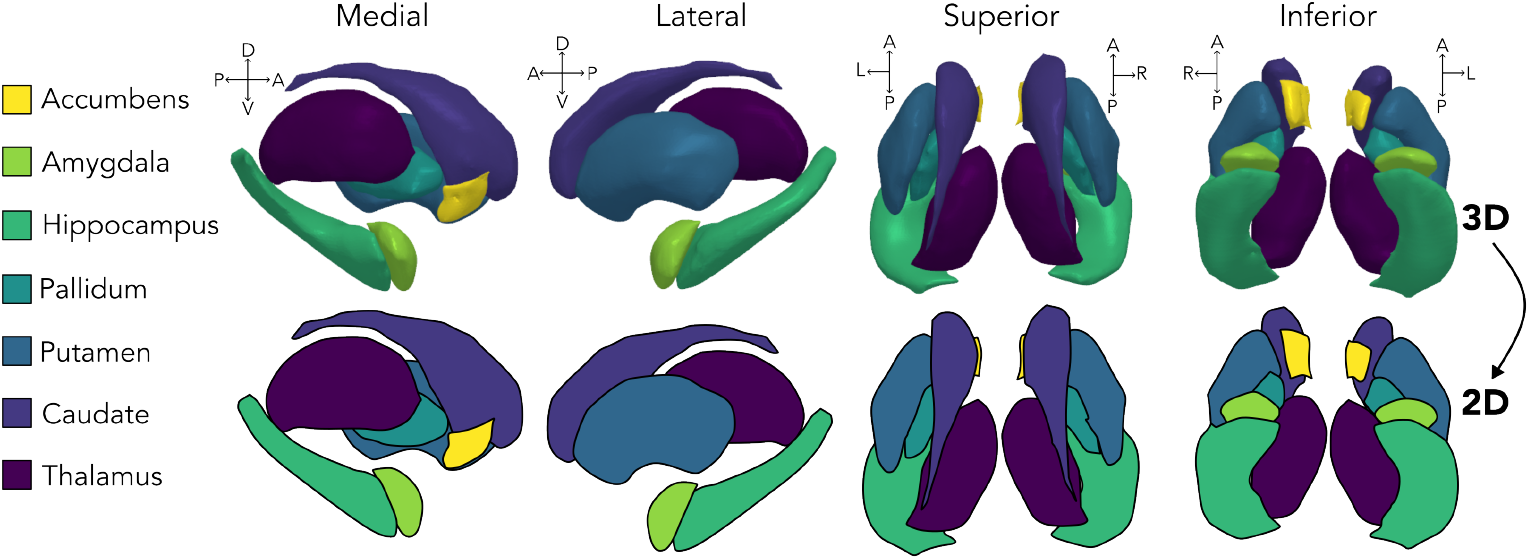
aseg subcortex atlas.

**Figure S2.**
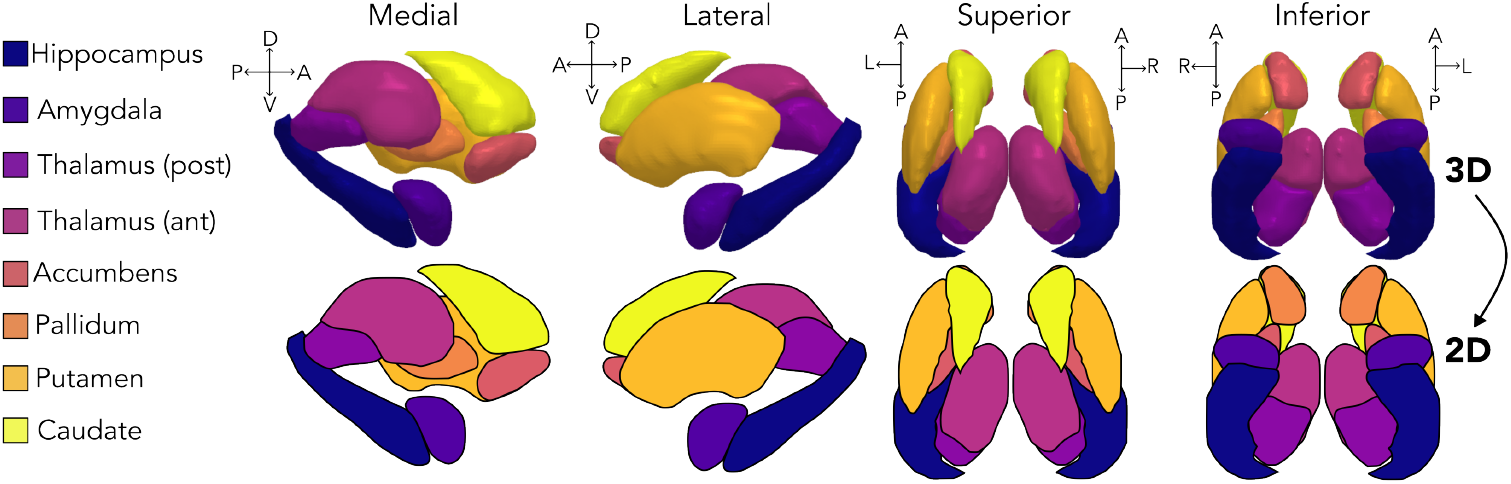
Melbourne (S1) subcortical atlas.

**Figure S3.**
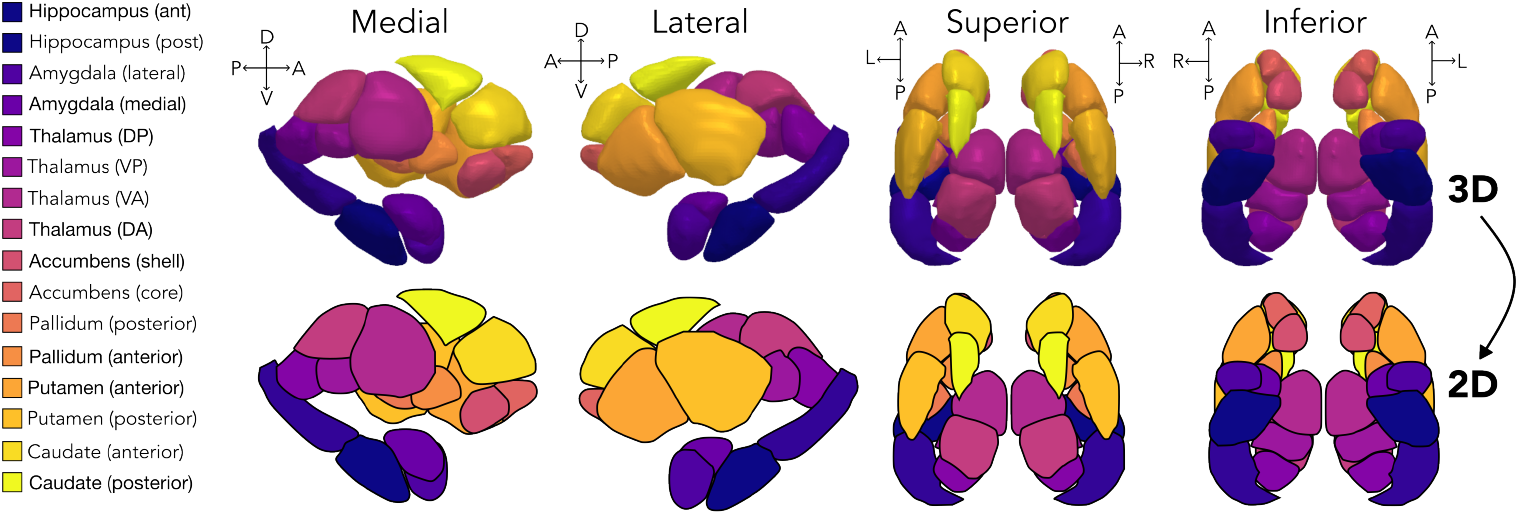
Melbourne (S2) subcortical atlas.

**Figure S4.**
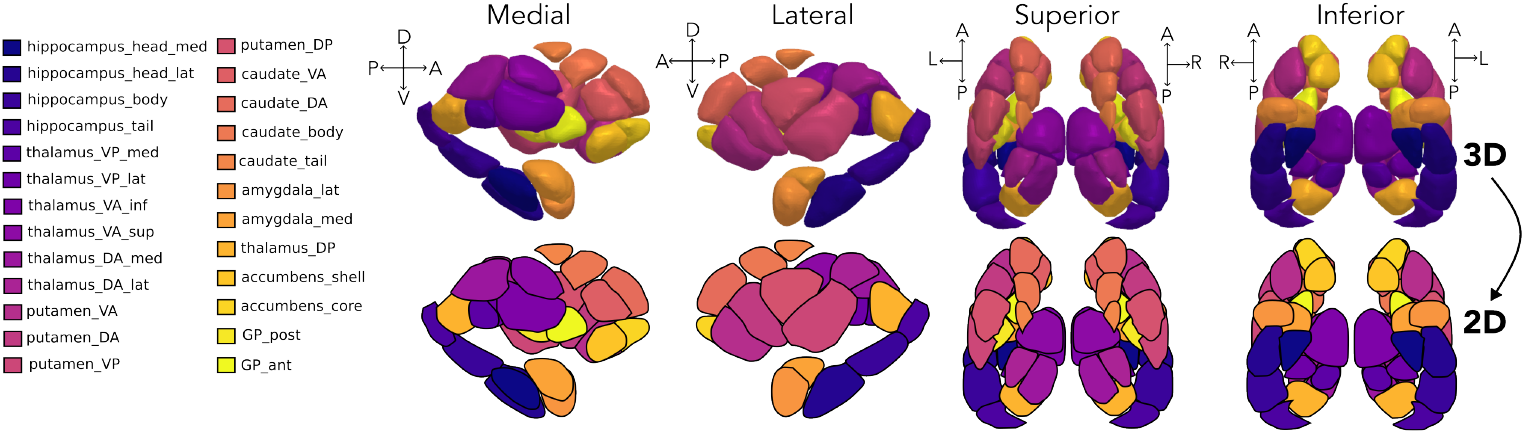
Melbourne (S3) subcortical atlas.

**Figure S5.**
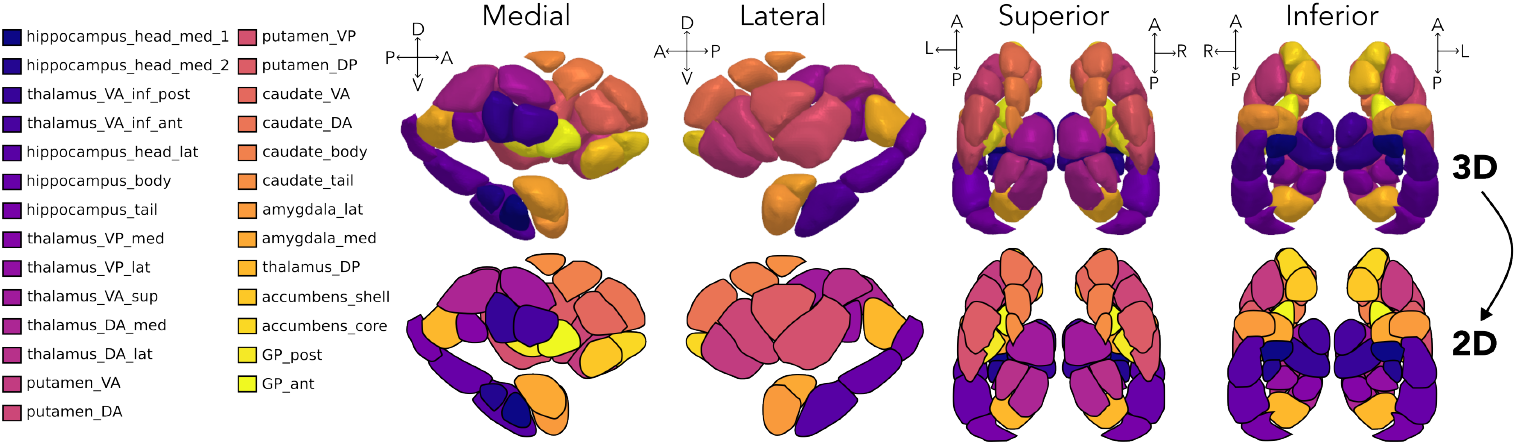
Melbourne (S4) subcortical atlas.

**Figure S6.**
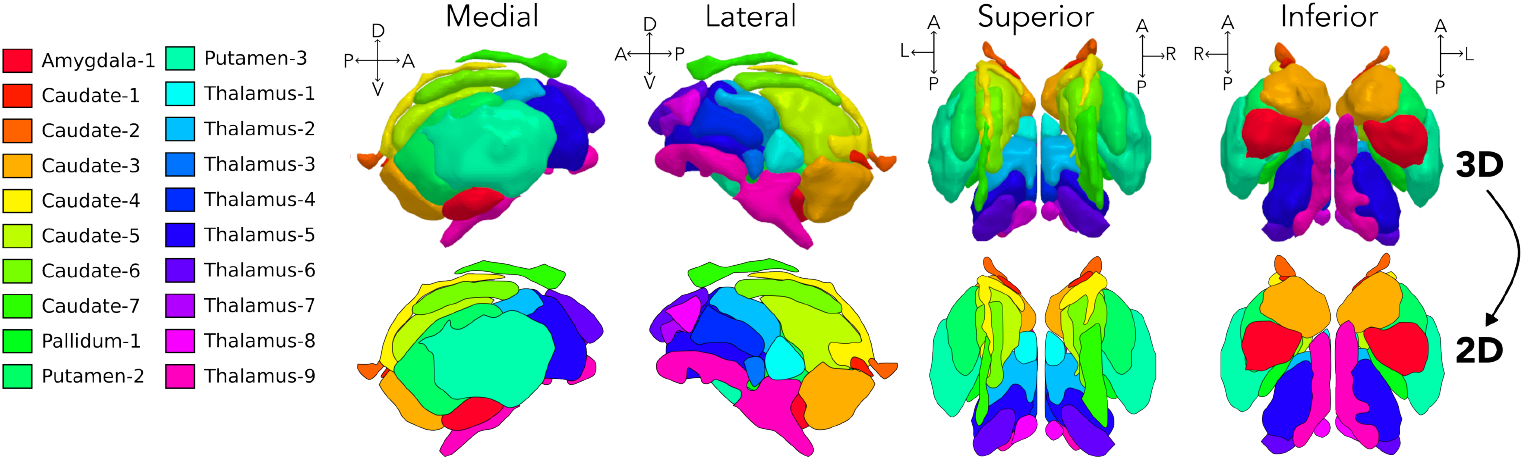
AICHA subcortical atlas.

**Figure S7.**
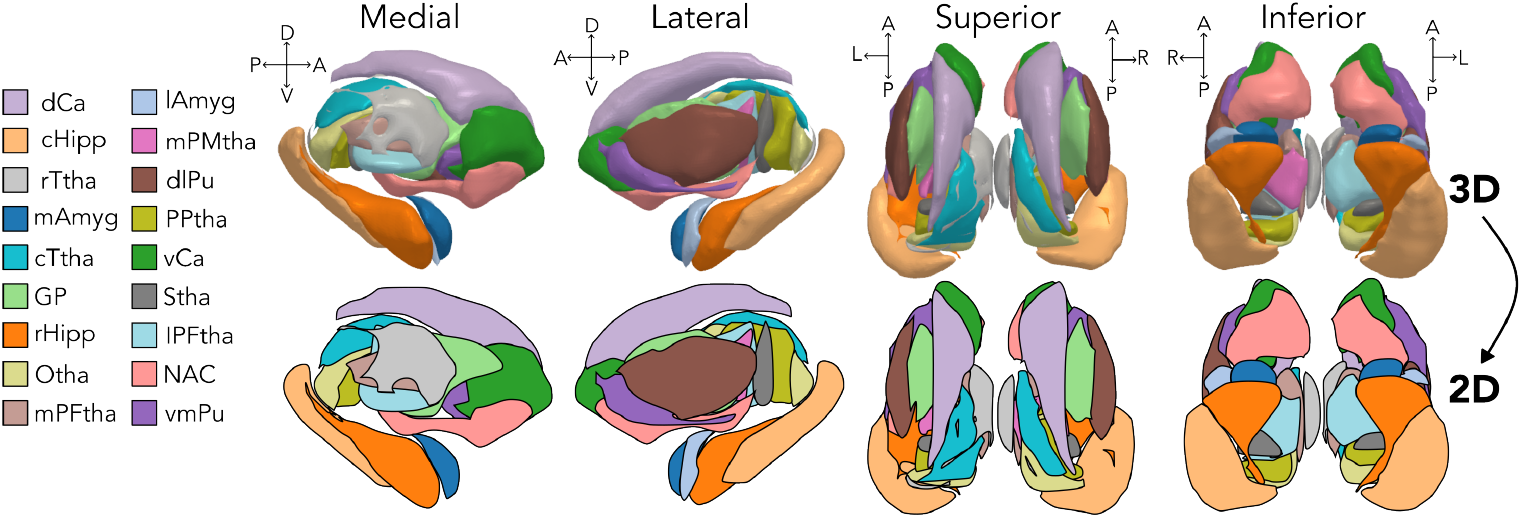
Brainnetome subcortical atlas.

**Figure S8.**
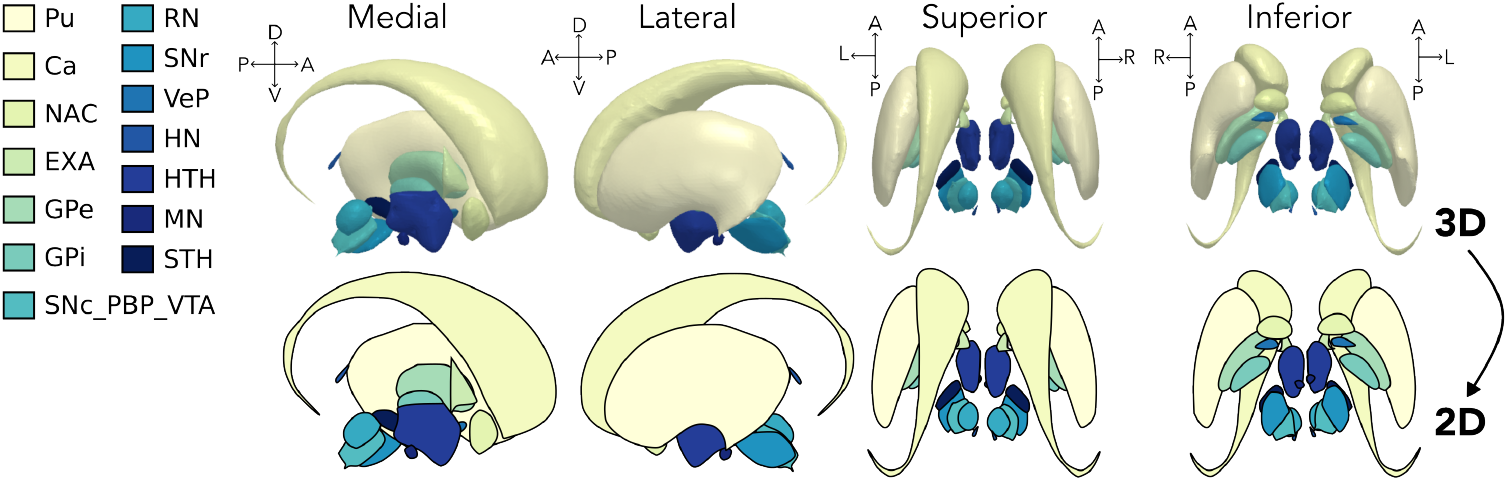
CIT168 subcortical atlas.

**Figure S9.**
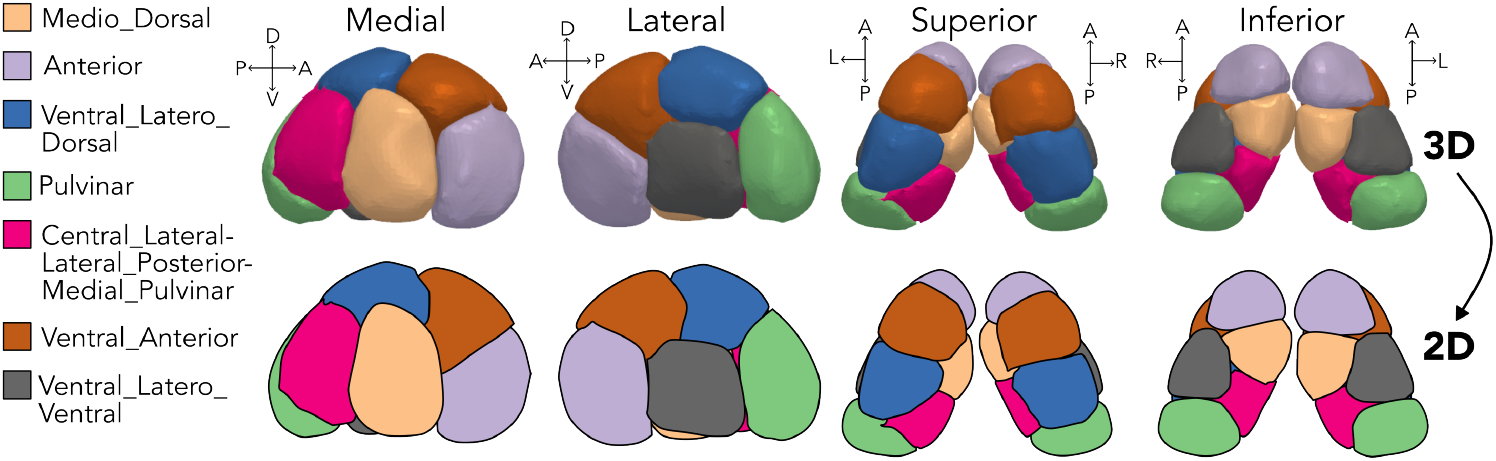
Thalamic nuclei (HCP) atlas.

**Figure S10.**
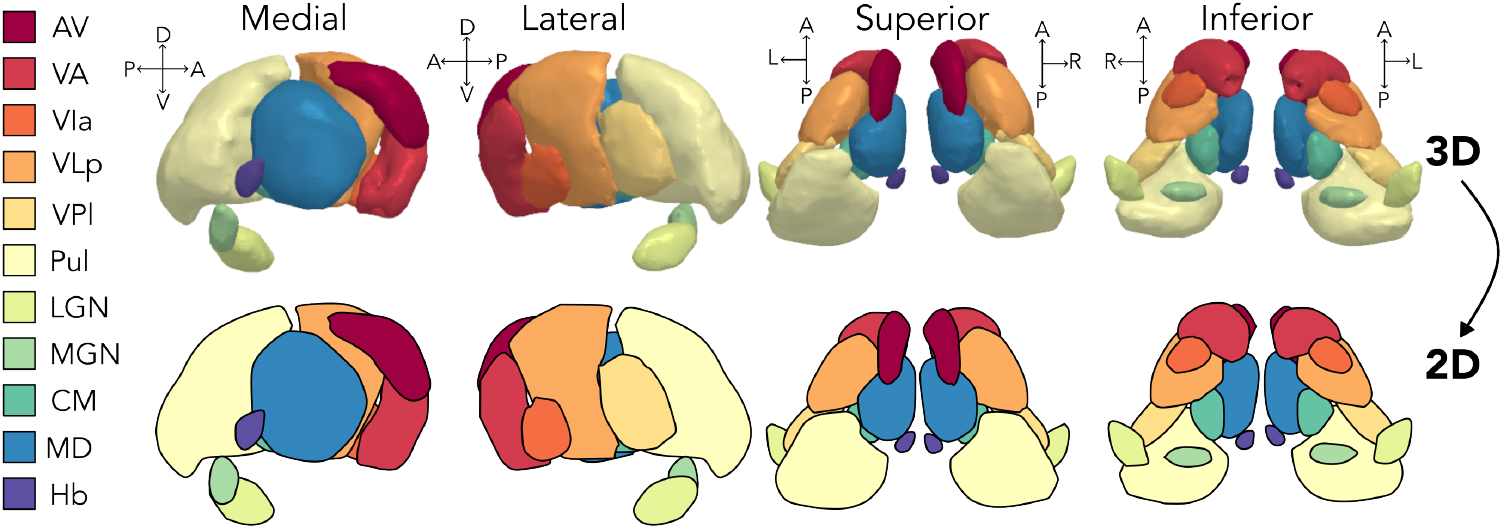
Thalamic nuclei (THOMAS) atlas.

**Figure S11.**
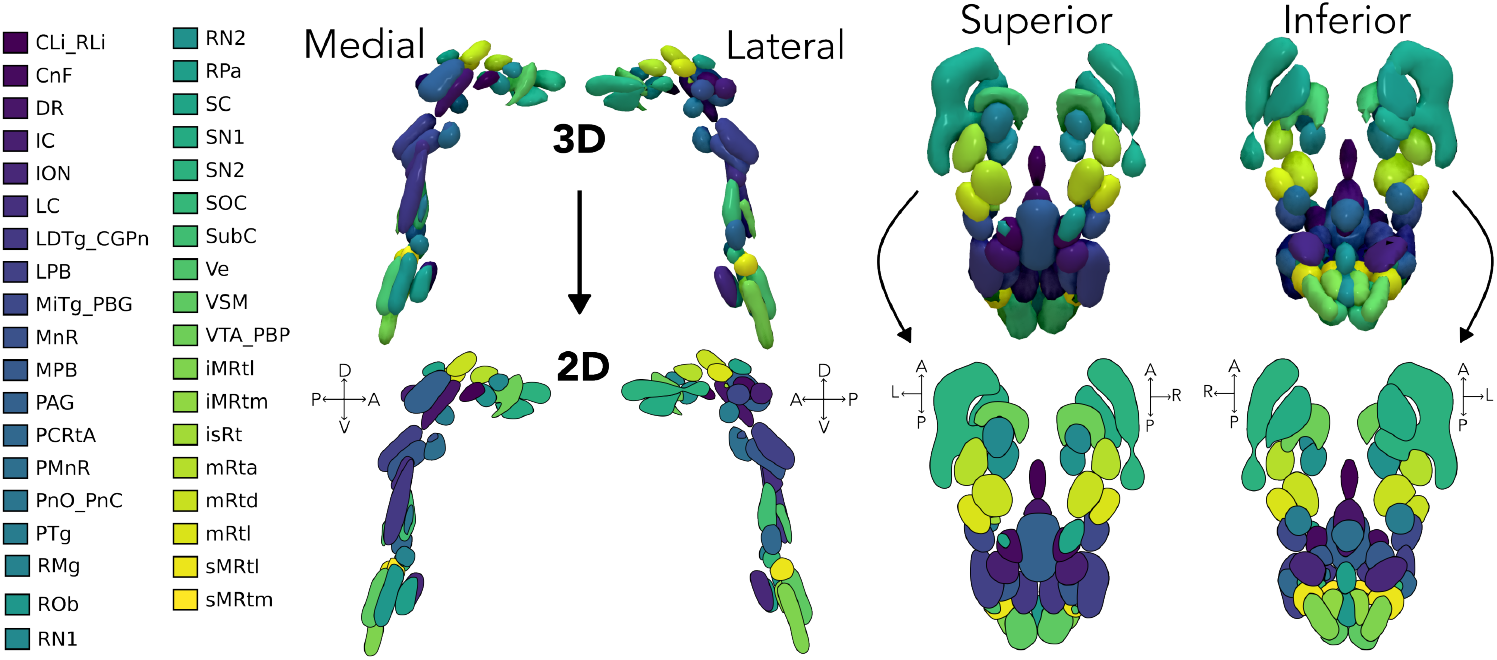
Brainstem Navigator atlas.

**Figure S12.**
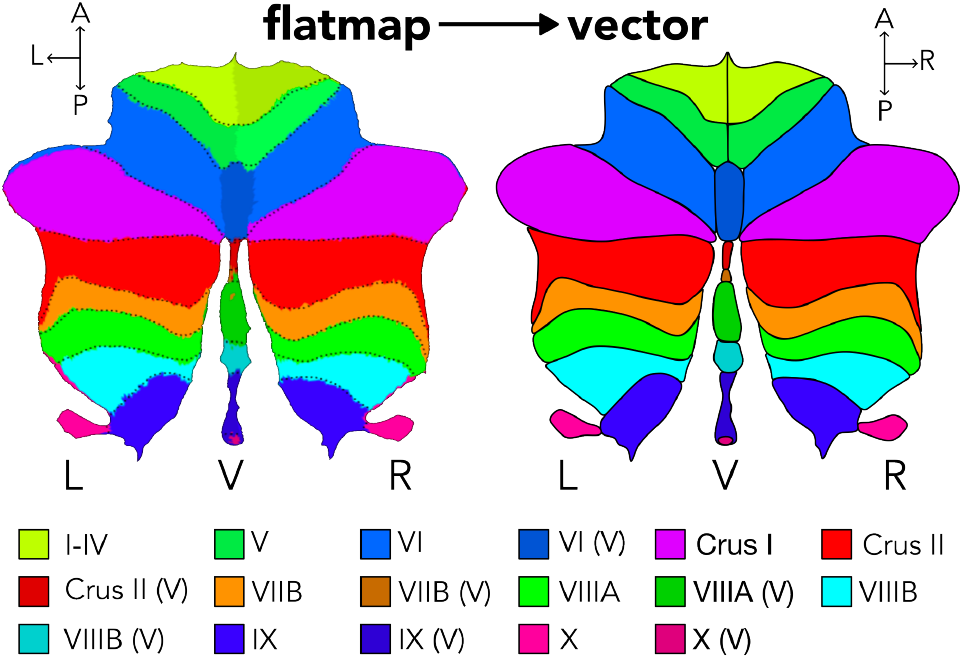
SUIT cerebellar lobular atlas.

## Appendix III Supplementary tables

**Table S1.**
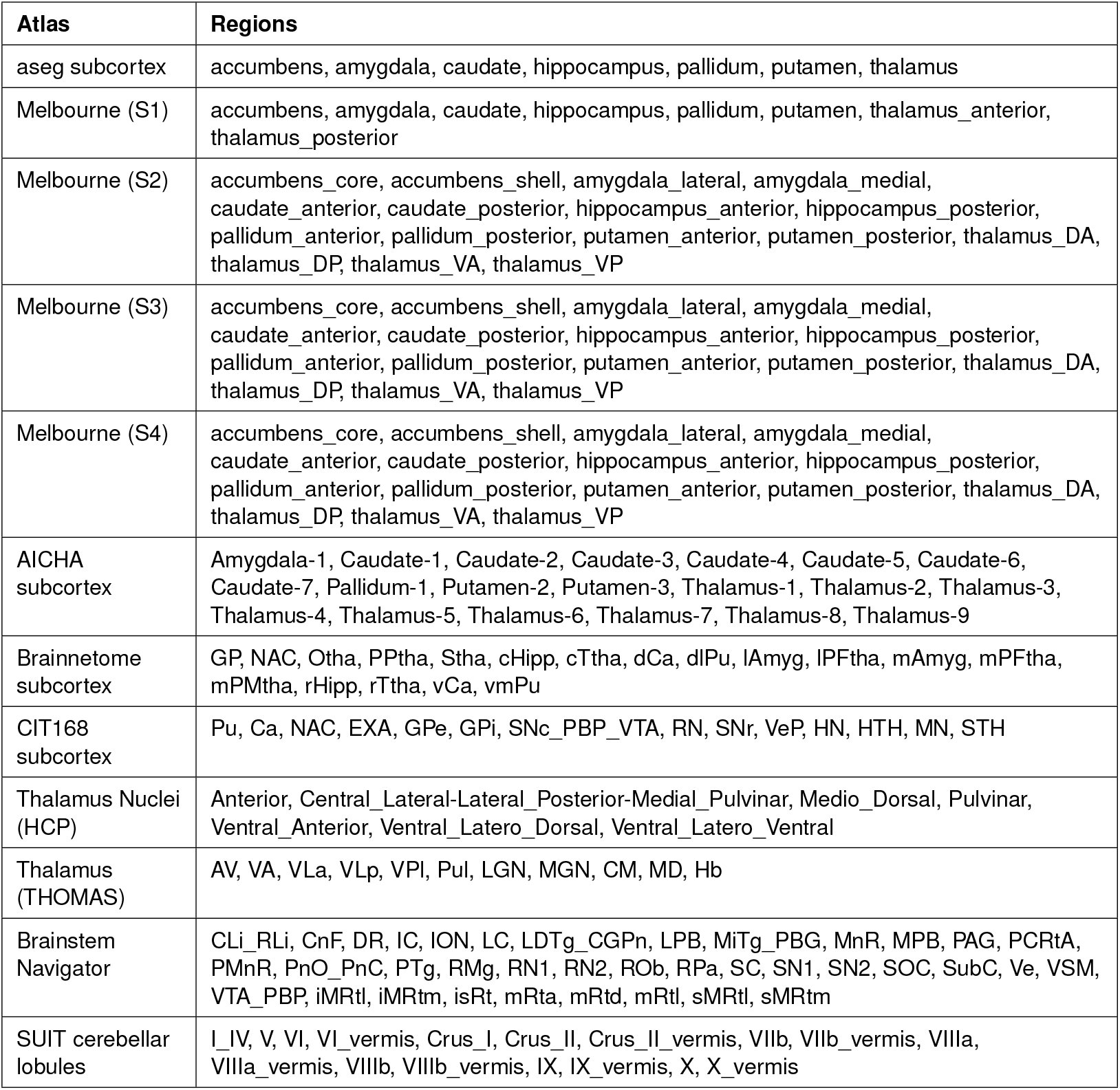
Individual regions within each atlas (per hemisphere).

**Table S2.** Information about original MNI space and download access for each atlas in our toolbox. Links were last accessed on 1 May 2026.

| Atlas | Template space | Download link |
| --- | --- | --- |
| aseg_subcortex | MNI152NLin6Asym | https://github.com/ThomasYeoLab/CBIG/tree/master/data/templates/volume/FSL_MNI152_FS4.5.0/ |
| Melbourne S1 | MNI152NLin6Asym | https://github.com/yetianmed/subcortex/blob/master/Group-Parcellation/3T/Subcortex-Only/Tian_Subcortex_S1_3T_1mm.nii.gz |
| Melbourne S2 | MNI152NLin6Asym | https://github.com/yetianmed/subcortex/blob/master/Group-Parcellation/3T/Subcortex-Only/Tian_Subcortex_S2_3T_1mm.nii.gz |
| Melbourne S3 | MNI152NLin6Asym | https://github.com/yetianmed/subcortex/blob/master/Group-Parcellation/3T/Subcortex-Only/Tian_Subcortex_S3_3T_1mm.nii.gz |
| Melbourne S4 | MNI152NLin6Asym | https://github.com/yetianmed/subcortex/blob/master/Group-Parcellation/3T/Subcortex-Only/Tian_Subcortex_S4_3T_1mm.nii.gz |
| AICHA | MNI152NLin6Asym | https://search.kg.ebrains.eu/instances/Dataset/b820fbed-d920-43c4-9b85-2e6be3d04cd1 |
| Brainnetome | MNI152NLin6Asym | http://www.brainnetome.org/resource/201910/t20191030_522618.html |
| CIT168 | MNI152NLin2009cAsym | https://osf.io/r2hvk/files/osfstorage |
| Thalamus (HCP) | MNI152NLin2009cSym | https://doi.org/10.5281/zenodo.1405484 |
| Thalamus (THOMAS) | MNI152NLin2009bAsym | https://zenodo.org/records/5045684 |
| Brainstem Navigator | MNI152NLin6Asym | https://www.nitrc.org/projects/brainstemnavig |
| SUIT cerebellar lobule | MNI152NLin2009cSym | https://github.com/DiedrichsenLab/cerebellar_atlases/blob/master/Diedrichsen_2009/atl-Anatom_space-MNISym_probseg.nii |

## Appendix IV Supplementary Methods

### 10 Additional atlas information

#### 10.1 aseg subcortex atlas

The aseg atlas was obtained as a volumetric segmentation in MNI152 space from the Computational Brain Imaging Group (CBIG; led by Professor Thomas Yeo), which is based on FreeSurfer’s ‘recon-all’ pipeline applied to the FSL MNI152 1mm isotropic template and was shared in MNI152NLin6Asym space. The results of this pipeline are openly available at https://github.com/ThomasYeoLab/CBIG/tree/master/data/templates/volume/FSL_MNI152_FS4.5.0/. The aseg atlas originally contains both cortical and subcortical regions, so we filtered the atlas to include only the subcortical regions based on the following indices: 10, 11, 12, 13, 17, 18, 26, 49, 50, 51, 52, 53, 54, 58.

#### 10.2 Melbourne Subcortex Atlas

The Melbourne Subcortex Atlas (S1, S2, S3, and S4 resolutions) were obtained from the main repository accompanying the original publication [8]. Specifically, the NIfTI files were downloaded from https://github.com/yetianmed/subcortex/tree/master/Group-Parcellation/3T/Subcortex-Only. For each atlas resolution, we downloaded the 3T volume resampled onto a 1mm isotropic anatomical grid in MNI152NLin6Asym space.

#### 10.3 AICHA subcortical atlas

The AICHA (v2) subcortical atlas NIfTI volume was obtained from EBRAINS at https://search.kg.ebrains.eu/instances/Dataset/b820fbed-d920-43c4-9b85-2e6be3d04cd1 (file name: ‘AICHA1mm.nii’). The AICHA atlas was originally provided in MNI152NLin2009cAsym space. The subcortical indices range from 345-384, which we used to extract the subcortical regions from the full atlas volume.

The full citation for the atlas dataset itself is: Joliot, M., & Naveau, M. (2021). AICHA: An atlas of intrinsic connectivity of homotopic areas (Version 2) [Data set]. EBRAINS. DOI: 10.25493/5NX4-HXR

#### 10.4 Brainnetome subcortical atlas

The Brainnetome atlas (including cortical and subcortical regions) was downloaded from the atlas website at http://www.brainnetome.org/resource/201910/t20191030_522618.html. Specifically, the file downloaded was titled ‘BN_Atlas_246_1mm.nii.gz’. The Brainnetome atlas was originally provided in MNI152NLin6Asym space. The subcortical indices range from 211-246, which we used to extract the subcortical regions from the full atlas volume. This atlas is provided in MNI152NLin2009cSym space.

#### 10.5 CIT168 subcortex atlas

The CIT168 subcortex atlas was obtained from OSF using a shell script prepared by the Penn Lifespan Informatics and Neuroimaging Center (PennLINC) and shared via their GitHub repository at the following link: https://github.com/PennLINC/AtlasPack/blob/main/CIT168/prepare_atlas.sh. The shell script calls a Python script that splits the hemispheres of the original atlas volume such that each region is assigned a unique index across hemispheres (e.g., the left caudate and right caudate are assigned different indices). The CIT168 atlas was originally provided in MNI152NLin2009cAsym space.

#### 10.6 Thalamus nuclei (HCP) atlas

The thalamic nuclei atlas based on HCP data was obtained from Zenodo at https://zenodo.org/record/1405484. This atlas was originally provided in MNI152NLin2009cSym space. The full citation for the atlas dataset is provided below.

Najdenovska, E., Alemán-Gómez, Y., & Bach Cuadra, M. (2018). Dataset In-vivo probabilistic atlas of human thalamic nuclei based on diffusion weighted magnetic resonance imaging (1.0) [Data set]. Zenodo. https://doi.org/10.5281/zenodo.1405484

#### 10.7 Thalamus nuclei (THOMAS) atlas

The THOMAS thalamic nuclei atlas was obtained from the Zenodo repository that accompanies the original publication at https://zenodo.org/records/5045684. Please note that the Zenodo record is publicly available but the files are restricted and require users to request access to download the files directly. The THOMAS atlas was originally provided in MNI152NLin2009bAsym space.

#### 10.8 Brainstem Navigator atlas

The Brainstem Navigator atlas was obtained as a directory of individual NIfTI files for each brainstem region from NITRC at https://www.nitrc.org/projects/brainstemnavigator. We selected the region of interest (ROI) files contained in the ‘2a.BrainstemNucleiAtlas_MNI/labels_thresholded_binary_0.35/’ directory to use binary segmentation volumes of each brainstem nucleus. The Brainstem Navigator atlas regions were all shared in MNI152NLin6Asym space.

#### 10.9 SUIT cerebellar lobule atlas

Our implementation of the SUIT cerebellar lobule atlas is based on the surface flatmap rendering shared by Diedrichsen and Zotow [54]. However, we also include a volumetric segmentation of the SUIT cerebellar lobules in our toolbox, which was obtained from the Diedrichsen Lab GitHub repository at the following link: https://github.com/DiedrichsenLab/cerebellar_atlases. The SUIT volumetric atlas was originally provided in MNI152NLin2009cSym space.

### 11 Further technical details about atlas alignment to MNI152NLin6Asym and MNI152NLin2009cAsym spaces

As shown in Table S2, the twelve atlases included in our toolbox are originally provided in either MNI152NLin6Asym, MNI152NLin2009cAsym, MNI152NLin2009cSym, or MNI152NLin2009bAsym space. Since the MNI152NLin6Asym and MNI152NLin2009cAsym spaces are the most commonly used template spaces in our toolbox, we aligned the atlases that were originally provided in other spaces to both of these template spaces. We opted to use the widely adopted `templateflow` framework to perform the atlas alignments, which provides pre-computed transformations between commonly used template spaces. For template pairs that did not already have pre-computed transformations available in TemplateFlow (e.g., MNI152NLin2009bAsym to MNI152NLin2009cAsym), we performed registration using the `registration` function from the ANTS toolbox in Python [81], using the `type_of_transform=`‘`SyN`’ function to specify symmetric normalization. In either case, we applied the corresponding transform file to each atlas as needed using the `apply_transforms` function from ANTS, specifying nearest-neighbor interpolation with the `interpolator=`‘`nearestNeighbor`’ to preserve discrete region indices.

First, we provide an example of the Python code we used to transform an atlas from MNI152NLin2009cAsym to MNI152NLin6Asym space, for which TemplateFlow provides a pre-computed transformation. The code snippet is shown in Listing 26.

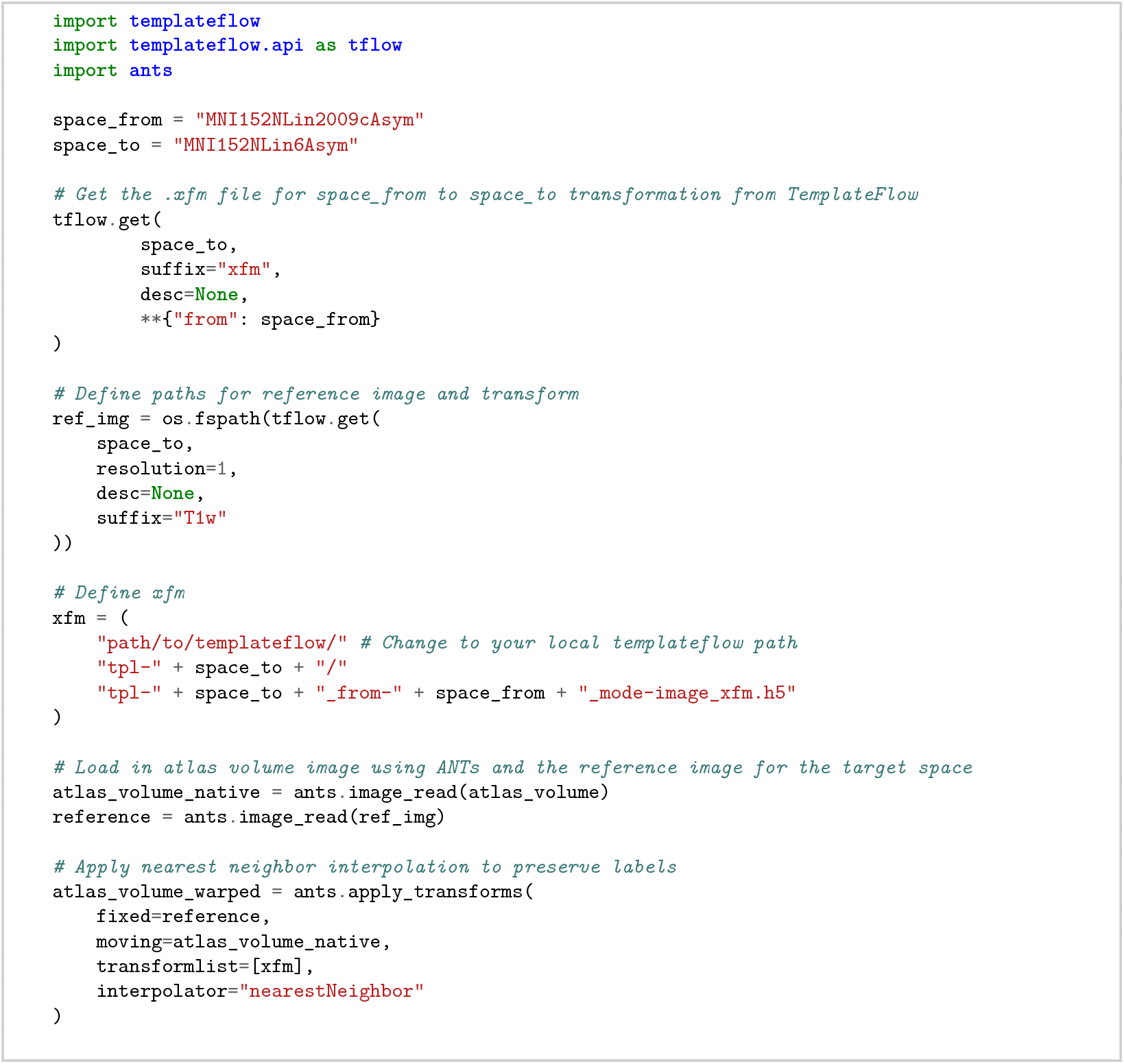

**Code snippet 26**. Example of how to align a segmentation atlas from MNI152NLin2009cAsym to MNI152NLin6Asym space [Python].

Alternatively, we show an example of how to perform registration using ANTS when a pre-computed transformation is not available in TemplateFlow. In this example, we align an atlas from MNI152NLin2009bAsym to MNI152NLin2009cAsym space. The code snippet is shown in Listing 27.

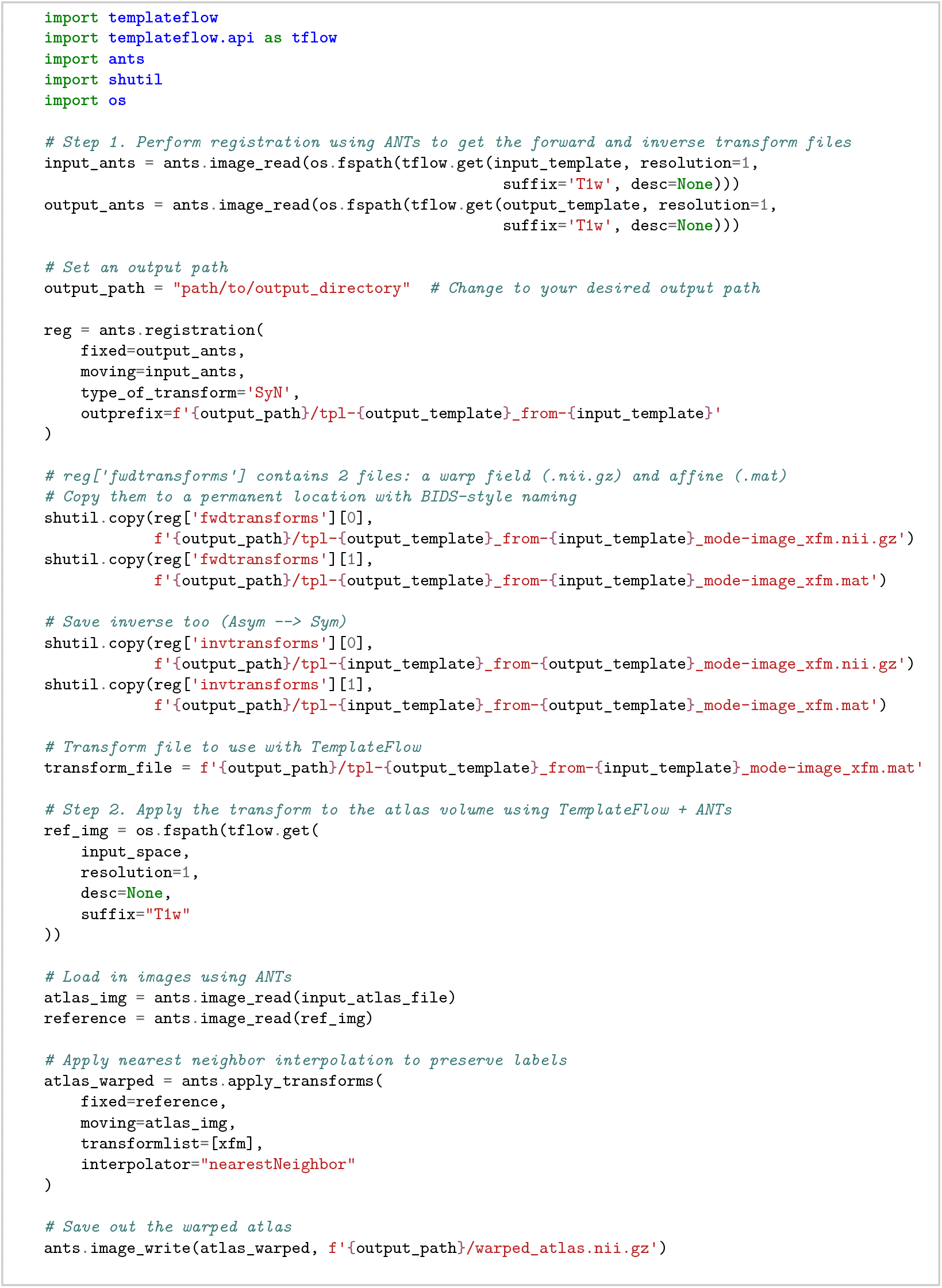

**Code snippet 27**. Example of how to align a segmentation atlas from MNI152NLin2009bAsym to MNI152NLin2009cAsym space [Python].

### 12 Further information about mesh-to-vector conversion

For the semi-automated mesh-to-vector conversion process, we used the following smoothing parameters with the `yabplot.build_subcortical_atlas` function:

**Table S3.**
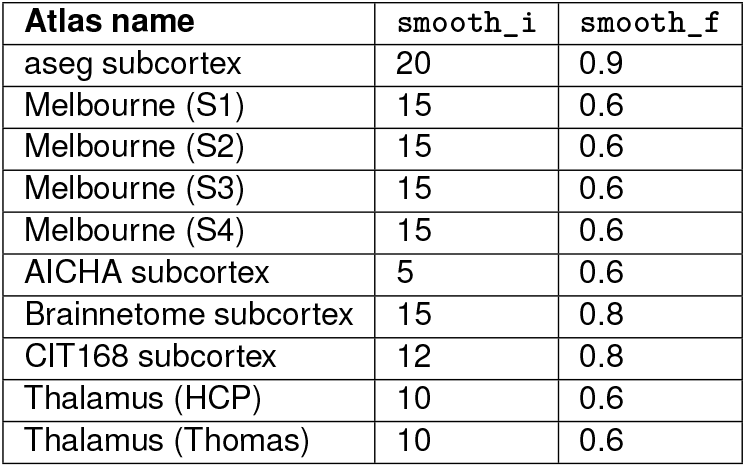
Mesh smoother parameters supplied to `yabplot.build_subcortical_atlas` for each atlas are shown here. The `smooth_i` parameter specifies the number of iterations of smoothing to perform, while the `smooth_f` parameter specifies the smoothing factor (between 0 and 1) that controls the amount of smoothing applied at each iteration.

## Footnotes

1 https://github.com/ggseg/python-ggseg/

2 https://anniegbryant.github.io/subcortex_visualization/

3 https://anniegbryant.github.io/subcortex_visualization/installation/

4 https://anniegbryant.github.io/subcortex_visualization/atlas_info/

5 https://anniegbryant.github.io/subcortex_visualization/custom_segmentation/

6 https://anniegbryant.github.io/subcortex_visualization/custom_segmentation/

## Notes

### Competing Interest Statement

The authors have declared no competing interest.

### Summary of Updates

Section 3 updated to include an alternative semi-automated pipeline for adding vectorized versions of new atlases; expanded functionality for statistical computation and additional view angles described; three new atlases added and documented.

